# Unveiling the Mystery of the Petroleum Genome Archive: Paleofauna Activities, Geological History, and aDNA Identification Methods in Nanyang Oilfield

**DOI:** 10.1101/2024.12.24.630223

**Authors:** Wan-Qian Zhao, Zhan-Yong Guo, Yu-Qi Guo, Mei-Jun Li, Gang-Qiang Cao, Zeng-Yuan Tian, Ran Chai, Li-You Qiu, Jin-Hua Zeng, Xin-Ge Zhang, Tian-Cang Qin, Jin-Yu Yang, Ming-Jie Chen, Mei-Rong Song, Fei Liang, Jun-Hui Geng, Chun-Yan Zhou, Shu-Jie Zhang, Li-Juan Zhao

## Abstract

Specific macromolecules in petroleum, preserved through geological processes from ancient organisms, are recognized as molecular fossils or geochemical markers (GMs). Our selected GMs indicate that the source rocks of Nanyang Oilfield formed in an anoxic, saline, and stratified lacustrine environment with marine influence. Contrary to science fiction notions that oil originates from ancient large animals, mainstream academia has consistently asserted that oil is derived from the organic matter of lower life forms, formed over extended geological processes, with intact DNA no longer persisting in oil post-formation. However, this study successfully extracted 3,159,020 DNA fragments from the petroleum samples using the nanoparticle affinity bead technology, followed by sequencing and classification via the mega screen method. These fragments, categorized as original *in situ* DNA (oriDNA), recent environmental DNA (preDNA), and paleo-environmental DNA (paeDNA), exhibit a “Genome Accumulated over Time” (GAT) pattern of “old-less, new-many”. This aligns with findings from GMs while offering more specific genetic information. The analysis reveals evidence of historical events, including animal extinctions, marine invasions, ancient bird species presence, Asian *Homo erectus* activities, and modern human influences in preDNA. Notably, the mitogenomic data positions Asian *Homo erectus* closer to modern humans than to chimpanzees at the maternal basis. The absence of deamination in petroleum ancient DNA (aDNA) fragments is likely due to the anaerobic, anhydrous environment and the screening process that excludes degraded fragments. Consequently, the authenticity criteria and scope of application of traditional aDNA methods require reevaluation. Surprisingly, the near-absence of oriDNA has impeded our efforts to determine the specific biological origins of the oil. These findings position petroleum as a novel, non-skeletal source of aDNA, offering insights into ancient animal activities, geological changes, and human origins while underscoring its potential as an untapped resource for studying genetic preservation across geological timescales.

## Introduction

Petroleum, a fossil fuel, is widely recognized as originating from fossilized organic matter, such as plankton and bacteria, which accumulated alongside sediments in ancient oceans or lakes and transformed into petroleum under thermal pressure ^1,2^. GMs, also termed molecular fossils or biomarkers, are organic compounds preserved in geological samples like sediments, rocks, crude oils, and coals. These molecules, exhibiting minimal structural alteration from their biological precursors in living organisms, offer valuable insights into past biological and environmental conditions. As such, they serve as tools for addressing challenges in exploration, development, production, environmental studies, and archeological or paleontological research ^3^. Prior studies indicate that the primary source rocks of the Biyang Sag oil field in the Nanxiang Basin lie within the third member (Eh_3_) of the Paleogene Hetaoyuan Formation ^4,5,6,7,8^. The organic matter in these Eh_3_ shales predominantly derives from phytoplankton, bacteria, and benthic algae, with minor contributions from animal and higher plant material. Furthermore, the presence of 24-n-propylcholestanes, indicative of marine organisms, suggests episodes of marine incursion in the Nanxiang Basin ^6,7,8^.

Paleontologists have claimed the ability to extract oriDNA from certain fossils dating back 1 million years (Ma) ^9^. However, fossil distributions are often sparse, geographically scattered, and temporally fragmented. Expanding the range of aDNA sources could significantly advance paleontological research. Experimental evidence has demonstrated that vegetable oil refined at 400°C retains amplifiable DNA fragments ^10,11,12,13^, suggesting that small quantities of DNA may persist in the deoxygenated and dehydrated environment. In contrast, petroleum formation involves the maturation of organic matter in source rocks, with hydrocarbon generation typically occurring at temperatures between 60°C and 180°C ^1^. This questions whether the oil DNA, including the oriDNA and environmental DNA (eDNA) from surrounding contexts, could be extracted and amplified. Such a success could provide critical insights into the genetic information preserved during petroleum’s formation, migration, accumulation, and long-term storage.

Employing nanoparticle affinity bead technology, we successfully extracted DNA from petroleum sourced from the Nanyang Oilfield, constructed DNA libraries, and recovered 3,159,020 DNA fragments. Using the mega screen method, we classified the petroleum DNA (pDNA) according to their origins, distinguishing preDNA from paeDNA. The paeDNA proved instrumental in validating key local historical events, including marine invasions, activities of unclassified *Homo species*, and the presence of ancient bird species. Contrary to expectations, oriDNA from the petroleum was virtually undetectable, hindering attempts to determine the specific species contributing to the oil formation. Nonetheless, this discovery positions petroleum as an innovative reservoir for retrieving aDNA fragments from non-skeletal materials, providing a valuable avenue for investigating the evolutionary trajectory of ancient regional life within the context of local paleontology and geological development. Notably, most petroleum aDNA fragments exhibited “non-deaminated” characteristics, challenging the scope and validity of Pääbo’s team’s reliance on deamination as a primary indicator of aDNA authenticity. This discovery underscores the need for a global pDNA database, which could serve as a transformative platform for interdisciplinary research spanning geology, industrial genomics, and paleontology.

## Results and Discussion

### I. Petroleum Geology of the Nanyang area

During the Early to Middle Triassic, the Nanyang Oilfield was situated within a trough in the Qinling region of southern China. This area subsequently consolidated into a cohesive landmass following the collision of the Sino-Indian Plate in the Middle to Late Triassic ^14^. The oil wells are situated in the Biyang Depression between the North China and the Yangtze Plates. The source rocks of these studied crude oil samples were deposited under a high-salinity and strongly reduced environment with a stratified water column. Geochemical studies on Eh_3_ shales, the most favorable hydrocarbon source rock in the oil fields, have revealed an average organic matter abundance of 2.96 % (wt%), indicating low to mature stages of maturity, with prevalent type I to type II kerogen dominance ^15^. The organic matter in source rocks primarily consists of bacteria, algae, and a small amount of higher plants from land.

The reservoirs are buried at depths of 90 to 1000 m, with oil layer thicknesses ranging from 1 to 4 m and assemblage thicknesses ranging from 2 to 15 m. The petroleum reservoirs in the Nanyang Oilfield experienced a complex filling history, including an early oil filling, significant uplift, a second phase of filling, and late-stage biodegradation. Therefore, the oils with a burial depth of less than 1100 m are characterized by higher density, higher viscosity, and depletion of light hydrocarbon fraction ^16,17,18^. The significant uplift of stratum and erosion made the reservoired oil susceptible to incorporation into the earth’s surface organisms. Fossil evidence suggests that the region was inhabited by Asian *Homo erectus* and a variety of mammals during the Middle Pleistocene dating back 0.5 Ma ^19^. This region has been densely populated since the arrival of modern agricultural civilization.

### II. Geochemical Markers as Indicators of Biological Origins

GMs exhibit limited ability to distinguish specific species contributing to organic inputs due to frequent overlap in their distributions across different species. However, certain GMs serve as reliable indicators of specific biological sources. In this study, β-carotene is associated with cyanobacteria or algae, while 3β-methylhopanes denote methanotrophic bacteria. Conversely, gammacerane, present in oil and sedimentary organic matter, derives from tetrahymanol produced by ciliates that feed on bacteria. Low pristane/phytane ratios and elevated gammacerane abundance typically correlate with anoxic, saline, and stratified water bodies ^3^. Similarly, a reduced pristane-to-phytane ratio paired with a higher gammacerane index suggests a predominant contribution from aquatic organisms, with minimal input from higher plants, in a saline lacustrine basin. The prominence of tricyclic terpanes (TT) with the high frequency of C_23_TT points to a marine or saline lacustrine depositional environment ^20^. Our GM findings posit that most petroleum molecules originate from lower plants and plankton, a perspective aligned with the prevailing scholarly views, as depicted in Figures S1 and S2.

### III. Extraction of DNA and Analysis via the Mega Screen Method

Using the nanoparticle affinity bead technology, the pDNA was extracted and sequenced, the 3,159,020 pDNA sequences were categorized for further analysis via the mega screen method (Materials and Methods). These sequences, derived from diverse species and sources, were broadly classified into oriDNA, paeDNA, and preDNA.

#### 1. Establishing lineage subsets for pDNA fragments

Non-discriminatory alignment was performed without predefined target genomes using the Best E-value mode (BE mode), aligning over 3.15 million total sequences (TS) with the NCBI database (Materials and Methods). These sequences were subsequently categorized into lineage subsets: 51.29% (1,620,178 fragments) were assigned to bacteria, 33.16% (1,047,532 fragments) to the human genome, 3.85% (121,735 fragments) to the fungal genome, and 0.29% (9,252 fragments) to the avian genome. Additionally, 0.15% (4,785 fragments) corresponded to non-primate mammals and 1.11% (35,108 fragments) to non-human primates (MHPs). A minimal fraction (5 fragments) was attributed to algae, excluding red or brown algae (Figure 1). Notably, these DNA-based findings differ from the above GM studies, as the species distribution and frequency indicated by DNA contrast with those inferred from GMs.

**Figure 1.**
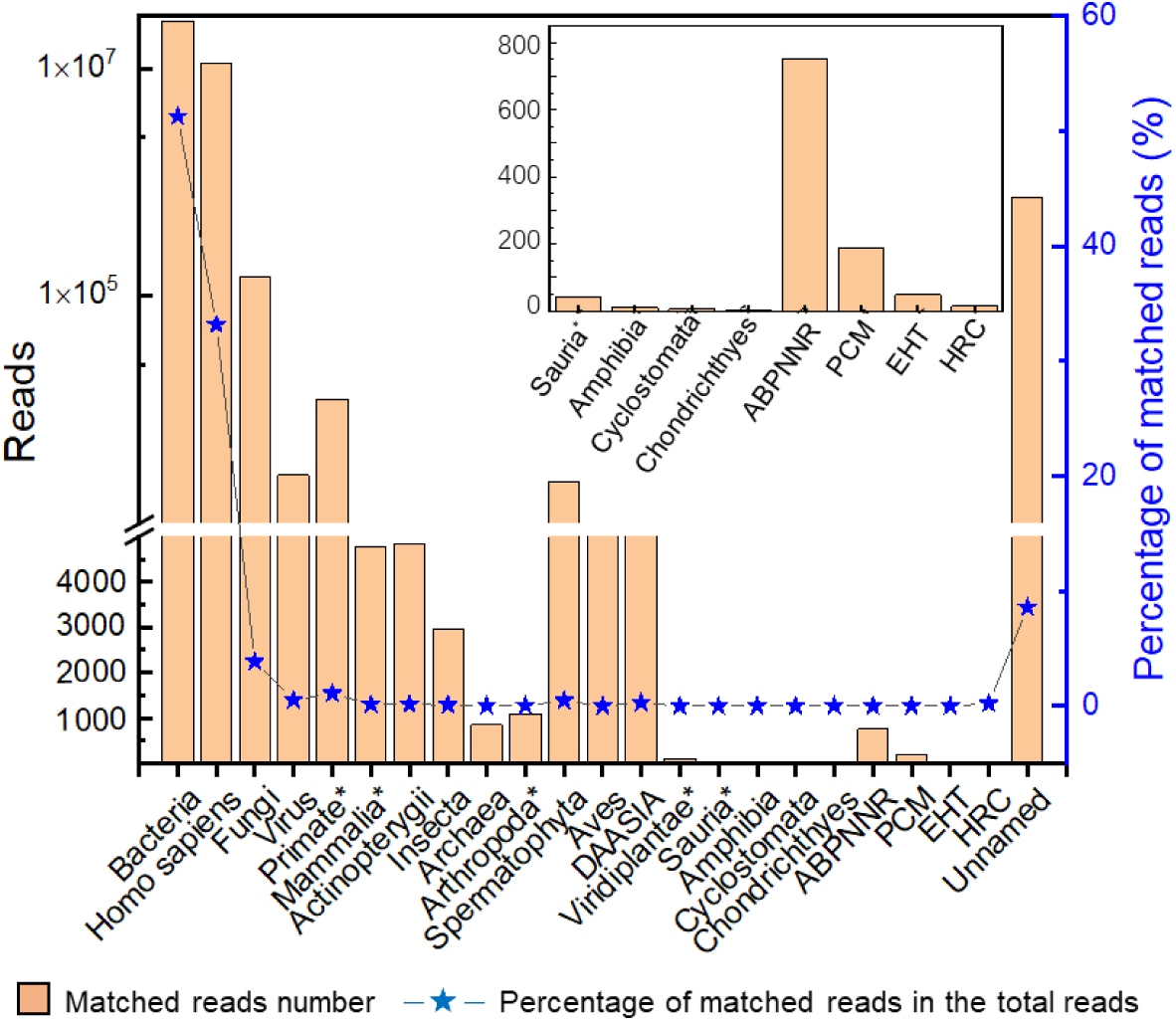
Analysis of pDNA sequences corresponding to different lineages The bars represent the number of pDNA sequences that match a lineage, while the lines represent the percentage of pDNA fragments in that lineage compared to the total number of pDNA fragments. The abbreviations used are as follows: ABPNNR (Rotifera, Nemertea, Nematoda, Platyhelminthes, Annelida, and Bryozoa), PCM (Porifera, Cnidaria, and Mollusca), EHT (Hemichordata and Tunicata), HRC (Haptophyta, Rhodophyta, Cryptophyceae), and DAASIA (Discoba, Apusozoa, Amoebozoa, Sar, Ichthyosporea, and Apicomplexa). It’s important to note that Viridiplantae* excludes Spermatophyta, Arthropoda* excludes Insecta, Sauria* excludes Aves, Primates* excludes *Homo*, and Mammalia* excludes Primates*. Additional details are provided in Tables S1 and S11.

#### 2. Identify the unique lineage source of each sequence

We employed the MS mode (Materials and Methods) to determine the specific species fragments (SSFs) and their lineage origin. Through this approach, at least 400 pDNA sequences have been successfully identified as SSFs and categorized into nine groups based on their morphological characteristics and habitat (Table 1), including lower plants (Group G), marine organisms (Group M), primates (Groups C, P, and T), and birds (Group L).Although these sequences cannot trace major oil-forming species, they provide valuable data reflecting local geology, history, and evolution, an unexplored reservoir for investigating local geological and species evolution. Due to the lack of genome data for certain lower species, some pDNA sequences could not be genealogically determined.

**Table 1.** Summary of the SSFs in pDNA sequences matched genomes.

| Group | Target | ID | Lineages | pDNA Count | Affinity |  |
| --- | --- | --- | --- | --- | --- | --- |
|  |  |  |  |  | Average | SD |
| G | Plastid + Nuclear DNA | 1G-8G | *Chlorophyta, Streptophyta* | 8 | 90.23% | 16.05% |
| H | Primate mtDNA | 1H-40H | *Homo sapiens* | 40 | 96.12% | 15.44% |
| N | Non-primate mtDNA | 1N | *Gallus gallus, Galliformes* | 1 | 99.32% | - |
|  |  | 2N | *Sus scrofa, Artiodactyla* | 1 | 99.48% | - |
|  |  | 3N-5N | *Mylopharyngodon piceus, Cypriniformes* | 3 | 100.00% | - |
| T | Primate nuclear gene | 1T-275T | *Pan troglodytes* | 275 | 96.91% | 4.47% |
| P |  | 21P-28P | *Pan paniscus* | 8 | 98.23% | 2.15% |
|  |  | 29P-36P | *Gorilla gorilla* | 8 | 98.42% | 2.03% |
|  |  | 6P-20P | *Cercopithecidae* (Old World monkeys) | 15 | 94.31% | 2.42% |
|  |  | 2P-5P | *Platyrrhini* (New World monkeys) | 4 | 94.31% | 5.80% |
|  |  | 1P | *Carlito syrichta (Tarsiiformes)* | 1 | 70.67% | - |
| C |  | 1C-9C | Primate protein-coding sequences | 9 | 92.11% | 4.04% |
| A | Nuclear gene of non-indigenous organism | 1A-7A | *Bos mutus, Artiodactyla* | 7 | 93.67% | 5.51% |
|  |  | 8A-15A | *Meleagris gallopavo, Galliformes* | 8 | 92.06% | 8.02% |
| M | Nuclear gene of marine organisms | 1M | *Scylliorhinus canicula, Chondrichthyes* | 1 | 50% | - |
|  |  | 2M-5M | *Mercenaria mercenaria, Mollusca* | 4 | 70.89% | 17.23% |
|  |  | 6M-10M | Marine fishes, *Actinopterygii* | 5 | 57.79% | 18.61% |
| L | Others | 1L-4L | *Matches with very low similarity* | 4 | < 50% | - |

Note: The sequences between 7C and 28P, and between 8C and 4P, are identical.

#### 3. Investigating local geological changes through algal distribution

We identified three sequences matching Chlorophyta and five aligning with Streptophyta (Table S1). Sequences ID_1G, ID_2G, and ID_3G display unique profiles distinct from known genomes, hinting at their potential classification as aDNA. Morphological evidence from core samples of the Hetaoyuan Formation (37.5 Ma) indicates a marine invasion, with preserved skeletons of red algae (Rhodophyta) and brown algae (Ochrophyta) reported by previous studies ^6, 7^. In contrast, our data detected no DNA from these algal groups, implying a significant loss of algal DNA in the reservoir since early times. The small size of these organisms likely renders them prone to rapid dissolution and rupture upon incorporation into the oil, resulting in the release and subsequent degradation of their DNA.

#### 4. Evidence of marine organisms’ DNA signature

The analysis identified sequences containing genetic signatures of marine organisms, including Chondrichthyes, Actinopterygii, Tunicata, and Mollusca (Table S2), indicating the occurrence of local marine transgressions. Four sequences correspond to ray-finned fish species, with two sequences attributed to Scombriformes and two to Eupercaria. These marine fishes emerged during the mid-to-late Cretaceous period ^22^, suggesting a marine invasion occurred after this age. This evidence lends circumstantial support to Xia’s view ^6^.

#### 5. Evidence of ancient animals’ migration and continental drift

Fragments associated with non-native species, such as yaks in the northern Himalayan foothills, turkeys in the Americas, and kiwi birds in the South Pacific Islands, were discovered (Table S3). It is proposed that these species are descendants of ancient Pangaea ancestors who migrated and reproduced in their local habitats due to continental drift. Their eventual disappearance left molecular evidence underground, offering new insights into the theory of continental drift.

#### 6. Unveiling species intermediate forms in avian ancestry

The Late Mesozoic strata underlying the Hetaoyuan Formation are thought to date back to an era marked by dinosaur reproduction and the emergence of early birds. The ID_1L fragment aligns with avian genomes, showing a 61.91 affinity value to the mRNA transcript sequence of *Calidris pugnax*, suggesting it derives from an ancient bird (Table S4). This sequence includes the protein-coding regions of exon 3, exon 4, and part of exon 5 but lacks introns, indicating its role as a pseudogene. Detailed analysis reveals a composite structure: the initial 25 base pairs upstream closely resemble the intergenic sequence of *Podarcis lilfordi* (Figure 2), pointing to a genomic blend of avian and reptilian origins. This combination of dinosaurian and avian traits may represent the first molecular evidence of transitional species within Darwin’s evolutionary framework. The lack of an exact match to modern bird sequences could reflect either the extinction of the host species or significant genomic divergence in its descendants. This discovery further corroborates the hypothesis that the Nanyang Basin was an ancestral habitat for early birds.

**Figure 2.**
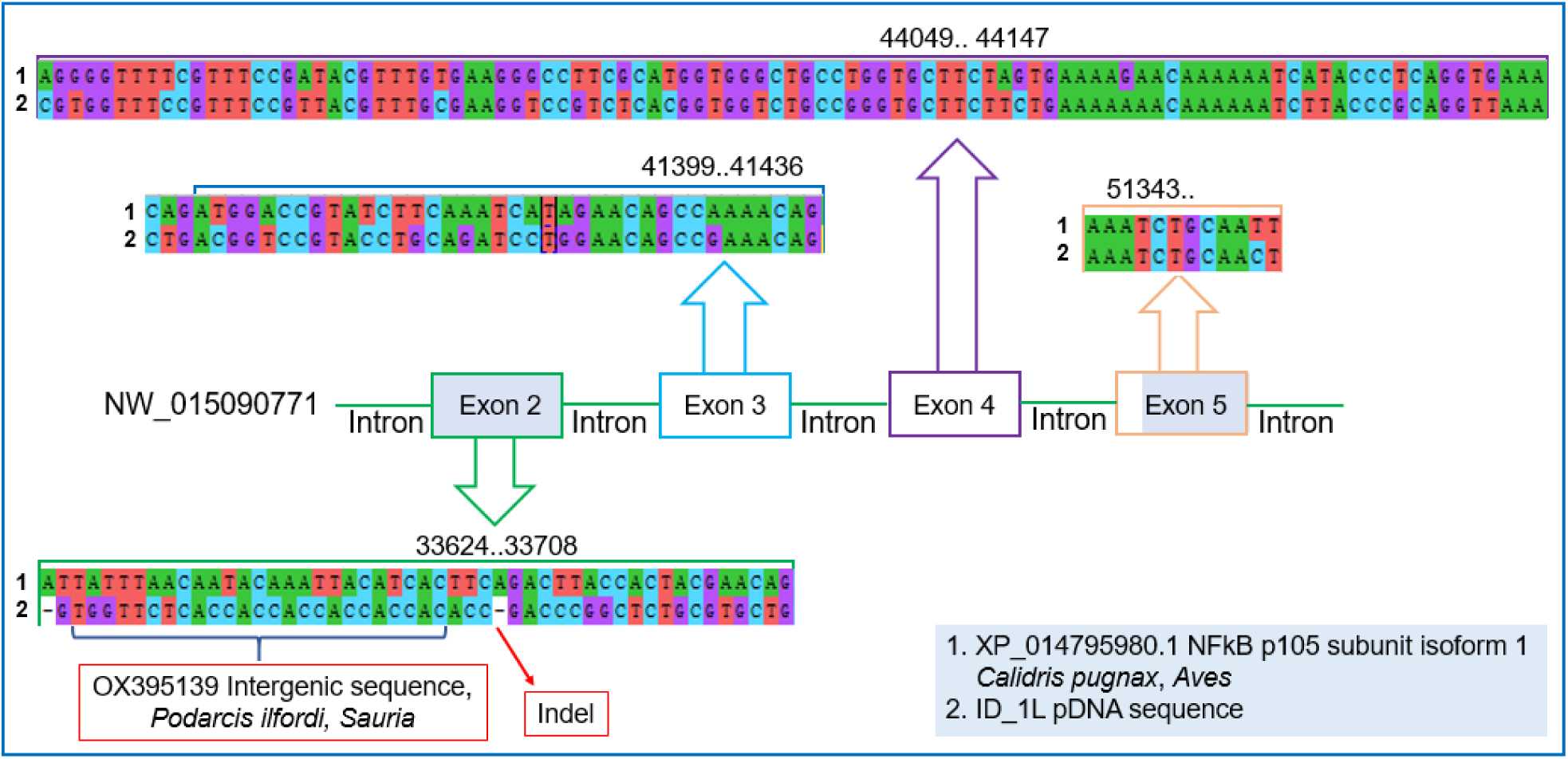
The Schematic diagram of *NFκB* p105 subunit-like DNAsequence (ID_1L)

#### 7. Discovery of unclassified *Homo* species via mitogenomes

We utilized the MS mode to analyze a subset of 1,047,532 human-related sequences (Figure 1) and discovered five non-primate mitogenomic sequences (Table S5) without non-human primate mitogenomic sequences. Of these, 845 sequences exhibiting 100% affinity to the human mitogenomes were excluded for further analysis. Additionally, 41 sequences showed divergent affinity values to the human mitogenome, ranging from 91.48% to 99.5%. Given that comprehensive human mitogenome data are available and potential misclassifications arising from minor unknown loci variations are rarely ruled out, these sequences are likely of ancient origin (Table S6).

Closer examination of two sequences, ID_17H and ID_20H, revealed 100% mutual coverage but distinct identities of 96.98% and 99.50%, respectively, suggesting they derive from separate maternal lineages. The pDNA fragment ID_18H, with an affinity value of 96.80%, was positioned via phylogenetic analysis on a node with *Homo sapiens*, hinting at a possible origin from an unrecognized species within the genus *Homo* (Figure S3, top). Fragment ID_19H, displaying an affinity value of 83.85% to human ND4, was phylogenetically placed between the human and chimpanzee genera, potentially representing the earliest local *Homo species*; RelTime-ML analysis estimated its divergence from humans at 0.49 Ma (Figure S3, middle & bottom). Notably, these pDNA sequences are absent from the modern human genome, indicating that the ancient humans bearing these fragments are likely extinct.

#### 8. Discovery of unclassified *Homo* species via nuclear genomes

In the pDNA, a large number of SSFs (with low affinities) aligned with nuclear sequences of non-human primates (NHPs) at varying frequencies: 1 with the Tarsiidae family, 4 with Platyrrhini (New World monkeys), 15 with *Macaca* (Old World monkeys), 8 with gorillas, 8 with pygmy chimpanzees (Table S7), and 271 with chimpanzees (Table S8). Among the 271 chimpanzee-aligned sequences, the second-hit species were modern humans in 241 cases (88.93%), other primates in 29 (10.70%), and non-primates in the remaining instances. Given the local fossil evidence of *Homo erectus* and the absence of non-human primate fossils in the region, we propose that these sequences maybe linked to Asian *Homo erectus*, particularly in light of the genomic relationships between modern humans and NHPs.

#### 9. Human activities and local eDNA

Unlike non-human apes, human burial practices have effectively slowed DNA decomposition, facilitating its preservation and accumulation in the environment surrounding burial sites. Fossils of *Homo erectus*, dated to approximately 0.5 Ma, have been uncovered in Nanzhao County near Nanyang oilfield ^19^. Based on this, we infer that the ID_19H pDNA fragment analyzed in this study, along with the majority of the 271 pDNA fragments previously noted, likely originated from Late Asian *Homo erectus*. Given that these DNA fragments are considerably "younger" than the 37.5 Ma age of the source rock, they are classified as paeDNA.

#### 10. Summary of the DNA Signatures of Late Asian *Homo erectus*

This study reveals that the mitogenomic divergence between Asian *Homo erectus* and modern humans is markedly smaller than that between Asian *Homo erectus* and chimpanzees. The study suggests that Asian *Homo erectus* are more closely related to humans on a maternal basis than to chimpanzees. They retained conservative skeletal traits, such as thick skulls and low foreheads, akin to Early African *Homo erectus*, with their extinction regarded as an "evolutionary dead end" ^23,24,25,26^. Previous studies indicate that ancient human populations experienced significant population bottlenecks and limited mitochondrial haplogroup diversity, organizing into small, female-centered clan tribes with strong kinship bonds ^27,28,29^. Determining the precise timing of reproductive isolation resulting from the genomic divergence between humans and chimpanzees remains a critical scientific question deserving further investigation.

Many SSFs matched the nuclear genes of non-human great apes (Table 1); however, central China is not a habitat for them. Consequently, these fragments likely originate from their closest relatives, Asian *Homo erectus*. The presence of a substantial number of NHP SSFs in oil samples is striking, suggesting a significant population of Asian *Homo erectus* during a prolonged period of residence in this region. Also, petroleum serves as an effective medium for preserving GAT genomes.

### IV. Hypothesized Pattern of DNA migration and preservation in oil reservoirs

DNA is predominantly water-soluble, with limited solubility in lipids. The lipid envelopes encasing cells and organelles facilitate their entry into oil reservoirs. Upon rupture of these envelopes, DNA intermingles with alkane molecules but is subsequently extracted into the aqueous phase at the water-oil interface upon contact with water. We hypothesize that tissues or cells, particularly fat tissue, adipocytes, or casein fragments, can readily infiltrate petroleum reservoirs. In more extensive remains, multiple layers of fat tissue may become encapsulated by oil, resulting in "oil immersion preservation". This process enables a gradual release of DNA from tissues adjacent to the oil phase, with successive layers contributing DNA as degradation progresses until the organism is fully dissolved. In addition, fat tissues containing DNA can liquefy, allowing DNA to flow into the reservoir, resulting in a prolonged, slow, and continuous release. In contrast, smaller entities like algae and plankton possess cells that rupture rapidly, quickly releasing their DNA into the reservoir. Consequently, pDNA experiences ongoing loss over time (Figure 1). This raises an intriguing question: Do the hosts of eDNA, including paeDNA and preDNA, also contribute to oil formation? Though their input may appear minor, the evidence suggests a positive response.

Research has noted a decline in oriDNA and an increase in eDNA, predominantly from microorganisms such as bacteria, in bone fossils ^30^. Over time, oriDNA and paeDNA in reservoirs undergo continuous degradation and loss, as eDNA persistently infiltrates the oil, creating a distribution of “old-less, new-many” for GAT genomes. Human activities, such as urban settlement, burial practices, and extensive cultivation of angiosperm crops, have further contributed to eDNA resources (Figure 3). The origins of oriDNA and paeDNA from the distant past remain inconclusive in this study due to a limited understanding of ancient paleoenvironments. In contrast, GMs are primarily oil-soluble, exhibiting minimal solubility in water. As a result, there is minimal concern regarding the loss of GMs from oil reservoirs, enhancing the effectiveness of their detection patterns in tracing the molecular origins of oil and gas. Although some environmental GMs may be continually incorporated, the “old-less, new-many” pattern observed in pDNA does not apply to GMs.

**Figure 3.**
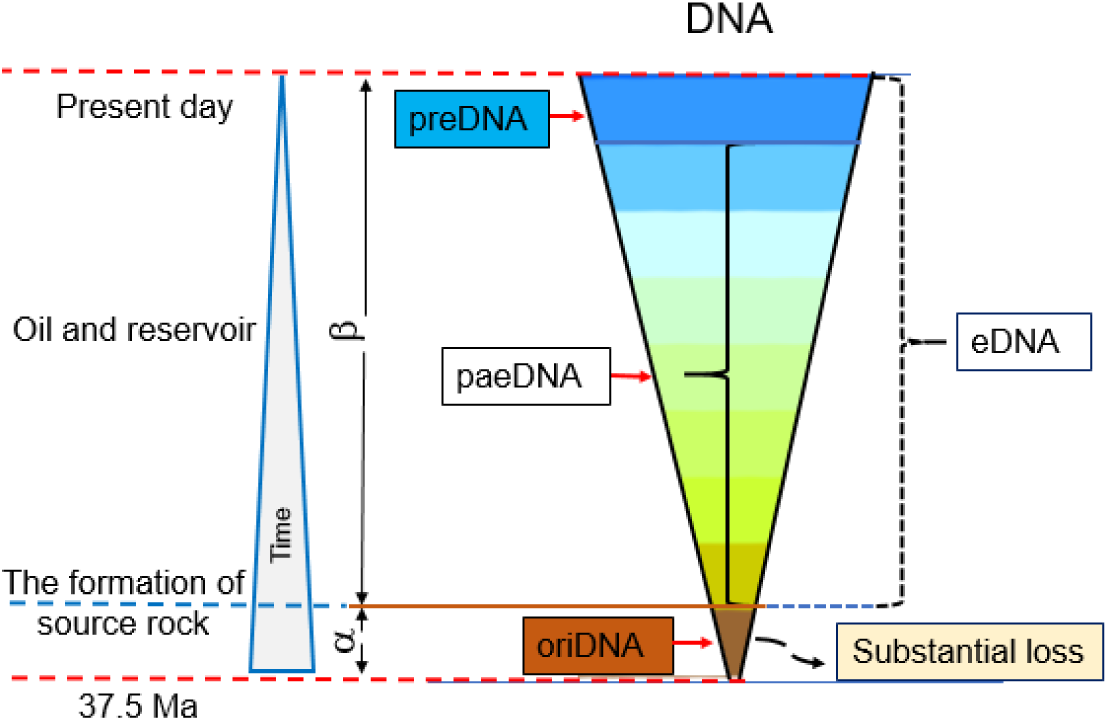
Preservation Patterns of DNA in Petroleum

Two examples highlight the “old-less, new-many” pattern of the GAT genome. First, DNA from lower life forms in crude oil has almost entirely disappeared, whereas DNA from modern species has progressively increased (Figure 1). Second, among 886 human mitogenome sequences, 845 from modern humans (preDNA) display a 100% affinity value, whereas 41 sequences from ancient *Homo species* (paeDNA) show varying degrees of divergence from the human mitogenome (Table S6).

### V. Challenges to aDNA authenticity criteria

In this study, many pDNA fragments within the GAT genome exhibited no signs of fragmental deamination. We propose several explanations for this observation. First, eDNA encapsulated in lipids enters the reservoir, an anaerobic and anhydrous environment that may inhibit deamination. Second, DNA deamination typically requires water as a critical component, which is scarce in this context. Third, fragments displaying severe deamination may have been excluded during the selection process of the mega screen method. Deamination is a critical marker for authenticating aDNA in the Pääbo method ^31, 32, 33^. However, Pääbo’s team rarely encounters material like petroleum in this study. Consequently, this primary indicator of aDNA authenticity does not apply to pDNA studies, highlighting the need to reassess the scope of applicability of this method.

### VI. Significance and outlook

#### Scientific Importance

Genetic information derived from DNA is essential for elucidating species relationships, life histories, and evolutionary mechanisms. This study marks the first successful extraction of DNA from petroleum, expanding the scope of molecular biology and challenging established disciplinary boundaries. It overturns the long-standing misconception that petroleum lacks DNA and provides the first molecular evidence demonstrating that Asian *Homo erectus* is more closely related to modern humans than to Chimpanzees in maternal side. Additionally, it contributes to geological theory by offering molecular evidence supporting marine invasion events in the Nanyang region. While Xia’s evidence based on the algae alone has been deemed insufficient to confirm such events and remains not widely accepted, our findings bolster this hypothesis ^6,7^. Moreover, this study facilitates the detection of intermediate animals, such as the ID_1L fragment mentioned before, which suggests activities of local avian ancestors (possibly Archaeopteryx?). This evidence significantly aids in addressing Darwin’s puzzle, the scarcity of intermediate species in the fossil record, revealing that molecular evolution may exhibit greater diversity than previously recognized.

#### Petroleum as an SSF Reservoir

DNA sequences show remarkable similarities across different species. For example, humans and chimpanzees share over 98% of their genomic sequence. This closeness highlights the importance of using SSFs for accurate identification. Although SSFs represent only a little portion of a genome, just a few pieces from a sample are enough to confirm the presence of a particular species while also functioning as valuable geological markers. Petroleum SSFs are indicators for exploring ancient animal activity patterns, spanning lower life forms, birds, and advanced mammals. As a significant amount of SSFs was identified in the fragments of the petroleum GAT genome, this quantity significantly exceeds that found in other sample types (e.g., fossils, rocks), highlighting petroleum ‘s role as a storage medium and an untapped reservoir for aDNA. Likewise, using the SSFs, we revealed traces of ancient animal activities from locally extinct species, such as yaks, Kiwi birds, sea fishes, and shellfish, laying the groundwork for reconstructing paleoenvironments and paleoecology.

#### Future Directions

Looking ahead, we aim to extract aDNA from petroleum, fossils, and source rocks to establish an integrated trinity of global aDNA libraries. This endeavor will pave the way for an innovative aDNA resource, enhancing our understanding of evolutionary and geological histories.

## Methods

### I. The geochemical survey

The oils were deasphalted with petroleum ether and then fractionated by column chromatography using a 2:1 (v/v) mixture of silica gel and alumina. The fractions were separated into saturated, aromatic, and resin components by sequential elution with 60 mL of petroleum ether, 40 mL of a dichloromethane/petroleum ether mixture (2:1, v/v), and 30 mL of a dichloromethane/methanol solution (93:7, v/v), respectively.

Gas chromatography (GC) analysis was conducted on the saturated fractions using an Agilent Model 6890 gas chromatograph installed with a silica column (HP-1, 30 mm × 0.25 mm i.d.). The oven temperature was programmed to increase from 100°C to 300°C for 10 minutes at a rate of 4°C/min. Helium was employed as the carrier gas.

Gas chromatography-mass spectrometry (GC-MS) analysis of the saturated and aromatic fractions was performed using an Agilent Model 6890 gas chromatograph coupled to a DB-5MS capillary column. The temperature program for the saturated fraction was as follows: the initial temperature was set at 100 °C for 1 minute, followed by a ramp of 4°C per minute to 220°C, and then increased to 300°C at a rate of 3°C per minute and held for 15 minutes. The mass spectrometer was operated in electron impact mode, with an ionization energy of 70 eV and a scan range of 50-600 Da. The biomarker parameters of the saturated and aromatic hydrocarbons were determined by integrating the peak areas on the mass chromatograms.

### II. DNA extraction, DNA library construction, sequencing, and NCBI nucleotide BLAST

The procedure was conducted in a BSL-2 industrial laboratory (Biosafety Level 2), following rigorous ancient DNA extraction protocols ^29^. The process involved the UV sterilization of all items used and the utilization of Geobio ^®^ DNA Remover (Jiaxing Jiesai Biotechnology Co., Ltd., Jiaxing, China) on clean benches and solid equipment. It is important to note that this laboratory has never been exposed to fish, NHPs, or plant samples before this work.

To extract DNA from petroleum samples, we followed the recommended protocol of the Geobio^®^ DPPS-PMNP DNA extraction kit developed by Jiaxing Jiesai Biotechnology Co ^34^. Due to various long-term factors, some pDNA molecules are cross-linked with other macromolecules, forming complexes. To estimate the extract concentration, we measured the optical density(OD) at the UV 260 nm peak of the DNA mix, containing DNA complexes and free DNA molecules. We performed the extraction experiments on the samples five times, and the results were consistent (Table <u>S9</u>), providing sufficient DNA (Table S10). We used NEB PreCR^®^Mix for DNA repair.

The NGS DNA library construction followed the SSDLSP standard procedure (Sangon Biotech, CORP). Sequencing was conducted on Illumina NovaSeq 6000 (S-4 Reagent Kit) using the “Dual index sequencing” standard procedure, and the reads met the quality control standards with satisfactory quality distribution. Sometimes, it’s necessary to manually remove extra adapter sequences at both ends of the read to obtain accurate information. After merging the forward and reverse reads, we performed local NCBI Nucleotide BLAST, Size 11, using the database (Version 5, Nucleotide sequence Database), identifying 3,159,020 valid sequences. The sequences were classified according to their biological characteristics, and their lengths were measured (Table S11). It is important to note that the processes of secondary oil collection may inadvertently lead to the loss of many aDNA fragments.

#### The mega screen method

First, we employed the Best E-value mode to group sequences by aligning fossil DNA with all known sequences from the NCBI database (Version 5, Nucleotide Sequence Database). We applied an E-value cutoff below 1E-07 to filter for qualified sequences (QS) and identify the best match at a taxonomic level.

In the second step, we used the MS mode by selecting the "exclude" option and conducting a search that omitted the first result. If the E-value difference between two searches is less than 1E-02 (meaning the quotient of the first E-value divided by the second one is ≤ 0.01), the fragment is considered specific to the first match, confirming its specificity. Conversely, if the difference exceeds the established threshold (1E-02), the fragment is not confirmed as originating from a single one. In this scenario, we cannot determine the fragment’s origin.

The whole process involves the following steps:

1. Nucleotide Blast is employed to align sequences to the entire NCBI database without any limitation.
2. Creation of subsets for sorting the sequences: Using the Best E-value mode, results are matched, and sequence fragments are grouped into the appropriate subsets, determining the total number of sequences (TS). Subsequently, an E-value threshold of < 1E-07 is established to filter for qualified sequences (QS).
3. The subset primarily containing aDNA is selected.
4. Each sequence within this subset is screened individually using the MS mode to identify those originating from a single species, categorizing the “unique” sequences (SSFs).
5. Considering various factors, including local climate shifting, geological changes, and evolutionary principles of the host species, we can ascertain if the sequence is oriDNA.

#### Sequence affinity (Affinity)

The metrics we set up to assess the similarity relationship between subject sequences and hit genomes were obtained by multiplying the two values of Identity and Cover obtained from NCBI BLAST and then converted to percentages (Identity× Cover × 100%). Mutations in the genomic and mitogenomic sequences occur continuously during the evolutionary process, leading to significant sequence differences between ancient and modern species. However, this variation is lower in conserved sequences than in non-conserved sequences. A high-affinity value indicates that the sequence is highly coherent and related to the genome of the modern species, which is likely to be either a conserved sequence or a modern eDNA. A low-affinity value indicates a low similarity, and the sequence may be from a distantly related species either has or has not yet been sequenced. It is necessary to consider various factors, including the sequence composition and conservation, host species information, host genome sequencing data, etc., rather than relying on the Affinity value alone when determining whether a sequence is an oriDNA.

## Data and software availability

The sequencing data have been deposited in the NCBI BioProject (PRJNA1091869). The pDNA sequences utilized in this manuscript are listed in the Appendix (Table S12). In this study, MEGA11 was used to construct the RelTime-ML local time tree ^36^.

## Acknowledgments

We extend our gratitude to the "Chao-Xiang Talent Program" foundation of Haining City, the support that covers the costs of petroleum DNA extraction and DNA sequencing from 2018 to 2022. The collection of petroleum samples from 2018 to 2021 received support from the Key Research & Development Special Program of Henan Province (231111111200), with the indispensable help of numerous workers and administrative staff from Nanyang oilfield (Sinopec Corp.), including Ms. Yu Liu, Mr. Pin-Lei Cai, and Mr. Peng Wei. Special thanks to Haining BoShang Biotechnology Corporation and Sangon Biotech (Shanghai) Corporation for their collaborative support in the success of this project, especially in the construction of the NGS DNA library using the SSDLSP standard procedure and sequencing. We also appreciate the contributions of Mr. HAN-Yu Zhu in creating a paleogeographic map and the assistance of Ms. Yun-Li Wang and Ms. Yue Zhang in processing the data. We extend our heartfelt appreciation to the following individuals for their invaluable contributions to this project: Dr. Hua-Jun Zheng from the Shanghai Institute for Biomedical and Pharmaceutical Technologies, Dr. Jun-Ye Ma from the Nanjing Institute of Geology and Paleontology, at the Chinese Academy of Sciences, Dr. Alida M. Bailleul from the Institute of Vertebrate Paleontology and Paleoanthropology, at the Chinese Academy of Sciences, Dr Shun-Ping He from the Institute of Hydrobiology, also at Chinese Academy of Sciences. Their stimulating discussions regarding the manuscript have greatly enriched our work.

## Author contribution

The study was conceptualized and designed by W.Q.Z. and Y.Q.G., Y.Q.G., T.C.Q., and W.Q.Z. supervised the transportation of oil samples. M.J.L. performed the geochemical marker analysis. Geographical and paleogeological data analysis, as well as map creation, were conducted by F.L. and H.Y.Z. Petroleum DNA extraction from the samples was carried out by W.Q.Z., C.Y.Z., and J.Y.Y. Alignment between the sequences and present genomes was completed by Z.Y.G. and M.J.C. DNA laboratory analysis, interpretations, taxonomic profiling, and annotation were conducted by Z.Y.G., Z.Y.T., S.J.Z, L.J.Z, and W.Q.Z. The statistical analyses were performed and finished by G.Q.C., X.G.Z., R.C., and J.H.Z. Phylogenetic analyses of mitogenomic DNA sequences were performed by J.H.G., under the supervision of G.Q.C. and W.Q.Z. Figures were designed and finished by R.C., J.H.Z., L.Y.Q., and M.R.S. The manuscript was written by W.Q.Z. and corrected by Z.Y.T., M.J.L., and L.Y.Q. The Project Coordinator was T.C.Q.

## Data and materials availability

All data are available in the main text or the supplementary materials.

## Supplementary Materials

**Figure S1.**
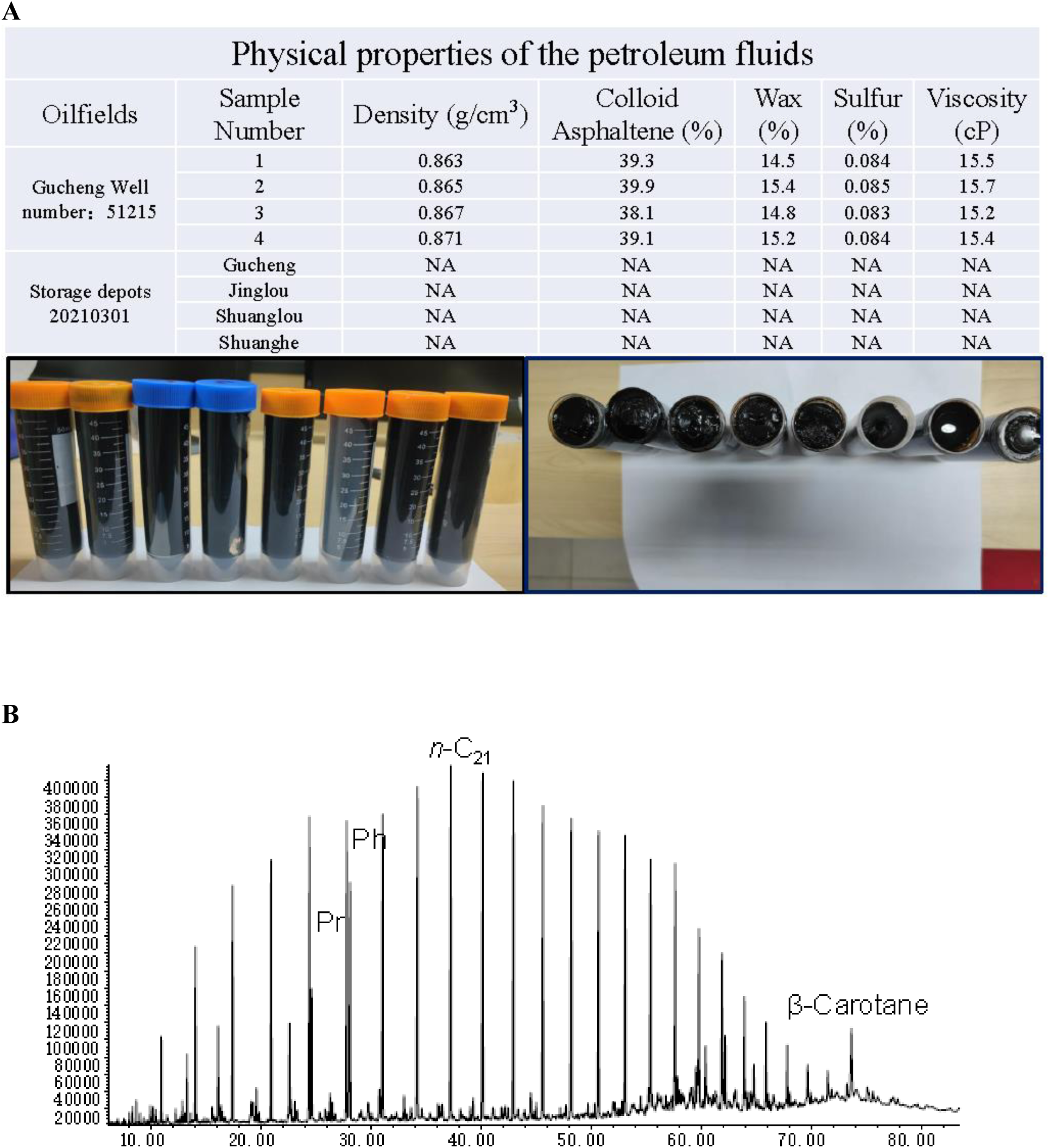
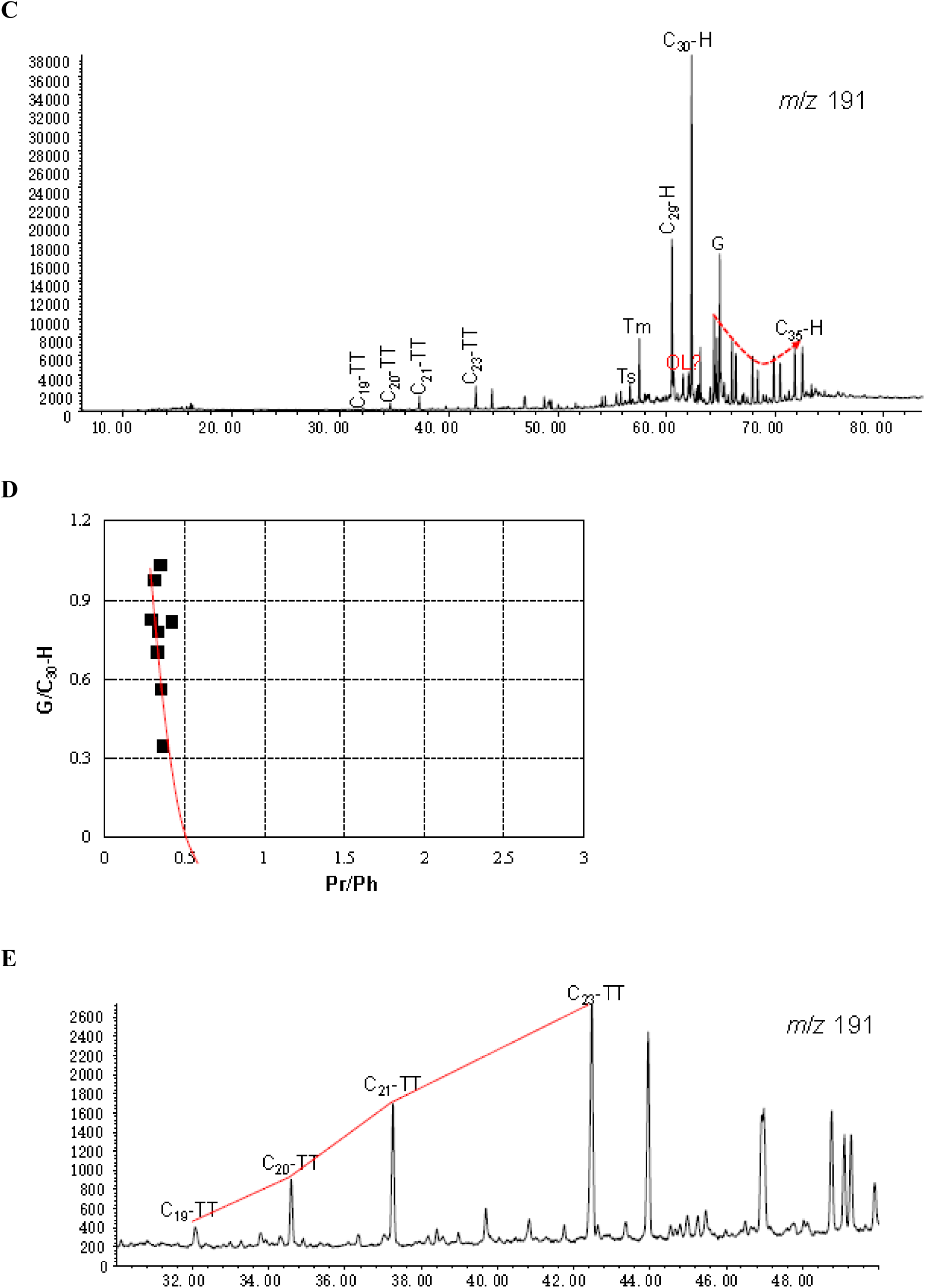
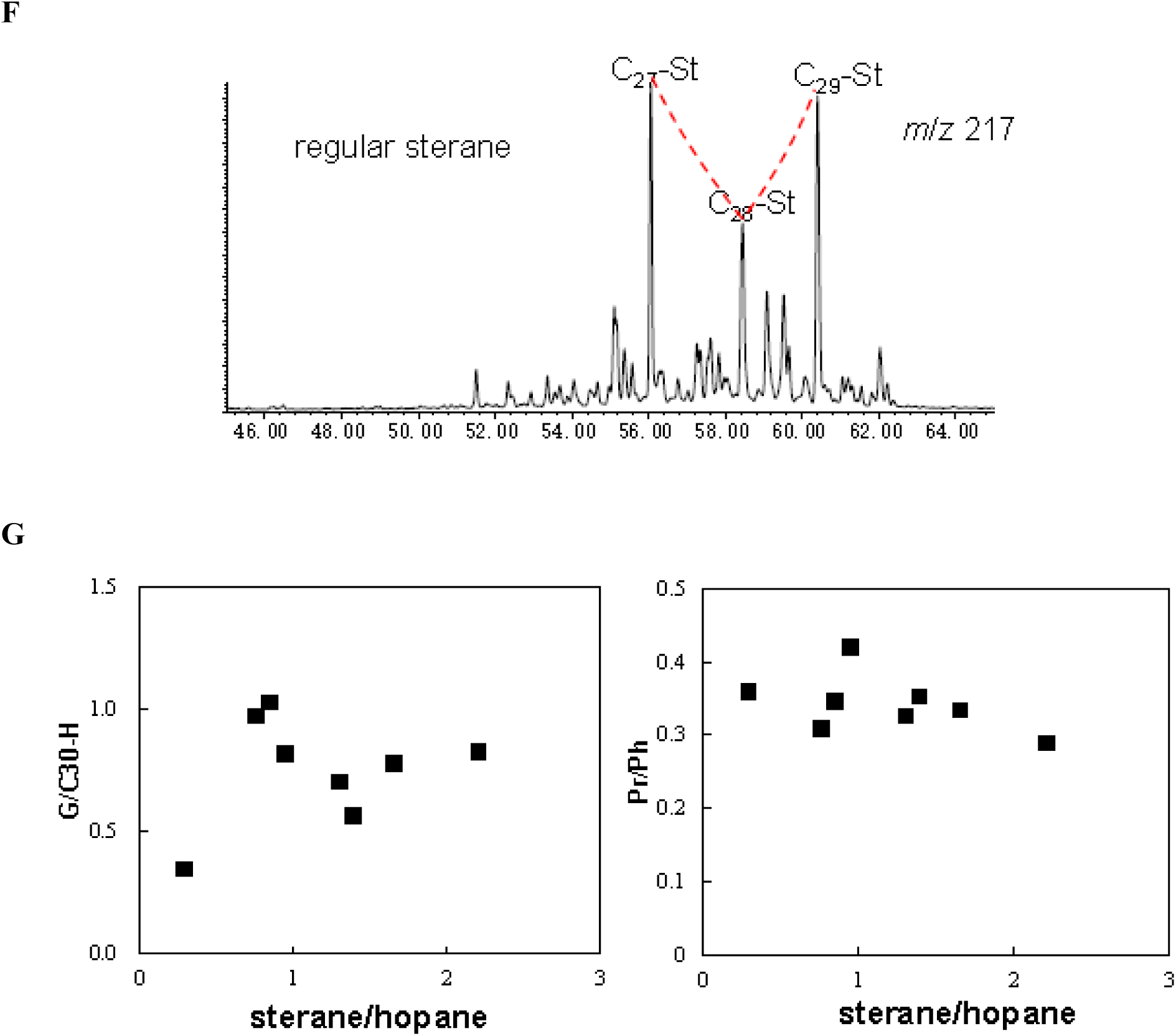
Geochemical analysis of petroleum samples **A**, the buk geochemical characteristics of the petroleum used in this study. **B**, the distribution of *n*-alkanes and isoprenoids in the petroleum used in this study. **C**, the distribution of hopanes in the petroleum used in this study. **D**, the distribution of hopanes-gammacerane in the petroleum used in this study. **E**, the distribution of tricyclic terpanes in the petroleum used in this study. **F**, the distribution of sterane in the petroleum used in this study. **G**, the distribution of sterane in the petroleum used in this study.

**Figure S2.**
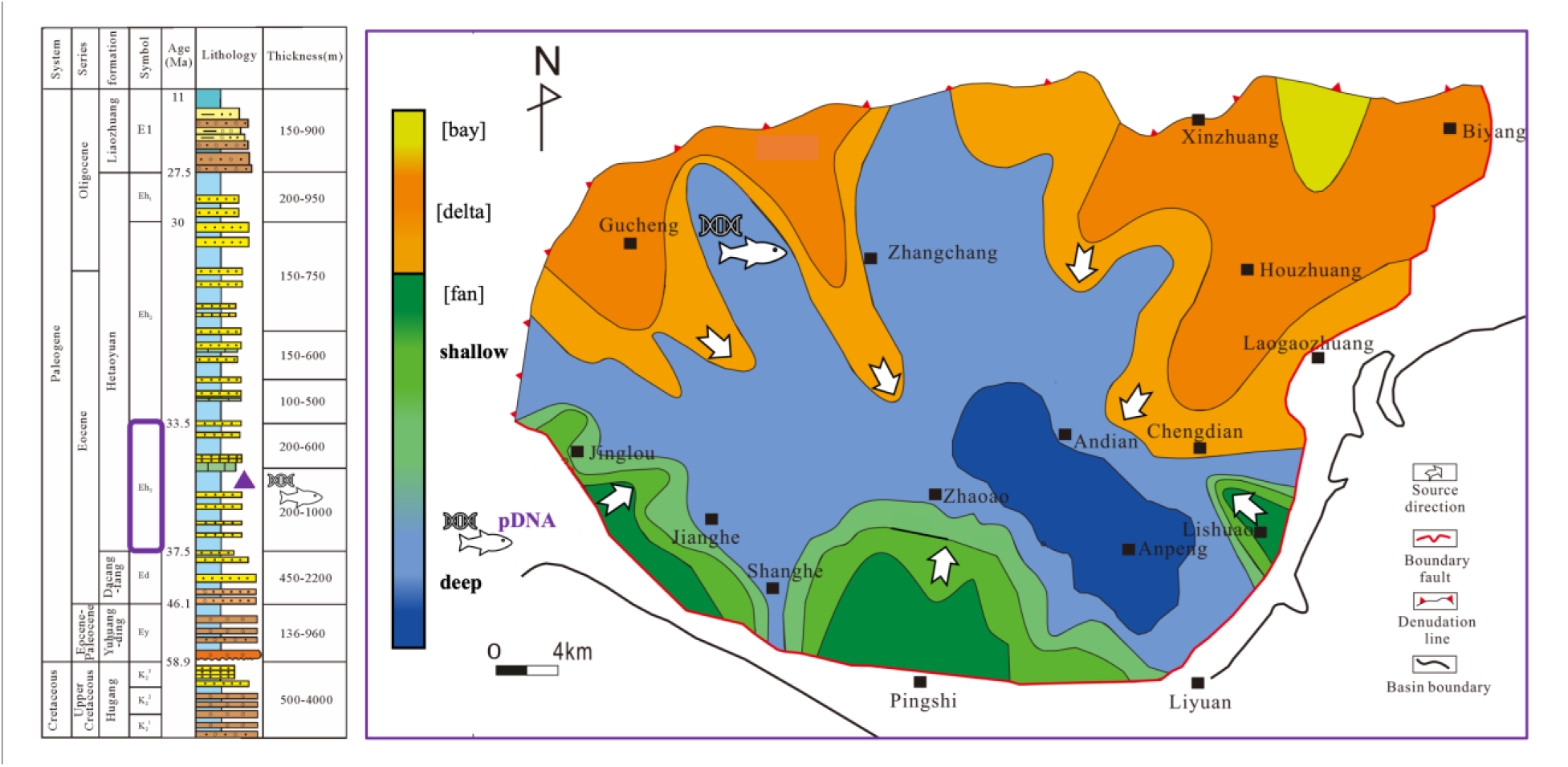
The lithostratigraphy of the Hetaoyuan Formation (left) and corresponding paleogeography in the Eocene ages (right) at the sample position. The paleogeographic map is modified from Li‘s work ^1–3^, based on the corrective information from Dong’s work^4^ and Xia^5^.The samples are located in Member 3 of the Hetaoyuan Formation.

---

**Figure S3.**
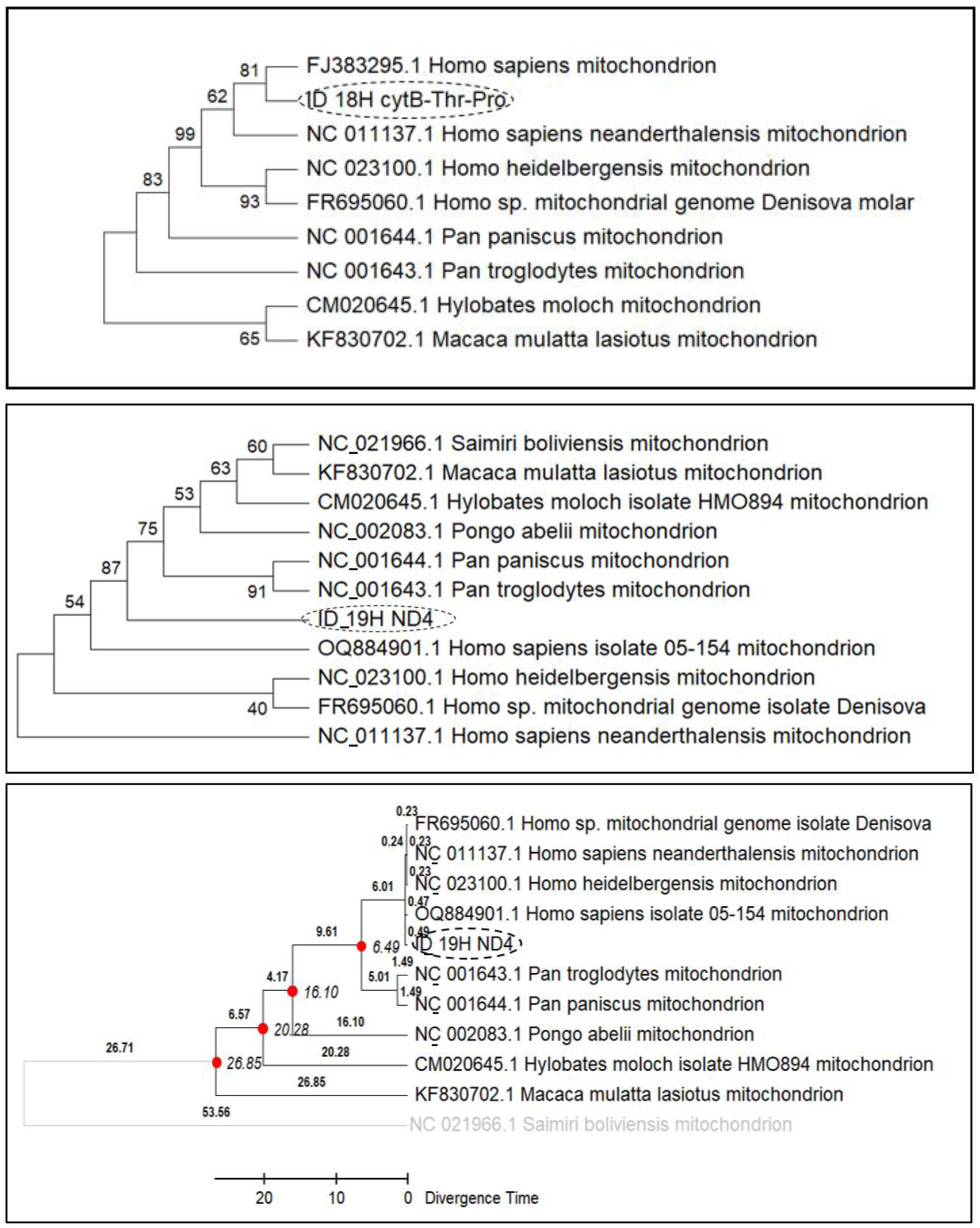
The phylogenetic analysis of the pDNA sequences The phylogenetic analyses are established using primate mitogenomic sequences as references. The data is being examined utilizing the Maximum Likelihood method in MEGA11 on a locally installed computational platform. The divergences are evaluated in Bootstrap consensus trees through specific calculations with 1000 bootstrap replicates under the HKY+G model and a Timetree under RelTime - ML mode. In each context, ID_18H (shown in the top) and ID_19H (shown in the middle and the bottom) correspond to the human cytB+Thr+Pro and ND4 regions, respectively (OQ884901.1). The Bootstrap value of the node on the Bootstrap consensus tree and the divergence time near the node on the Timetree are numerically represented and displayed for analysis.

**Figure S4.**
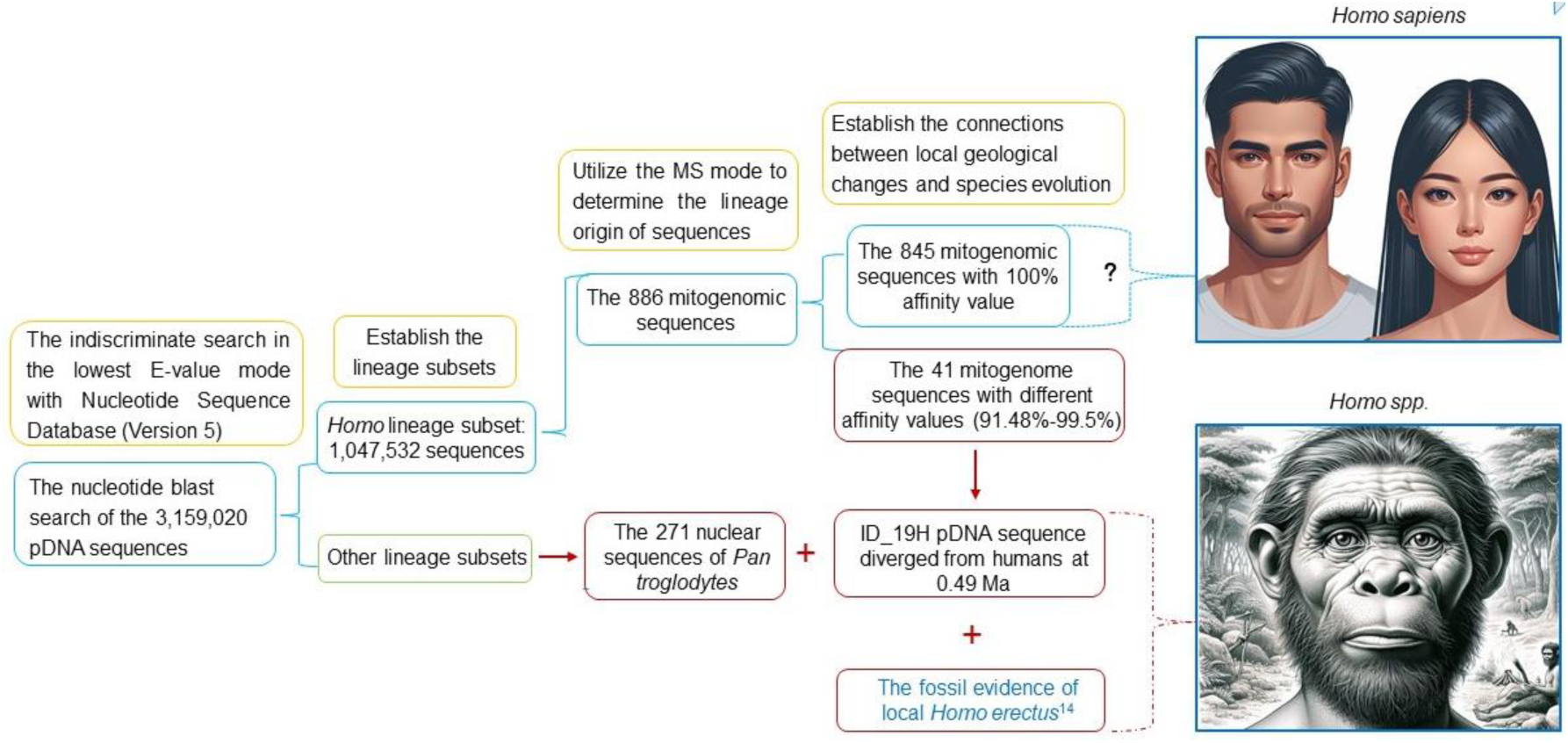
The molecular evidence for the presence of unclassified *Homo species*

**Table S1.**
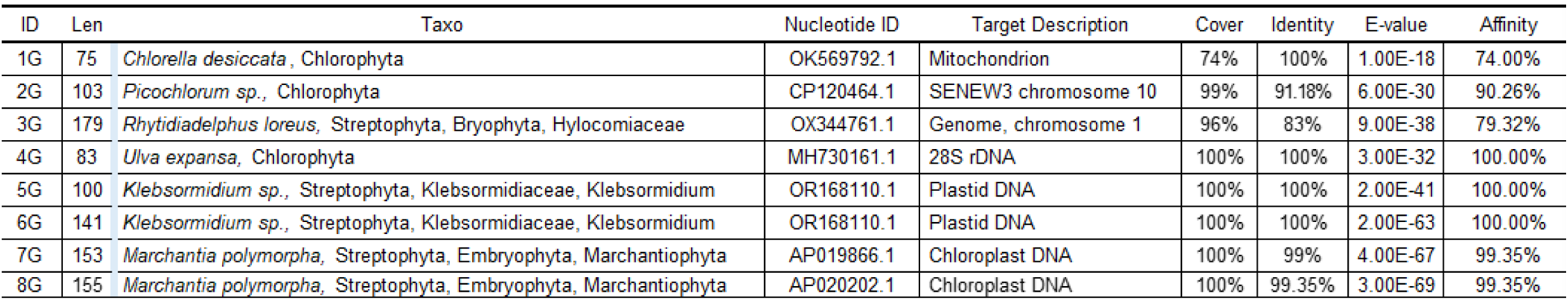
The pDNA sequences corresponding to low plant genomes.

**Table S2.** The pDNA sequences corresponding to the marine organisms’ genomes.

| pDNA |  | Results of the Nucleotide Blast |  |  |  |
| --- | --- | --- | --- | --- | --- |
| ID | Len | Taxo | Nucleotide | E-value | Affinity |
| 1M | 78 | *Scyliorhinus canicula*, Chondrichthyes | LR744051.1 | 2.00E-06 | 50% |
| 2M | 80 | *Mercenaria mercenaria*, Mollusca | XM_045350734.2 | 9.00E-18 | 83% |
| 3M | 92 | *Nucula nucleus*, Mollusca | LC123061.1 | 5.00E-22 | 78% |
| 4M | 112 | *Ruditapes philippinarum*, Mollusca | KU682310.1 | 2.00E-16 | 77% |
| 5M | 184 | *Phorcus lineatus*, Mollusca | OV121097.1 | 2.00E-07 | 45% |
| 6M | 253 | *Lateolabrax maculatus*, Actinopterygii | CP027279.1 | 2.00E-52 | 68% |
| 7M | 203 | *Hyperoplus immaculatus*, Actinopterygii | OX442354.1 | 1.00E-29 | 44% |
| 8M | 187 | *Thunnus maccoyii*, Actinopterygii | OU343202.1 | 8.00E-19 | 70% |
| 9M | 83 | *Thunnus maccoyii*, Actinopterygii | OU343200.1 | 9.00E-12 | 72% |
| 10M | 131 | *Aplidium turbinatum*, Tunicata | OU964919.12 | 3.00E-09 | 35% |

**Table S3.** The 15 pDNA sequences corresponding to the genomes of non-indigenous organisms.

| pDNA |  | Results of the Nucleotide Blast |  |  |  |
| --- | --- | --- | --- | --- | --- |
| ID | Len | Taxo | Nucleotide ID | E-value | Affinity |
| 1A | 117 | *Bos mutus* | CP027076.1 | 3.00E-46 | 97.44% |
| 2A | 149 | *Bos mutus* | CP027076.1 | 8.00E-55 | 92.98% |
| 3A | 144 | *Bos mutus* | CP027076.1 | 2.00E-61 | 96.62% |
| 4A | 97 | *Bos mutus* | CP027076.1 | 5.00E-23 | 81.66% |
| 5A | 162 | *Bos mutus* | CP027078.1 | 3.00E-67 | 95.92% |
| 6A | 119 | *Bos mutus* | CP027078.1 | 3.00E-46 | 96.64% |
| 7A | 140 | *Bos mutus* | CP027087.1 | 9.00E-54 | 94.45% |
| 8A | 223 | *Meleagris gallopavo* | OW982293.1 | 4.00E-93 | 92.34% |
| 9A | 151 | *Meleagris gallopavo* | OW982293.1 | 2.00E-49 | 95.26% |
| 10A | 117 | *Meleagris gallopavo* | OW982293.1 | 8.00E-41 | 97.20% |
| 11A | 198 | *Meleagris gallopavo* | OW982293.1 | 5.00E-53 | 94.28% |
| 12A | 288 | *Meleagris gallopavo* | OW982293.1 | 9.00E-128 | 97.20% |
| 13A | 92 | *Meleagris gallopavo* | OW982297.1 | 2.00E-20 | 72.90% |
| 14A | 154 | *Meleagris gallopavo* | OW982293.1 | 4.00E-53 | 95.26% |
| 15A | 103 | *Apteryx mantelli* | LK358553.1 | 2.00E-08 | 39.05% |

**Table S4.**
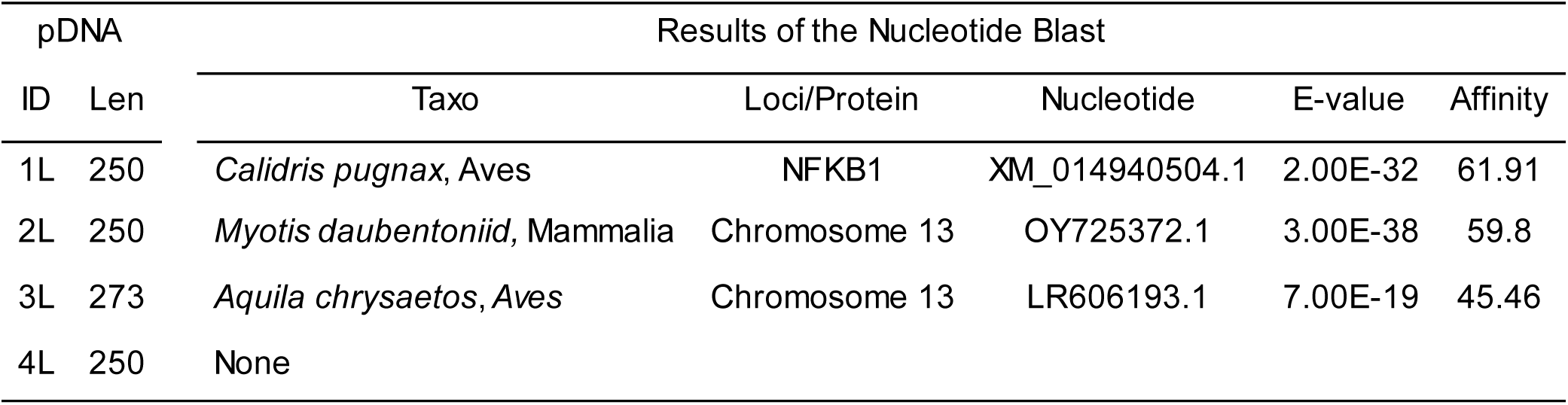
The pDNA sequences exhibiting low affinity values.

**Table S5.** The 5 pDNA sequences corresponding to non-primate mitogenomes.

| pDNA |  | Results of the Nucleotide Blast |  |  |  |  |
| --- | --- | --- | --- | --- | --- | --- |
| ID | Len | Taxo | Nucleotide | Range/Area | E-value | Affinity |
| 1N | 146 | *Gallus gallus*, Galliformes | OP740918.1 | 503-648; 12S | 2.00E-66 | 99.3200% |
| 2N | 193 | *Sus scrofa*, Artiodactyla | MN124249.1 | 344-536; COX1 | 2.00E-92 | 99.4800% |
| 3N | 81 | *Mylopharyngodon piceus*, Cypriniformes | MF687133.1 | 13586-13666; ND5 | 3.00E-32 | 100.0000% |
| 4N | 203 | *Ctenopharyngodon Idella*, Cypriniformes | MG827396.1 | 10382-10582; ND4 | 6.00E-97 | 100.0000% |
| 5N | 113 | *Ctenopharyngodon Idella*, Cypriniformes | MG827396.1 | 400-288; 12S | 2.00E-48 | 100.0000% |

**Table S6.** The 41 pDNA sequences corresponding to human mitogenome.

| pDNA |  | Results of the Nucleotide Blast |  |  |  |  |
| --- | --- | --- | --- | --- | --- | --- |
| ID | Len | Taxo | Nucleotide ID | Mito Range/Area | E-value | Affinity |
| 1H | 249 | *Home sapiens* | OR182493.1 | 13856-14104; ND5 | 2.00E-125 | 99.60% |
| 2H | 238 | *Home sapiens* | ON422324.1 | 8347-8584; ATP8+ATP6 | 2.00E-119 | 99.58% |
| 3H | 216 | *Home sapiens* | MH449153.1 | 9787-10002; COX3+trnG | 3.00E-107 | 99.54% |
| 4H | 215 | *Home sapiens* | OQ388438.1 | 47-261; D-loop | 1.00E-106 | 99.53% |
| 5H | 212 | *Home sapiens* | ON946746.1 | 765-554; D-loop | 5.00E-105 | 99.53% |
| 6H | 194 | *Home sapiens* | MH449153.1 | 9974-9781; COX3 | 5.00E-95 | 99.48% |
| 7H | 186 | *Home sapiens* | MG660585.1 | 8790-8975; ATP6 | 1.00E-90 | 99.46% |
| 8H | 176 | *Home sapiens* | JQ664512.1 | 4404-4578; trnM+ND2 | 5.00E-85 | 98.44% |
| 9H | 164 | *Home sapiens* | MN503417.1 | 11624-11787; trnH+trnS+trnL+ND5 | 2.00E-76 | 99.39% |
| 10H | 153 | *Home sapiens* | MN207699.1 | 652-804; D-loop: | 2.00E-72 | 99.35% |
| 11H | 111 | *Home sapiens* | OR182493.1 | 13525-13415; ND5 | 3.00E-49 | 99.10% |
| 12H | 200 | *Home sapiens* | LS997688.1 | 28-227; D-loop: | 1.00E-96 | 99.00% |
| 13H | 280 | *Home sapiens* | MF362729.1 | 11759-12038; ND4 | 7.00E-140 | 98.93% |
| 14H | 245 | *Home sapiens* | OQ707209.1 | 8658-8902; ATP6 | 6.00E-120 | 98.78% |
| 15H | 199 | *Home sapiens* | MG660585.1 | 7993-8191; COX2: | 1.00E-92 | 98.49% |
| 16H | 189 | *Home sapiens* | KF162400.1 | 8394-8206; ATP8 | 2.00E-89 | 98.41% |
| 17H | 199 | *Home sapiens* | OR182493.1 | 13537-13735; ND5 | 2.00E-89 | 96.98% |
| 18H | 250 | *Home sapiens* | FJ383295.1 | 16068-15820; CYTB+trnT+trnP | 1.00E-113 | 96.80% |
| 19H | 177 | *Home sapiens* | OQ884901.1 | 11485-11325; ND4 | 5.00E-60 | 83.85% |
| 20H | 199 | *Home sapiens* | OR182493.1 | 13537-13735; ND5 | 8.00E-98 | 99.50% |
| 21H | 189 | *Home sapiens* | OR182493.1 | 15872-15684; CYTB | 3.00E-92 | 99.47% |
| 22H | 188 | *Home sapiens* | OR159678.1 | 3294-3481; trnL-ND1 | 1.00E-91 | 99.47% |
| 23H | 179 | *Home sapiens* | OR182493.1 | 14442-14620; ND6 | 1.00E-86 | 99.44% |
| 24H | 175 | *Home sapiens* | OR159678.1 | 4656-4482; ND2 | 2.00E-84 | 99.43% |
| 25H | 137 | *Home sapiens* | OR159678.1 | 3223-3359; 16S+trnL+ND1 | 2.00E-63 | 99.27% |
| 26H | 128 | *Home sapiens* | OQ707209.1 | 10974-10847; ND4 | 1.00E-58 | 99.22% |
| 27H | 128 | *Home sapiens* | LC733705.1 | 10847-10974; ND4 | 1.00E-58 | 99.22% |
| 28H | 127 | *Home sapiens* | OR182493.1 | 11643-11769; ND4 | 5.00E-58 | 99.21% |
| 29H | 123 | *Home sapiens* | OR182493.1 | 3674-3796; ND1 | 8.00E-56 | 99.19% |
| 30H | 118 | *Home sapiens* | OR182493.1 | 3430-3547; ND1 | 5.00E-53 | 99.15% |
| 31H | 106 | *Home sapiens* | OR182493.1 | 14567-14672; ND6 | 2.00E-46 | 99.06% |
| 32H | 199 | *Home sapiens* | OR182493.1 | 15529-15727; CYTB | 4.00E-96 | 98.99% |
| 33H | 99 | *Home sapiens* | OR182493.1 | 13226-13128; ND5 | 1.00E-42 | 98.99% |
| 34H | 83 | *Home sapiens* | OR182493.1 | 11867-11785; ND4 | 8.00E-34 | 98.80% |
| 35H | 173 | *Home sapiens* | OR182493.2 | 3320-3148; 16S+trnL+ND1 | 4.00E-80 | 98.27% |
| 36H | 200 | *Home sapiens* | OR182493.1 | 12487-12288; trnL+ND5 | 2.00E-93 | 98.00% |
| 37H | 259 | *Home sapiens* | OR182493.1 | 14124-14382; trnL+ND5 | 5.00E-121 | 97.30% |
| 38H | 143 | *Home sapiens* | OR182493.1 | 13614-13754; ND5 | 2.00E-67 | 98.00% |
| 39H | 152 | *Home sapiens* | MG660585.1 | 7949-8040/COX2: | 4.00E-65 | 96.71% |
| 40H | 225 | *Home sapiens* | ON422324.1 | 8367-8589/ATP8-ATP6 | 5.00E-81 | 90.57% |
| 41H | 123 | *Home sapiens* | OR182493.1 | 3674-3796/ND1 | 8.00E-56 | 99.19% |

**Table S7.** The 36 pDNA sequences aligning with non-human primate nuclear sequences.

| pDNA |  | Results of the Nucleotide Blast |  |  |  |
| --- | --- | --- | --- | --- | --- |
| ID | Len | Taxo | Nucleotide | E-value | Affinity |
| 1P | 150 | *Carlito syrichta* | XM_021713054.1 | 3.00E-20 | 70.67% |
| 2P | 122 | *Sapajus apella* | XR_004259191.1 | 8.00E-41 | 92.62% |
| 3P | 63 | *Callithrix jacchus* | AC188585.3 | 5.00E-19 | 96.83% |
| 4P | 160 | *Sapajus apella* | XM_032273219.1 | 1.00E-47 | 84.12% |
| 5P | 164 | *Aotus nancymaae* | XM_012443881.1 | 1.00E-68 | 95.96% |
| 6P | 166 | *Macaca thibetana* | XM_050789932.1 | 4.00E-66 | 95.18% |
| 7P | 187 | *Macaca mulatta* | AC188451.1 | 1.00E-86 | 99.47% |
| 8P | 93 | *Macaca mulatta* | AC187507.2 | 1.00E-29 | 93.55% |
| 9P | 178 | *Macaca mulatta* | AC214608.1 | 3.00E-67 | 93.82% |
| 10P | 121 | *Macaca mulatta* | AC279264.1 | 4.00E-45 | 95.87% |
| 11P | 150 | *Macaca mulatta* | AC211796.4 | 3.00E-61 | 96.67% |
| 12P | 262 | *Macaca mulatta* | AC204076.9 | 1.00E-99 | 92.37% |
| 13P | 118 | *Macaca mulatta* | OM502028.1 | 3.00E-40 | 97.17% |
| 14P | 139 | *Macaca thibetana* | XM_050772658.1 | 2.00E-48 | 90.72% |
| 15P | 149 | *Macaca nemestrina* | XR_003018358.1 | 7.00E-56 | 94.63% |
| 16P | 118 | *Macaca mulatta* | AC280286.1 | 5.00E-43 | 95.00% |
| 17P | 196 | *Macaca mulatta* | AC197276.3 | 6.00E-71 | 91.33% |
| 18P | 140 | *Macaca mulatta* | AC202605.16 | 3.00E-53 | 95.00% |
| 19P | 250 | *Macaca mulatta* | AB128049.1 | 4.00E-94 | 91.67% |
| 20P | 121 | *Macaca nemestrina* | XR_996760.2 | 7.00E-42 | 92.24% |
| 21P | 71 | *Pan paniscus* | XM_034938079.2 | 1.00E-21 | 95.77% |
| 22P | 146 | *Pan paniscus* | AC280336.1 | 3.00E-66 | 100.00% |
| 23P | 141 | *Pan paniscus* | AC280342.1 | 2.00E-63 | 100.00% |
| 24P | 163 | *Pan paniscus* | XM_055115310.1 | 6.00E-70 | 97.55% |
| 25P | 148 | *Pan paniscus* | AC280341.1 | 3.00E-67 | 100.00% |
| 26P | 153 | *Pan paniscus* | XM_008974855.5 | 3.00E-67 | 98.71% |
| 27P | 173 | *Pan paniscus* | AC280344.1 | 4.00E-79 | 99.42% |
| 28P | 139 | *Pan paniscus* | XM034963660.2 | 1.00E-52 | 94.37% |
| 29P | 97 | *Gorilla gorilla* | AC270505.1 | 3.00E-38 | 98.97% |
| 30P | 97 | *Gorilla gorilla* | AC270505.1 | 5.00E-41 | 100.00% |
| 31P | 135 | *Gorilla gorilla* | AC275387.1 | 3.00E-60 | 99.26% |
| 32P | 103 | *Gorilla gorilla* | AC145355.3 | 2.00E-44 | 100.00% |
| 33P | 209 | *Gorilla gorilla* | AC145852.3 | 3.00E-81 | 93.78% |
| 34P | 163 | *Gorilla gorilla* | AC280153.1 | 9.00E-71 | 97.55% |
| 35P | 150 | *Gorilla gorilla* | AC275724.1 | 6.00E-67 | 98.67% |
| 36P | 114 | *Gorilla gorilla* | AC280262.1 | 1.00E-48 | 99.12% |

**Table S8.**
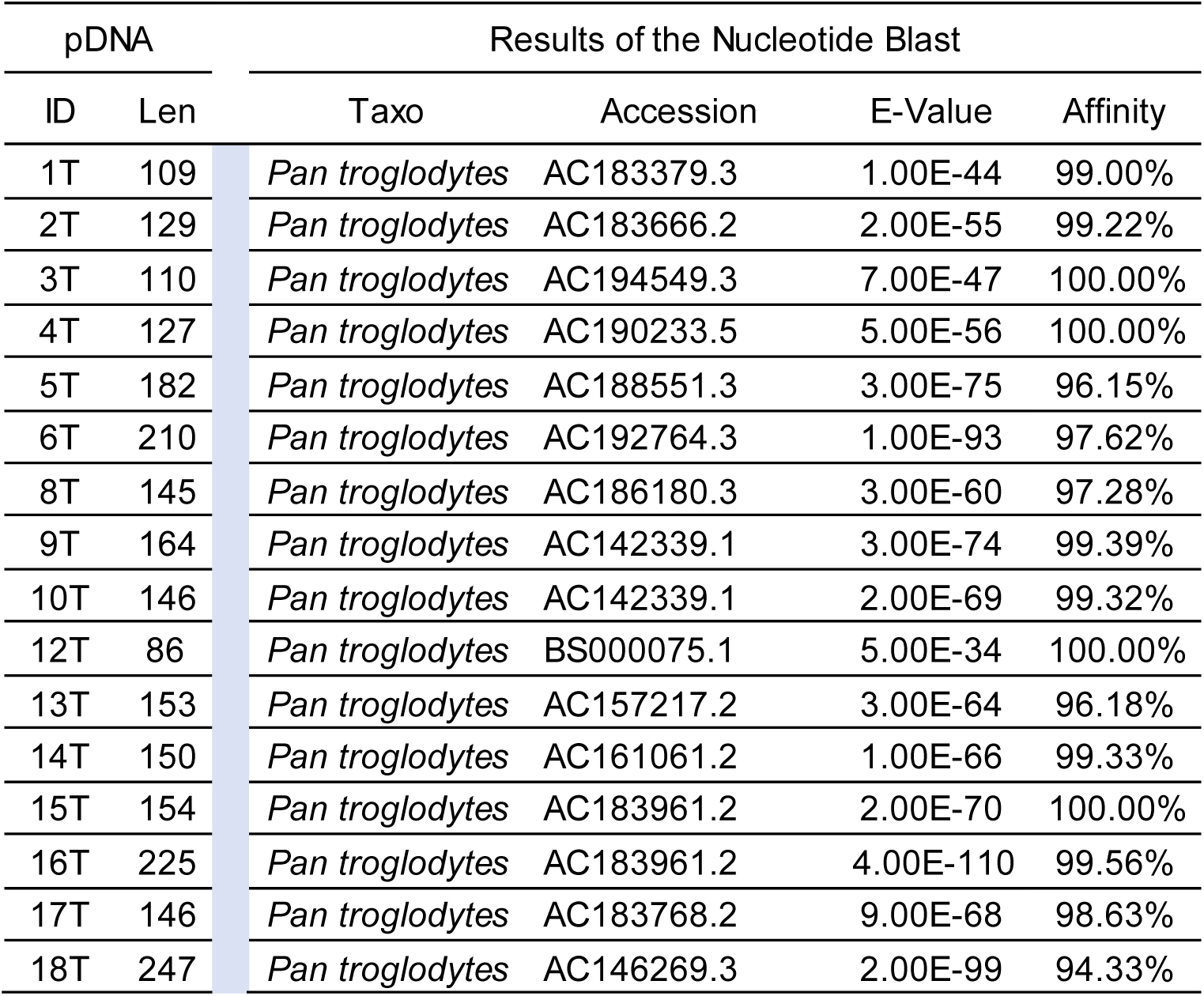
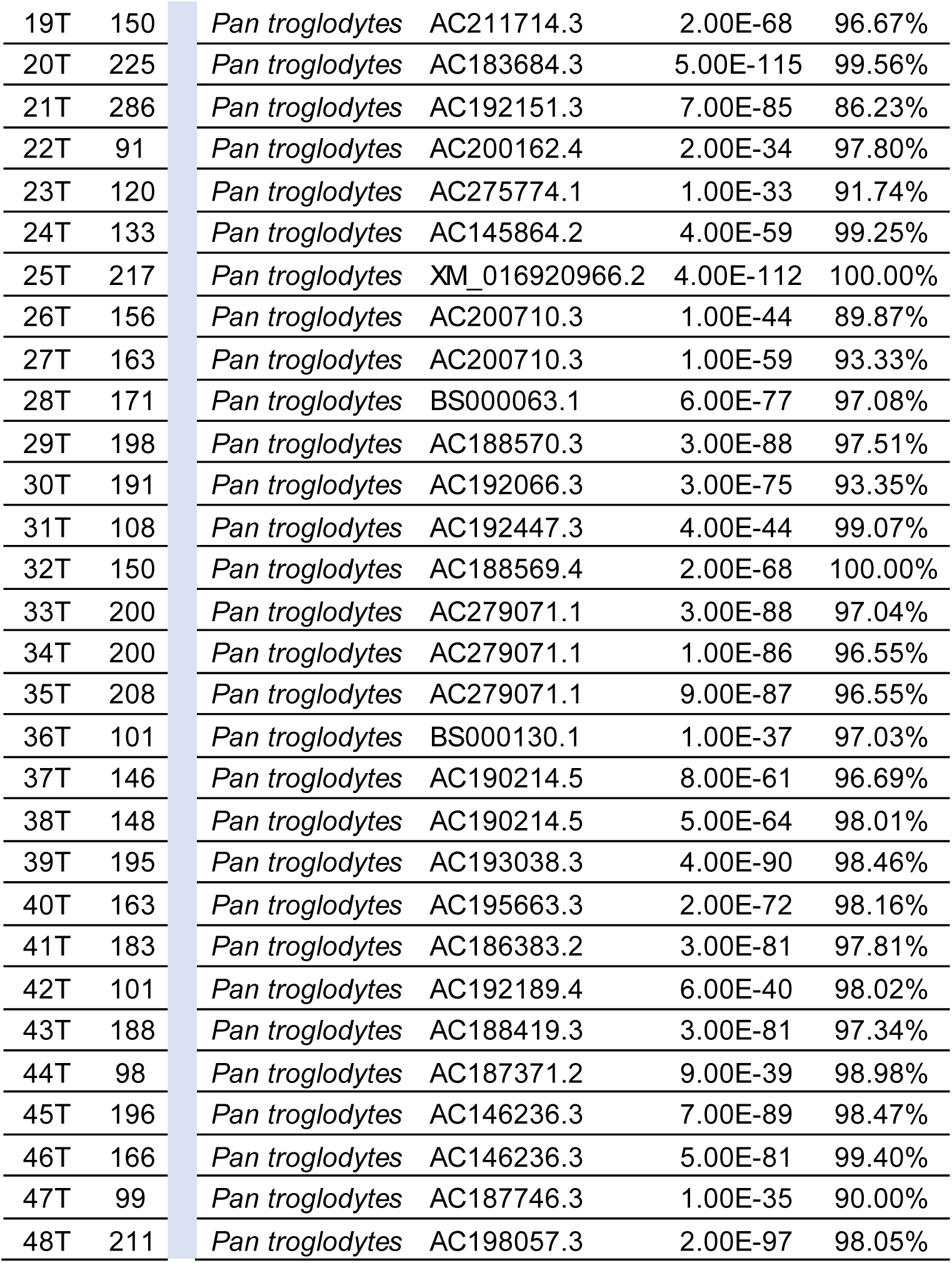
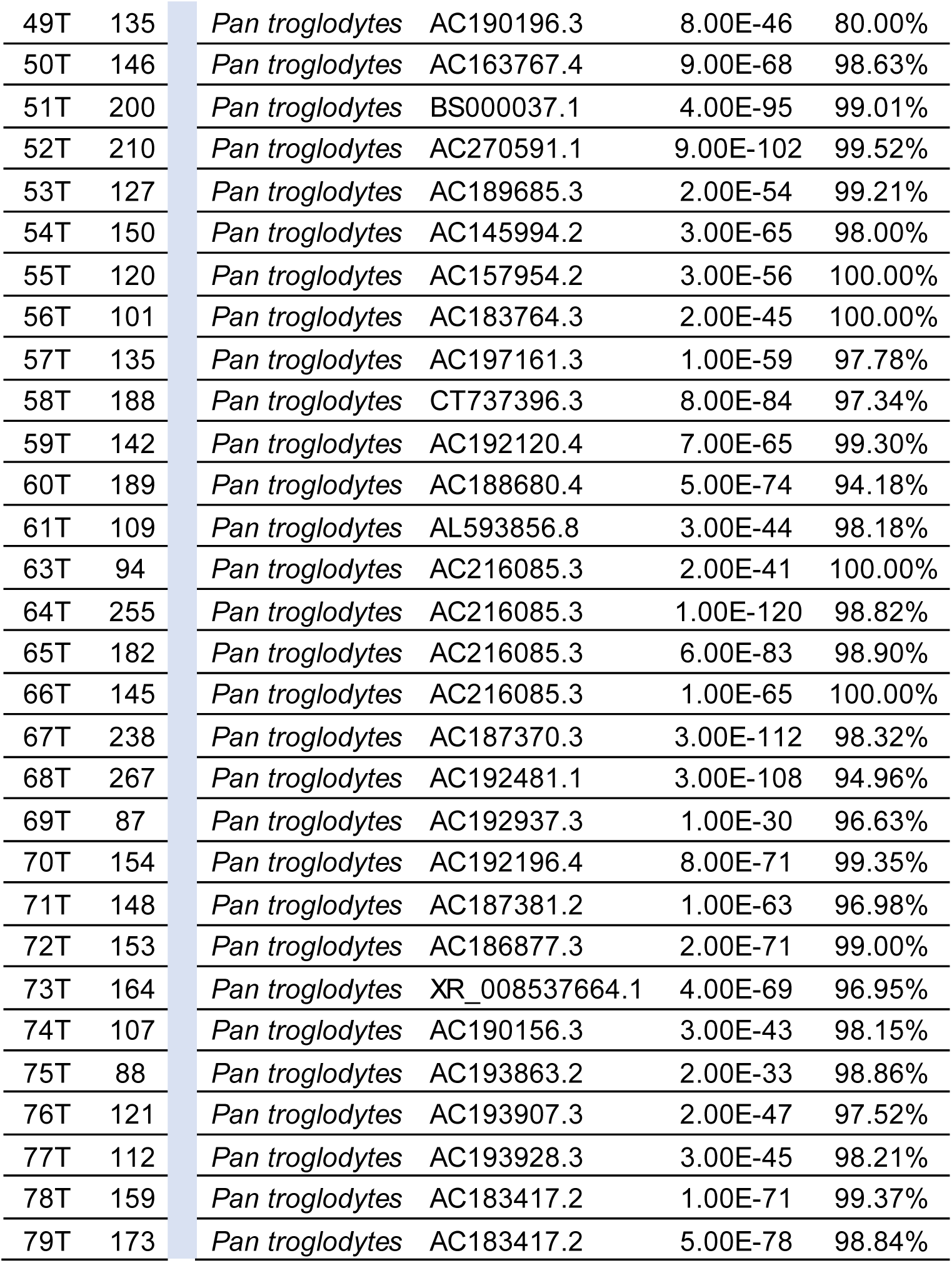
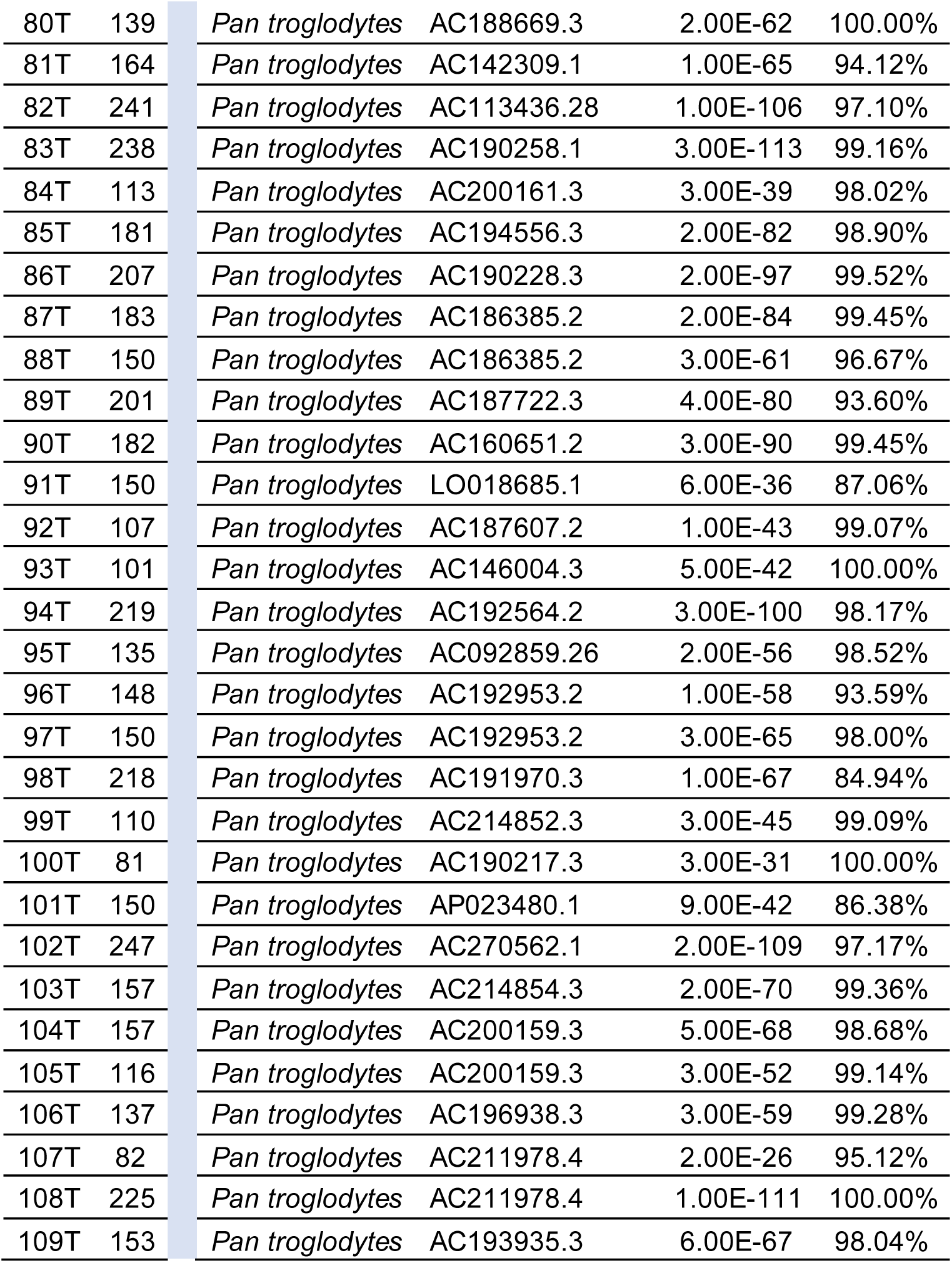
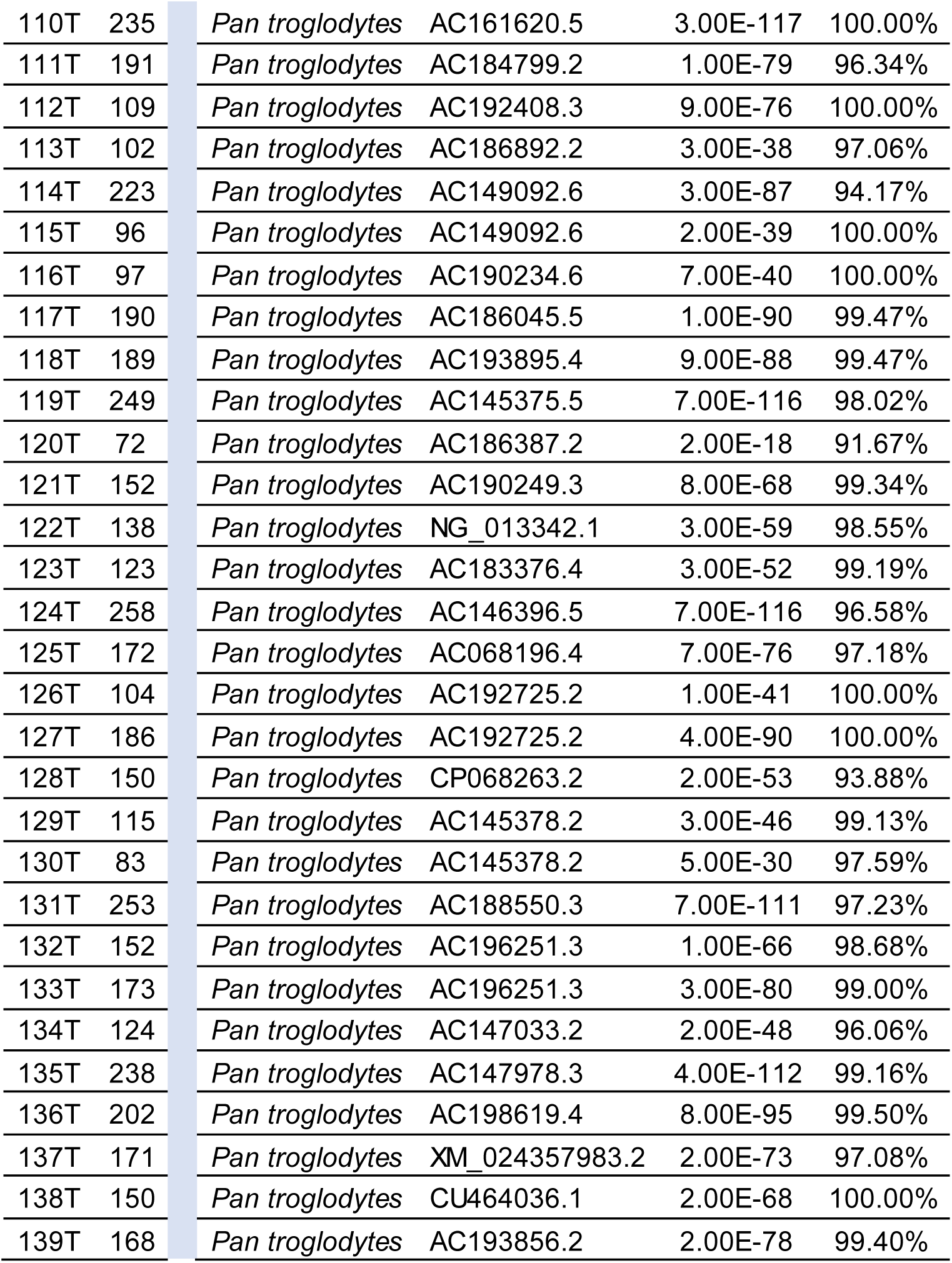
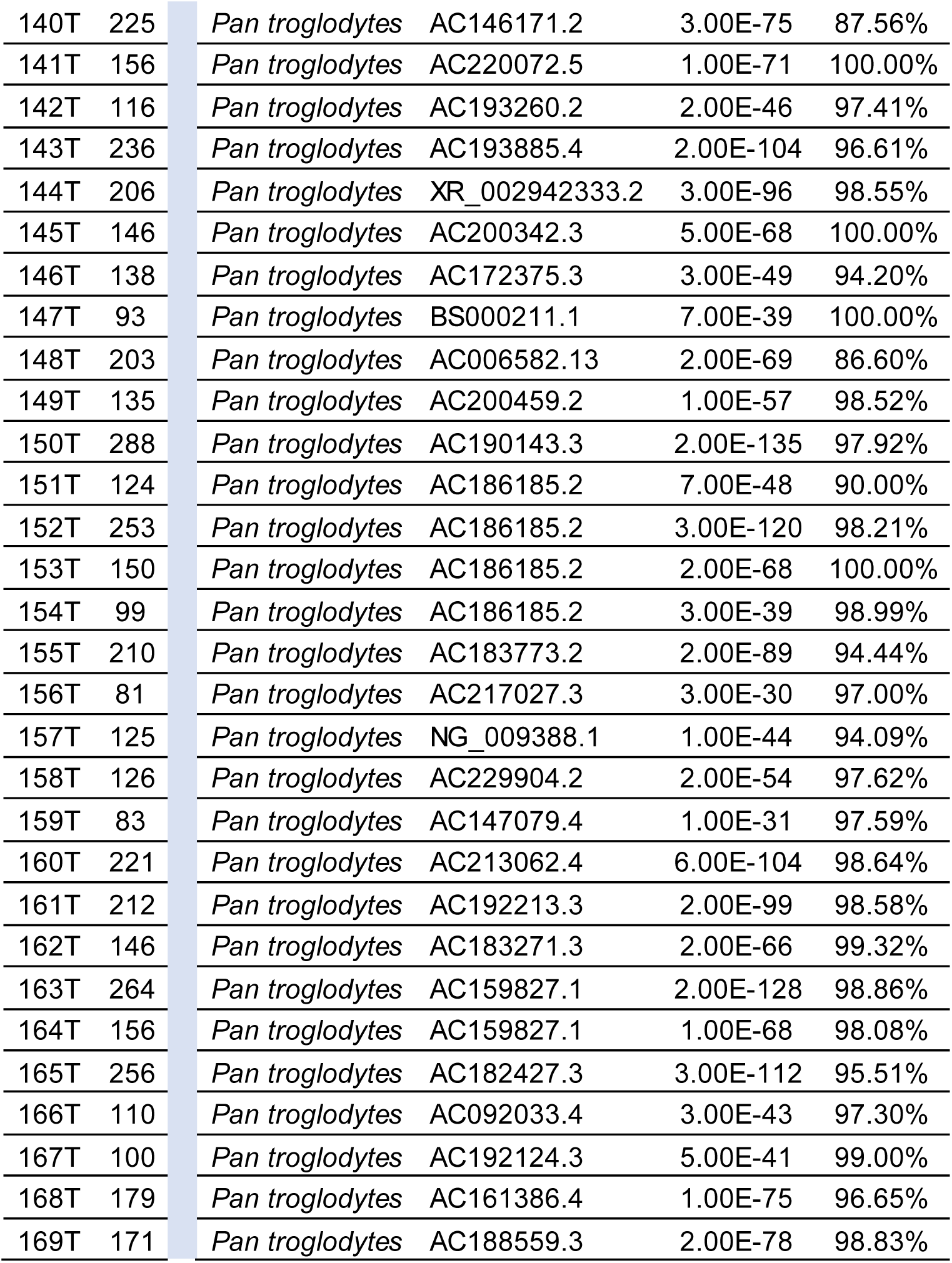
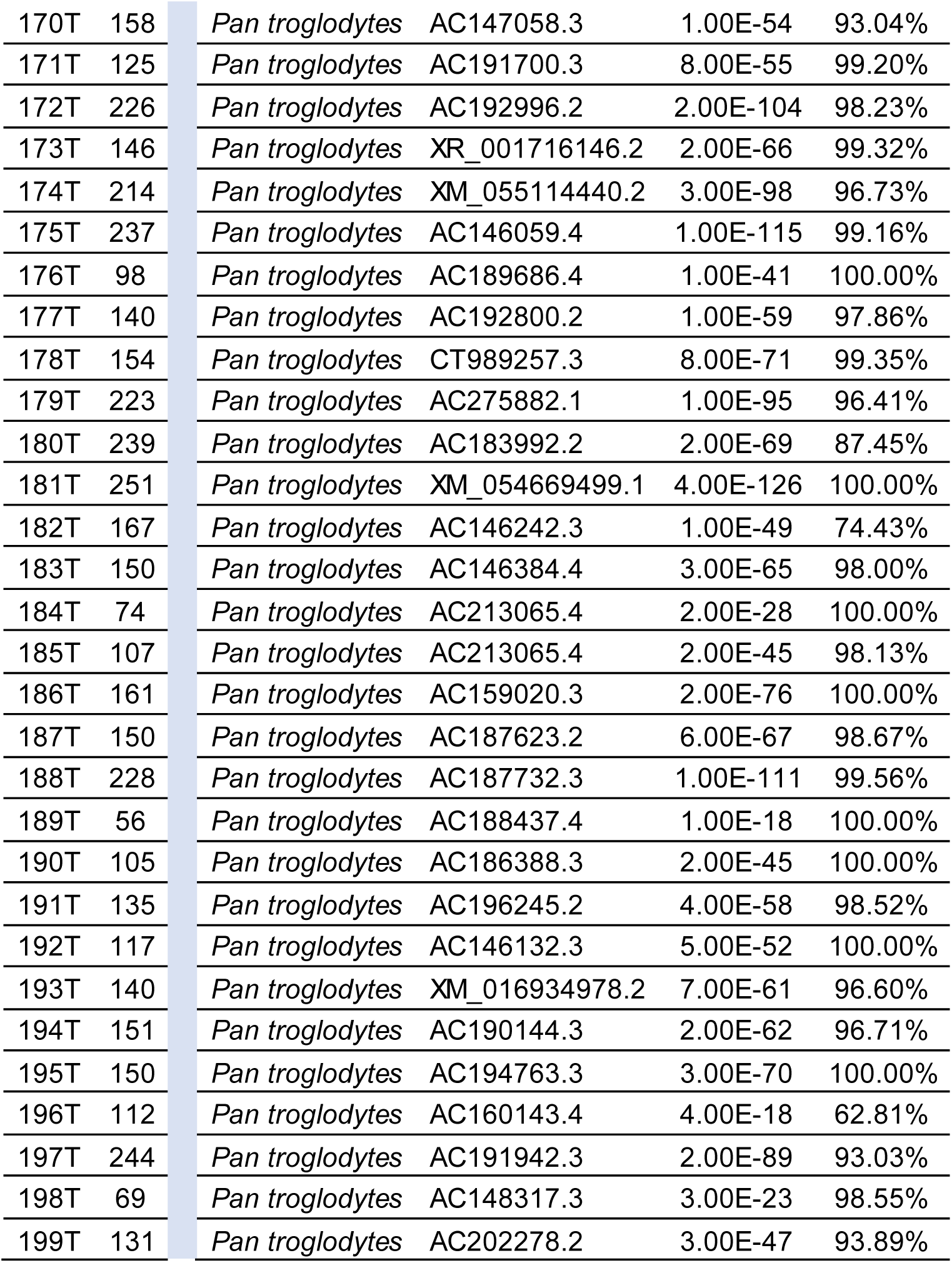
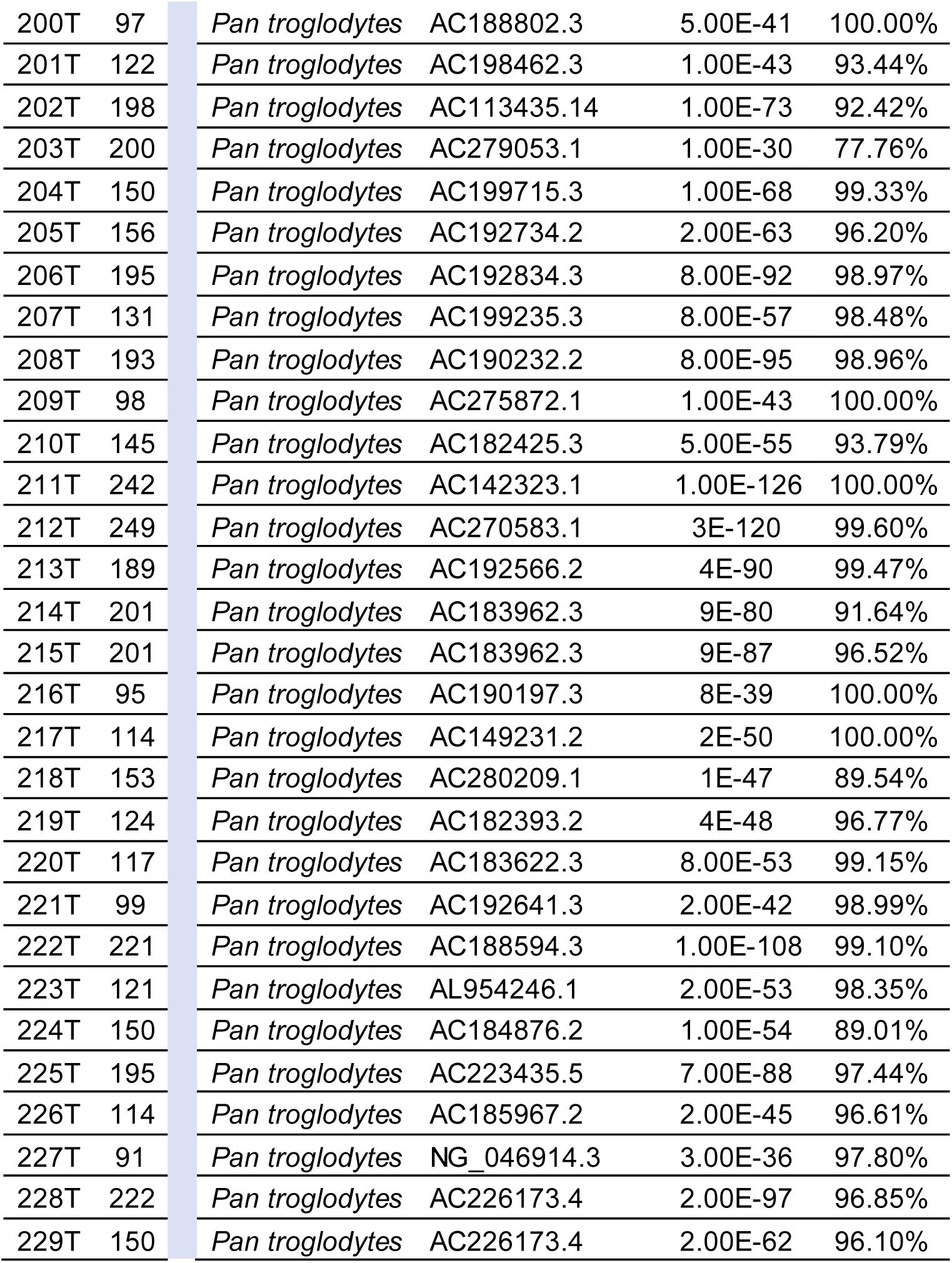
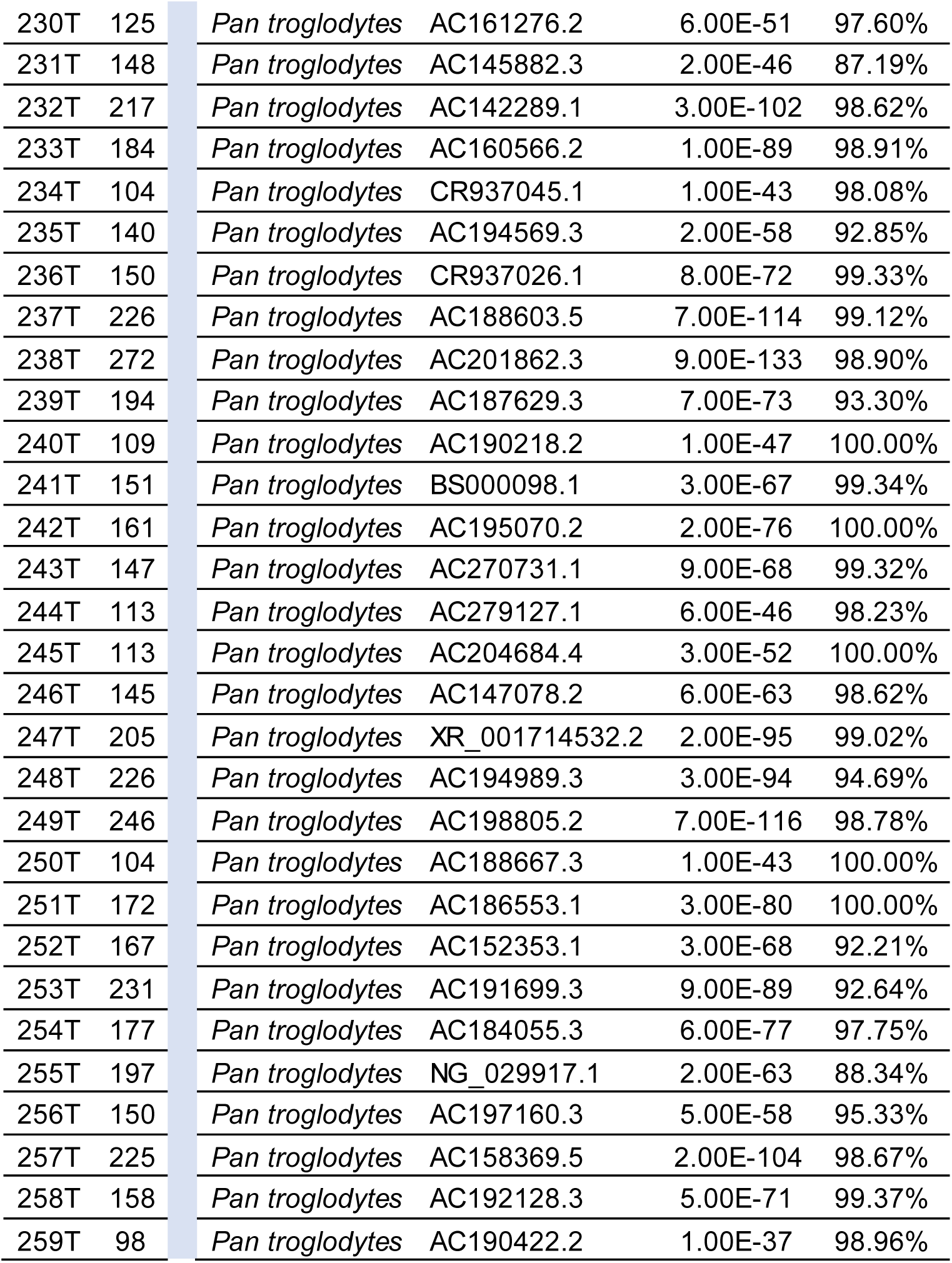
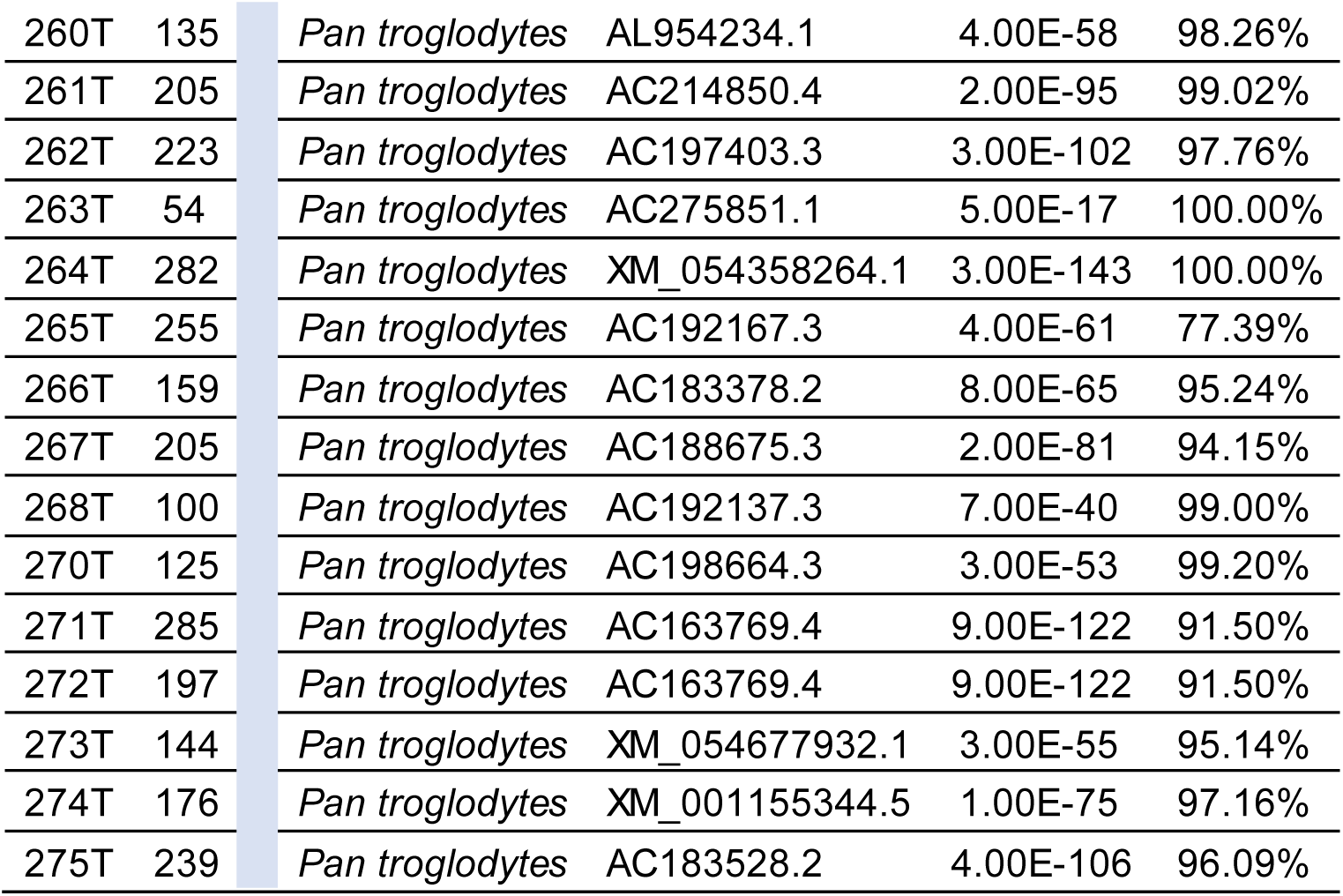
The 271 pDNA sequences matching the nuclear sequences of *Pan troglodyte*.

| pDNA |  | Results of the Nucleotide Blast |  |  |  |
| --- | --- | --- | --- | --- | --- |
| ID | Len | Taxo | Accession | E-Value | Affinity |
| 1T | 109 | *Pan troglodytes* | AC183379.3 | 1.00E-44 | 99.00% |
| 2T | 129 | *Pan troglodytes* | AC183666.2 | 2.00E-55 | 99.22% |
| 3T | 110 | *Pan troglodytes* | AC194549.3 | 7.00E-47 | 100.00% |
| 4T | 127 | *Pan troglodytes* | AC190233.5 | 5.00E-56 | 100.00% |
| 5T | 182 | *Pan troglodytes* | AC188551.3 | 3.00E-75 | 96.15% |
| 6T | 210 | *Pan troglodytes* | AC192764.3 | 1.00E-93 | 97.62% |
| 8T | 145 | *Pan troglodytes* | AC186180.3 | 3.00E-60 | 97.28% |
| 9T | 164 | *Pan troglodytes* | AC142339.1 | 3.00E-74 | 99.39% |
| 10T | 146 | *Pan troglodytes* | AC142339.1 | 2.00E-69 | 99.32% |
| 12T | 86 | *Pan troglodytes* | BS000075.1 | 5.00E-34 | 100.00% |
| 13T | 153 | *Pan troglodytes* | AC157217.2 | 3.00E-64 | 96.18% |
| 14T | 150 | *Pan troglodytes* | AC161061.2 | 1.00E-66 | 99.33% |
| 15T | 154 | *Pan troglodytes* | AC183961.2 | 2.00E-70 | 100.00% |
| 16T | 225 | *Pan troglodytes* | AC183961.2 | 4.00E-110 | 99.56% |
| 17T | 146 | *Pan troglodytes* | AC183768.2 | 9.00E-68 | 98.63% |
| 18T | 247 | *Pan troglodytes* | AC146269.3 | 2.00E-99 | 94.33% |
| 19T | 150 | *Pan troglodytes* | AC211714.3 | 2.00E-68 | 96.67% |
| 20T | 225 | *Pan troglodytes* | AC183684.3 | 5.00E-115 | 99.56% |
| 21T | 286 | *Pan troglodytes* | AC192151.3 | 7.00E-85 | 86.23% |
| 22T | 91 | *Pan troglodytes* | AC200162.4 | 2.00E-34 | 97.80% |
| 23T | 120 | *Pan troglodytes* | AC275774.1 | 1.00E-33 | 91.74% |
| 24T | 133 | *Pan troglodytes* | AC145864.2 | 4.00E-59 | 99.25% |
| 25T | 217 | *Pan troglodytes* | XM_016920966.2 | 4.00E-112 | 100.00% |
| 26T | 156 | *Pan troglodytes* | AC200710.3 | 1.00E-44 | 89.87% |
| 27T | 163 | *Pan troglodytes* | AC200710.3 | 1.00E-59 | 93.33% |
| 28T | 171 | *Pan troglodytes* | BS000063.1 | 6.00E-77 | 97.08% |
| 29T | 198 | *Pan troglodytes* | AC188570.3 | 3.00E-88 | 97.51% |
| 30T | 191 | *Pan troglodytes* | AC192066.3 | 3.00E-75 | 93.35% |
| 31T | 108 | *Pan troglodytes* | AC192447.3 | 4.00E-44 | 99.07% |
| 32T | 150 | *Pan troglodytes* | AC188569.4 | 2.00E-68 | 100.00% |
| 33T | 200 | *Pan troglodytes* | AC279071.1 | 3.00E-88 | 97.04% |
| 34T | 200 | *Pan troglodytes* | AC279071.1 | 1.00E-86 | 96.55% |
| 35T | 208 | *Pan troglodytes* | AC279071.1 | 9.00E-87 | 96.55% |
| 36T | 101 | *Pan troglodytes* | BS000130.1 | 1.00E-37 | 97.03% |
| 37T | 146 | *Pan troglodytes* | AC190214.5 | 8.00E-61 | 96.69% |
| 38T | 148 | *Pan troglodytes* | AC190214.5 | 5.00E-64 | 98.01% |
| 39T | 195 | *Pan troglodytes* | AC193038.3 | 4.00E-90 | 98.46% |
| 40T | 163 | *Pan troglodytes* | AC195663.3 | 2.00E-72 | 98.16% |
| 41T | 183 | *Pan troglodytes* | AC186383.2 | 3.00E-81 | 97.81% |
| 42T | 101 | *Pan troglodytes* | AC192189.4 | 6.00E-40 | 98.02% |
| 43T | 188 | *Pan troglodytes* | AC188419.3 | 3.00E-81 | 97.34% |
| 44T | 98 | *Pan troglodytes* | AC187371.2 | 9.00E-39 | 98.98% |
| 45T | 196 | *Pan troglodytes* | AC146236.3 | 7.00E-89 | 98.47% |
| 46T | 166 | *Pan troglodytes* | AC146236.3 | 5.00E-81 | 99.40% |
| 47T | 99 | *Pan troglodytes* | AC187746.3 | 1.00E-35 | 90.00% |
| 48T | 211 | *Pan troglodytes* | AC198057.3 | 2.00E-97 | 98.05% |
| 49T | 135 | *Pan troglodytes* | AC190196.3 | 8.00E-46 | 80.00% |
| 50T | 146 | *Pan troglodytes* | AC163767.4 | 9.00E-68 | 98.63% |
| 51T | 200 | *Pan troglodytes* | BS000037.1 | 4.00E-95 | 99.01% |
| 52T | 210 | *Pan troglodytes* | AC270591.1 | 9.00E-102 | 99.52% |
| 53T | 127 | *Pan troglodytes* | AC189685.3 | 2.00E-54 | 99.21% |
| 54T | 150 | *Pan troglodytes* | AC145994.2 | 3.00E-65 | 98.00% |
| 55T | 120 | *Pan troglodytes* | AC157954.2 | 3.00E-56 | 100.00% |
| 56T | 101 | *Pan troglodytes* | AC183764.3 | 2.00E-45 | 100.00% |
| 57T | 135 | *Pan troglodytes* | AC197161.3 | 1.00E-59 | 97.78% |
| 58T | 188 | *Pan troglodytes* | CT737396.3 | 8.00E-84 | 97.34% |
| 59T | 142 | *Pan troglodytes* | AC192120.4 | 7.00E-65 | 99.30% |
| 60T | 189 | *Pan troglodytes* | AC188680.4 | 5.00E-74 | 94.18% |
| 61T | 109 | *Pan troglodytes* | AL593856.8 | 3.00E-44 | 98.18% |
| 63T | 94 | *Pan troglodytes* | AC216085.3 | 2.00E-41 | 100.00% |
| 64T | 255 | *Pan troglodytes* | AC216085.3 | 1.00E-120 | 98.82% |
| 65T | 182 | *Pan troglodytes* | AC216085.3 | 6.00E-83 | 98.90% |
| 66T | 145 | *Pan troglodytes* | AC216085.3 | 1.00E-65 | 100.00% |
| 67T | 238 | *Pan troglodytes* | AC187370.3 | 3.00E-112 | 98.32% |
| 68T | 267 | *Pan troglodytes* | AC192481.1 | 3.00E-108 | 94.96% |
| 69T | 87 | *Pan troglodytes* | AC192937.3 | 1.00E-30 | 96.63% |
| 70T | 154 | *Pan troglodytes* | AC192196.4 | 8.00E-71 | 99.35% |
| 71T | 148 | *Pan troglodytes* | AC187381.2 | 1.00E-63 | 96.98% |
| 72T | 153 | *Pan troglodytes* | AC186877.3 | 2.00E-71 | 99.00% |
| 73T | 164 | *Pan troglodytes* | XR_008537664.1 | 4.00E-69 | 96.95% |
| 74T | 107 | *Pan troglodytes* | AC190156.3 | 3.00E-43 | 98.15% |
| 75T | 88 | *Pan troglodytes* | AC193863.2 | 2.00E-33 | 98.86% |
| 76T | 121 | *Pan troglodytes* | AC193907.3 | 2.00E-47 | 97.52% |
| 77T | 112 | *Pan troglodytes* | AC193928.3 | 3.00E-45 | 98.21% |
| 78T | 159 | *Pan troglodytes* | AC183417.2 | 1.00E-71 | 99.37% |
| 79T | 173 | *Pan troglodytes* | AC183417.2 | 5.00E-78 | 98.84% |
| 80T | 139 | *Pan troglodytes* | AC188669.3 | 2.00E-62 | 100.00% |
| 81T | 164 | *Pan troglodytes* | AC142309.1 | 1.00E-65 | 94.12% |
| 82T | 241 | *Pan troglodytes* | AC113436.28 | 1.00E-106 | 97.10% |
| 83T | 238 | *Pan troglodytes* | AC190258.1 | 3.00E-113 | 99.16% |
| 84T | 113 | *Pan troglodytes* | AC200161.3 | 3.00E-39 | 98.02% |
| 85T | 181 | *Pan troglodytes* | AC194556.3 | 2.00E-82 | 98.90% |
| 86T | 207 | *Pan troglodytes* | AC190228.3 | 2.00E-97 | 99.52% |
| 87T | 183 | *Pan troglodytes* | AC186385.2 | 2.00E-84 | 99.45% |
| 88T | 150 | *Pan troglodytes* | AC186385.2 | 3.00E-61 | 96.67% |
| 89T | 201 | *Pan troglodytes* | AC187722.3 | 4.00E-80 | 93.60% |
| 90T | 182 | *Pan troglodytes* | AC160651.2 | 3.00E-90 | 99.45% |
| 91T | 150 | *Pan troglodytes* | LO018685.1 | 6.00E-36 | 87.06% |
| 92T | 107 | *Pan troglodytes* | AC187607.2 | 1.00E-43 | 99.07% |
| 93T | 101 | *Pan troglodytes* | AC146004.3 | 5.00E-42 | 100.00% |
| 94T | 219 | *Pan troglodytes* | AC192564.2 | 3.00E-100 | 98.17% |
| 95T | 135 | *Pan troglodytes* | AC092859.26 | 2.00E-56 | 98.52% |
| 96T | 148 | *Pan troglodytes* | AC192953.2 | 1.00E-58 | 93.59% |
| 97T | 150 | *Pan troglodytes* | AC192953.2 | 3.00E-65 | 98.00% |
| 98T | 218 | *Pan troglodytes* | AC191970.3 | 1.00E-67 | 84.94% |
| 99T | 110 | *Pan troglodytes* | AC214852.3 | 3.00E-45 | 99.09% |
| 100T | 81 | *Pan troglodytes* | AC190217.3 | 3.00E-31 | 100.00% |
| 101T | 150 | *Pan troglodytes* | AP023480.1 | 9.00E-42 | 86.38% |
| 102T | 247 | *Pan troglodytes* | AC270562.1 | 2.00E-109 | 97.17% |
| 103T | 157 | *Pan troglodytes* | AC214854.3 | 2.00E-70 | 99.36% |
| 104T | 157 | *Pan troglodytes* | AC200159.3 | 5.00E-68 | 98.68% |
| 105T | 116 | *Pan troglodytes* | AC200159.3 | 3.00E-52 | 99.14% |
| 106T | 137 | *Pan troglodytes* | AC196938.3 | 3.00E-59 | 99.28% |
| 107T | 82 | *Pan troglodytes* | AC211978.4 | 2.00E-26 | 95.12% |
| 108T | 225 | *Pan troglodytes* | AC211978.4 | 1.00E-111 | 100.00% |
| 109T | 153 | *Pan troglodytes* | AC193935.3 | 6.00E-67 | 98.04% |
| 110T | 235 | *Pan troglodytes* | AC161620.5 | 3.00E-117 | 100.00% |
| 111T | 191 | *Pan troglodytes* | AC184799.2 | 1.00E-79 | 96.34% |
| 112T | 109 | *Pan troglodytes* | AC192408.3 | 9.00E-76 | 100.00% |
| 113T | 102 | *Pan troglodytes* | AC186892.2 | 3.00E-38 | 97.06% |
| 114T | 223 | *Pan troglodytes* | AC149092.6 | 3.00E-87 | 94.17% |
| 115T | 96 | *Pan troglodytes* | AC149092.6 | 2.00E-39 | 100.00% |
| 116T | 97 | *Pan troglodytes* | AC190234.6 | 7.00E-40 | 100.00% |
| 117T | 190 | *Pan troglodytes* | AC186045.5 | 1.00E-90 | 99.47% |
| 118T | 189 | *Pan troglodytes* | AC193895.4 | 9.00E-88 | 99.47% |
| 119T | 249 | *Pan troglodytes* | AC145375.5 | 7.00E-116 | 98.02% |
| 120T | 72 | *Pan troglodytes* | AC186387.2 | 2.00E-18 | 91.67% |
| 121T | 152 | *Pan troglodytes* | AC190249.3 | 8.00E-68 | 99.34% |
| 122T | 138 | *Pan troglodytes* | NG_013342.1 | 3.00E-59 | 98.55% |
| 123T | 123 | *Pan troglodytes* | AC183376.4 | 3.00E-52 | 99.19% |
| 124T | 258 | *Pan troglodytes* | AC146396.5 | 7.00E-116 | 96.58% |
| 125T | 172 | *Pan troglodytes* | AC068196.4 | 7.00E-76 | 97.18% |
| 126T | 104 | *Pan troglodytes* | AC192725.2 | 1.00E-41 | 100.00% |
| 127T | 186 | *Pan troglodytes* | AC192725.2 | 4.00E-90 | 100.00% |
| 128T | 150 | *Pan troglodytes* | CP068263.2 | 2.00E-53 | 93.88% |
| 129T | 115 | *Pan troglodytes* | AC145378.2 | 3.00E-46 | 99.13% |
| 130T | 83 | *Pan troglodytes* | AC145378.2 | 5.00E-30 | 97.59% |
| 131T | 253 | *Pan troglodytes* | AC188550.3 | 7.00E-111 | 97.23% |
| 132T | 152 | *Pan troglodytes* | AC196251.3 | 1.00E-66 | 98.68% |
| 133T | 173 | *Pan troglodytes* | AC196251.3 | 3.00E-80 | 99.00% |
| 134T | 124 | *Pan troglodytes* | AC147033.2 | 2.00E-48 | 96.06% |
| 135T | 238 | *Pan troglodytes* | AC147978.3 | 4.00E-112 | 99.16% |
| 136T | 202 | *Pan troglodytes* | AC198619.4 | 8.00E-95 | 99.50% |
| 137T | 171 | *Pan troglodytes* | XM_024357983.2 | 2.00E-73 | 97.08% |
| 138T | 150 | *Pan troglodytes* | CU464036.1 | 2.00E-68 | 100.00% |
| 139T | 168 | *Pan troglodytes* | AC193856.2 | 2.00E-78 | 99.40% |
| 140T | 225 | *Pan troglodytes* | AC146171.2 | 3.00E-75 | 87.56% |
| 141T | 156 | *Pan troglodytes* | AC220072.5 | 1.00E-71 | 100.00% |
| 142T | 116 | *Pan troglodytes* | AC193260.2 | 2.00E-46 | 97.41% |
| 143T | 236 | *Pan troglodytes* | AC193885.4 | 2.00E-104 | 96.61% |
| 144T | 206 | *Pan troglodytes* | XR_002942333.2 | 3.00E-96 | 98.55% |
| 145T | 146 | *Pan troglodytes* | AC200342.3 | 5.00E-68 | 100.00% |
| 146T | 138 | *Pan troglodytes* | AC172375.3 | 3.00E-49 | 94.20% |
| 147T | 93 | *Pan troglodytes* | BS000211.1 | 7.00E-39 | 100.00% |
| 148T | 203 | *Pan troglodytes* | AC006582.13 | 2.00E-69 | 86.60% |
| 149T | 135 | *Pan troglodytes* | AC200459.2 | 1.00E-57 | 98.52% |
| 150T | 288 | *Pan troglodytes* | AC190143.3 | 2.00E-135 | 97.92% |
| 151T | 124 | *Pan troglodytes* | AC186185.2 | 7.00E-48 | 90.00% |
| 152T | 253 | *Pan troglodytes* | AC186185.2 | 3.00E-120 | 98.21% |
| 153T | 150 | *Pan troglodytes* | AC186185.2 | 2.00E-68 | 100.00% |
| 154T | 99 | *Pan troglodytes* | AC186185.2 | 3.00E-39 | 98.99% |
| 155T | 210 | *Pan troglodytes* | AC183773.2 | 2.00E-89 | 94.44% |
| 156T | 81 | *Pan troglodytes* | AC217027.3 | 3.00E-30 | 97.00% |
| 157T | 125 | *Pan troglodytes* | NG_009388.1 | 1.00E-44 | 94.09% |
| 158T | 126 | *Pan troglodytes* | AC229904.2 | 2.00E-54 | 97.62% |
| 159T | 83 | *Pan troglodytes* | AC147079.4 | 1.00E-31 | 97.59% |
| 160T | 221 | *Pan troglodytes* | AC213062.4 | 6.00E-104 | 98.64% |
| 161T | 212 | *Pan troglodytes* | AC192213.3 | 2.00E-99 | 98.58% |
| 162T | 146 | *Pan troglodytes* | AC183271.3 | 2.00E-66 | 99.32% |
| 163T | 264 | *Pan troglodytes* | AC159827.1 | 2.00E-128 | 98.86% |
| 164T | 156 | *Pan troglodytes* | AC159827.1 | 1.00E-68 | 98.08% |
| 165T | 256 | *Pan troglodytes* | AC182427.3 | 3.00E-112 | 95.51% |
| 166T | 110 | *Pan troglodytes* | AC092033.4 | 3.00E-43 | 97.30% |
| 167T | 100 | *Pan troglodytes* | AC192124.3 | 5.00E-41 | 99.00% |
| 168T | 179 | *Pan troglodytes* | AC161386.4 | 1.00E-75 | 96.65% |
| 169T | 171 | *Pan troglodytes* | AC188559.3 | 2.00E-78 | 98.83% |
| 170T | 158 | *Pan troglodytes* | AC147058.3 | 1.00E-54 | 93.04% |
| 171T | 125 | *Pan troglodytes* | AC191700.3 | 8.00E-55 | 99.20% |
| 172T | 226 | *Pan troglodytes* | AC192996.2 | 2.00E-104 | 98.23% |
| 173T | 146 | *Pan troglodytes* | XR_001716146.2 | 2.00E-66 | 99.32% |
| 174T | 214 | *Pan troglodytes* | XM_055114440.2 | 3.00E-98 | 96.73% |
| 175T | 237 | *Pan troglodytes* | AC146059.4 | 1.00E-115 | 99.16% |
| 176T | 98 | *Pan troglodytes* | AC189686.4 | 1.00E-41 | 100.00% |
| 177T | 140 | *Pan troglodytes* | AC192800.2 | 1.00E-59 | 97.86% |
| 178T | 154 | *Pan troglodytes* | CT989257.3 | 8.00E-71 | 99.35% |
| 179T | 223 | *Pan troglodytes* | AC275882.1 | 1.00E-95 | 96.41% |
| 180T | 239 | *Pan troglodytes* | AC183992.2 | 2.00E-69 | 87.45% |
| 181T | 251 | *Pan troglodytes* | XM_054669499.1 | 4.00E-126 | 100.00% |
| 182T | 167 | *Pan troglodytes* | AC146242.3 | 1.00E-49 | 74.43% |
| 183T | 150 | *Pan troglodytes* | AC146384.4 | 3.00E-65 | 98.00% |
| 184T | 74 | *Pan troglodytes* | AC213065.4 | 2.00E-28 | 100.00% |
| 185T | 107 | *Pan troglodytes* | AC213065.4 | 2.00E-45 | 98.13% |
| 186T | 161 | *Pan troglodytes* | AC159020.3 | 2.00E-76 | 100.00% |
| 187T | 150 | *Pan troglodytes* | AC187623.2 | 6.00E-67 | 98.67% |
| 188T | 228 | *Pan troglodytes* | AC187732.3 | 1.00E-111 | 99.56% |
| 189T | 56 | *Pan troglodytes* | AC188437.4 | 1.00E-18 | 100.00% |
| 190T | 105 | *Pan troglodytes* | AC186388.3 | 2.00E-45 | 100.00% |
| 191T | 135 | *Pan troglodytes* | AC196245.2 | 4.00E-58 | 98.52% |
| 192T | 117 | *Pan troglodytes* | AC146132.3 | 5.00E-52 | 100.00% |
| 193T | 140 | *Pan troglodytes* | XM_016934978.2 | 7.00E-61 | 96.60% |
| 194T | 151 | *Pan troglodytes* | AC190144.3 | 2.00E-62 | 96.71% |
| 195T | 150 | *Pan troglodytes* | AC194763.3 | 3.00E-70 | 100.00% |
| 196T | 112 | *Pan troglodytes* | AC160143.4 | 4.00E-18 | 62.81% |
| 197T | 244 | *Pan troglodytes* | AC191942.3 | 2.00E-89 | 93.03% |
| 198T | 69 | *Pan troglodytes* | AC148317.3 | 3.00E-23 | 98.55% |
| 199T | 131 | *Pan troglodytes* | AC202278.2 | 3.00E-47 | 93.89% |
| 200T | 97 | *Pan troglodytes* | AC188802.3 | 5.00E-41 | 100.00% |
| 201T | 122 | *Pan troglodytes* | AC198462.3 | 1.00E-43 | 93.44% |
| 202T | 198 | *Pan troglodytes* | AC113435.14 | 1.00E-73 | 92.42% |
| 203T | 200 | *Pan troglodytes* | AC279053.1 | 1.00E-30 | 77.76% |
| 204T | 150 | *Pan troglodytes* | AC199715.3 | 1.00E-68 | 99.33% |
| 205T | 156 | *Pan troglodytes* | AC192734.2 | 2.00E-63 | 96.20% |
| 206T | 195 | *Pan troglodytes* | AC192834.3 | 8.00E-92 | 98.97% |
| 207T | 131 | *Pan troglodytes* | AC199235.3 | 8.00E-57 | 98.48% |
| 208T | 193 | *Pan troglodytes* | AC190232.2 | 8.00E-95 | 98.96% |
| 209T | 98 | *Pan troglodytes* | AC275872.1 | 1.00E-43 | 100.00% |
| 210T | 145 | *Pan troglodytes* | AC182425.3 | 5.00E-55 | 93.79% |
| 211T | 242 | *Pan troglodytes* | AC142323.1 | 1.00E-126 | 100.00% |
| 212T | 249 | *Pan troglodytes* | AC270583.1 | 3E-120 | 99.60% |
| 213T | 189 | *Pan troglodytes* | AC192566.2 | 4E-90 | 99.47% |
| 214T | 201 | *Pan troglodytes* | AC183962.3 | 9E-80 | 91.64% |
| 215T | 201 | *Pan troglodytes* | AC183962.3 | 9E-87 | 96.52% |
| 216T | 95 | *Pan troglodytes* | AC190197.3 | 8E-39 | 100.00% |
| 217T | 114 | *Pan troglodytes* | AC149231.2 | 2E-50 | 100.00% |
| 218T | 153 | *Pan troglodytes* | AC280209.1 | 1E-47 | 89.54% |
| 219T | 124 | *Pan troglodytes* | AC182393.2 | 4E-48 | 96.77% |
| 220T | 117 | *Pan troglodytes* | AC183622.3 | 8.00E-53 | 99.15% |
| 221T | 99 | *Pan troglodytes* | AC192641.3 | 2.00E-42 | 98.99% |
| 222T | 221 | *Pan troglodytes* | AC188594.3 | 1.00E-108 | 99.10% |
| 223T | 121 | *Pan troglodytes* | AL954246.1 | 2.00E-53 | 98.35% |
| 224T | 150 | *Pan troglodytes* | AC184876.2 | 1.00E-54 | 89.01% |
| 225T | 195 | *Pan troglodytes* | AC223435.5 | 7.00E-88 | 97.44% |
| 226T | 114 | *Pan troglodytes* | AC185967.2 | 2.00E-45 | 96.61% |
| 227T | 91 | *Pan troglodytes* | NG_046914.3 | 3.00E-36 | 97.80% |
| 228T | 222 | *Pan troglodytes* | AC226173.4 | 2.00E-97 | 96.85% |
| 229T | 150 | *Pan troglodytes* | AC226173.4 | 2.00E-62 | 96.10% |
| 230T | 125 | *Pan troglodytes* | AC161276.2 | 6.00E-51 | 97.60% |
| 231T | 148 | *Pan troglodytes* | AC145882.3 | 2.00E-46 | 87.19% |
| 232T | 217 | *Pan troglodytes* | AC142289.1 | 3.00E-102 | 98.62% |
| 233T | 184 | *Pan troglodytes* | AC160566.2 | 1.00E-89 | 98.91% |
| 234T | 104 | *Pan troglodytes* | CR937045.1 | 1.00E-43 | 98.08% |
| 235T | 140 | *Pan troglodytes* | AC194569.3 | 2.00E-58 | 92.85% |
| 236T | 150 | *Pan troglodytes* | CR937026.1 | 8.00E-72 | 99.33% |
| 237T | 226 | *Pan troglodytes* | AC188603.5 | 7.00E-114 | 99.12% |
| 238T | 272 | *Pan troglodytes* | AC201862.3 | 9.00E-133 | 98.90% |
| 239T | 194 | *Pan troglodytes* | AC187629.3 | 7.00E-73 | 93.30% |
| 240T | 109 | *Pan troglodytes* | AC190218.2 | 1.00E-47 | 100.00% |
| 241T | 151 | *Pan troglodytes* | BS000098.1 | 3.00E-67 | 99.34% |
| 242T | 161 | *Pan troglodytes* | AC195070.2 | 2.00E-76 | 100.00% |
| 243T | 147 | *Pan troglodytes* | AC270731.1 | 9.00E-68 | 99.32% |
| 244T | 113 | *Pan troglodytes* | AC279127.1 | 6.00E-46 | 98.23% |
| 245T | 113 | *Pan troglodytes* | AC204684.4 | 3.00E-52 | 100.00% |
| 246T | 145 | *Pan troglodytes* | AC147078.2 | 6.00E-63 | 98.62% |
| 247T | 205 | *Pan troglodytes* | XR_001714532.2 | 2.00E-95 | 99.02% |
| 248T | 226 | *Pan troglodytes* | AC194989.3 | 3.00E-94 | 94.69% |
| 249T | 246 | *Pan troglodytes* | AC198805.2 | 7.00E-116 | 98.78% |
| 250T | 104 | *Pan troglodytes* | AC188667.3 | 1.00E-43 | 100.00% |
| 251T | 172 | *Pan troglodytes* | AC186553.1 | 3.00E-80 | 100.00% |
| 252T | 167 | *Pan troglodytes* | AC152353.1 | 3.00E-68 | 92.21% |
| 253T | 231 | *Pan troglodytes* | AC191699.3 | 9.00E-89 | 92.64% |
| 254T | 177 | *Pan troglodytes* | AC184055.3 | 6.00E-77 | 97.75% |
| 255T | 197 | *Pan troglodytes* | NG_029917.1 | 2.00E-63 | 88.34% |
| 256T | 150 | *Pan troglodytes* | AC197160.3 | 5.00E-58 | 95.33% |
| 257T | 225 | *Pan troglodytes* | AC158369.5 | 2.00E-104 | 98.67% |
| 258T | 158 | *Pan troglodytes* | AC192128.3 | 5.00E-71 | 99.37% |
| 259T | 98 | *Pan troglodytes* | AC190422.2 | 1.00E-37 | 98.96% |
| 260T | 135 | *Pan troglodytes* | AL954234.1 | 4.00E-58 | 98.26% |
| 261T | 205 | *Pan troglodytes* | AC214850.4 | 2.00E-95 | 99.02% |
| 262T | 223 | *Pan troglodytes* | AC197403.3 | 3.00E-102 | 97.76% |
| 263T | 54 | *Pan troglodytes* | AC275851.1 | 5.00E-17 | 100.00% |
| 264T | 282 | *Pan troglodytes* | XM_054358264.1 | 3.00E-143 | 100.00% |
| 265T | 255 | *Pan troglodytes* | AC192167.3 | 4.00E-61 | 77.39% |
| 266T | 159 | *Pan troglodytes* | AC183378.2 | 8.00E-65 | 95.24% |
| 267T | 205 | *Pan troglodytes* | AC188675.3 | 2.00E-81 | 94.15% |
| 268T | 100 | *Pan troglodytes* | AC192137.3 | 7.00E-40 | 99.00% |
| 270T | 125 | *Pan troglodytes* | AC198664.3 | 3.00E-53 | 99.20% |
| 271T | 285 | *Pan troglodytes* | AC163769.4 | 9.00E-122 | 91.50% |
| 272T | 197 | *Pan troglodytes* | AC163769.4 | 9.00E-122 | 91.50% |
| 273T | 144 | *Pan troglodytes* | XM_054677932.1 | 3.00E-55 | 95.14% |
| 274T | 176 | *Pan troglodytes* | XM_001155344.5 | 1.00E-75 | 97.16% |
| 275T | 239 | *Pan troglodytes* | AC183528.2 | 4.00E-106 | 96.09% |

**Table S9.**
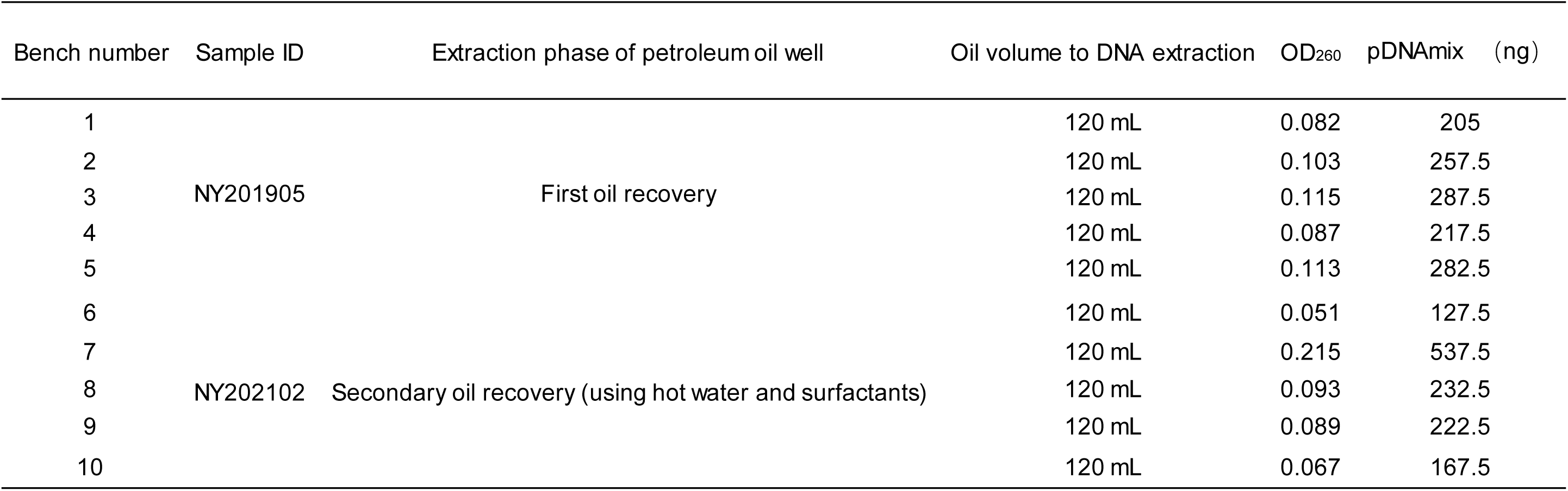
Quantification of DNA extraction for petroleum samples.

**Table S10.** Results of DNA extraction from petroleum and the illumina NovaSeq sequencing.

| Bench ID | Sample ID | Extraction phase of petroleum oil well | Oil volume to DNA extraction | OD<sub>260</sub> | pDNAMix (ng) | Total reads amount | Sequences from non-indigenous species |
| --- | --- | --- | --- | --- | --- | --- | --- |
| 1 | NY201905 | First oil extraction | 720 mL | 0.618 | 3708 | 3,159,002 | Multiple |
| 2 | NY202102 | Secondary oil recovery (using hot water and surfactants) | 720 mL | 0.675 | 4050 | 390,932 | None |

**Table S11.**
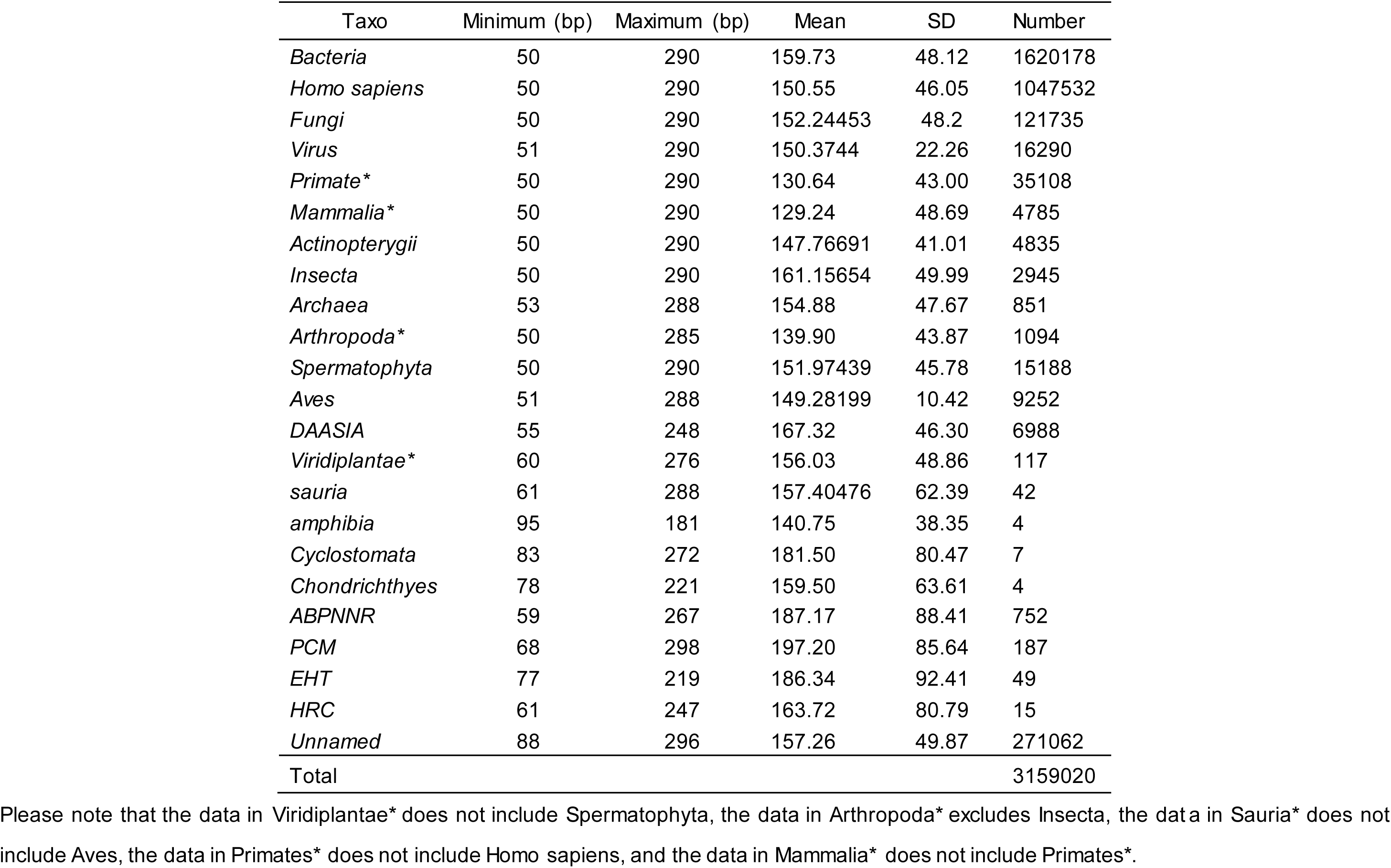
The length of pDNAs and the Blast results.

| Taxo | Minimum (bp) | Maximum (bp) | Mean | SD | Number |
| --- | --- | --- | --- | --- | --- |
| *Bacteria* | 50 | 290 | 159.73 | 48.12 | 1620178 |
| *Homo sapiens* | 50 | 290 | 150.55 | 46.05 | 1047532 |
| *Fungi* | 50 | 290 | 152.24453 | 48.2 | 121735 |
| *Virus* | 51 | 290 | 150.3744 | 22.26 | 16290 |
| *Primate\** | 50 | 290 | 130.64 | 43.00 | 35108 |
| *Mammalia\** | 50 | 290 | 129.24 | 48.69 | 4785 |
| *Actinopterygii* | 50 | 290 | 147.76691 | 41.01 | 4835 |
| *Insecta* | 50 | 290 | 161.15654 | 49.99 | 2945 |
| *Archaea* | 53 | 288 | 154.88 | 47.67 | 851 |
| *Arthropoda\** | 50 | 285 | 139.90 | 43.87 | 1094 |
| *Spermatophyta* | 50 | 290 | 151.97439 | 45.78 | 15188 |
| *Aves* | 51 | 288 | 149.28199 | 10.42 | 9252 |
| *DAASIA* | 55 | 248 | 167.32 | 46.30 | 6988 |
| *Viridiplantae\** | 60 | 276 | 156.03 | 48.86 | 117 |
| *sauria* | 61 | 288 | 157.40476 | 62.39 | 42 |
| *amphibia* | 95 | 181 | 140.75 | 38.35 | 4 |
| *Cyclostomata* | 83 | 272 | 181.50 | 80.47 | 7 |
| *Chondrichthyes* | 78 | 221 | 159.50 | 63.61 | 4 |
| *ABPNNR* | 59 | 267 | 187.17 | 88.41 | 752 |
| *PCM* | 68 | 298 | 197.20 | 85.64 | 187 |
| *EHT* | 77 | 219 | 186.34 | 92.41 | 49 |
| *HRC* | 61 | 247 | 163.72 | 80.79 | 15 |
| *Unnamed* | 88 | 296 | 157.26 | 49.87 | 271062 |
| Total |  |  |  |  | 3159020 |

Please note that the data in Viridiplantae\* does not include Spermatophyta, the data in Arthropoda\* excludes Insecta, the data in Sauria\* does not include Aves, the data in Primates\* does not include Homo sapiens, and the data in Mammalia\* does not include Primates\*.

**Table S12.**
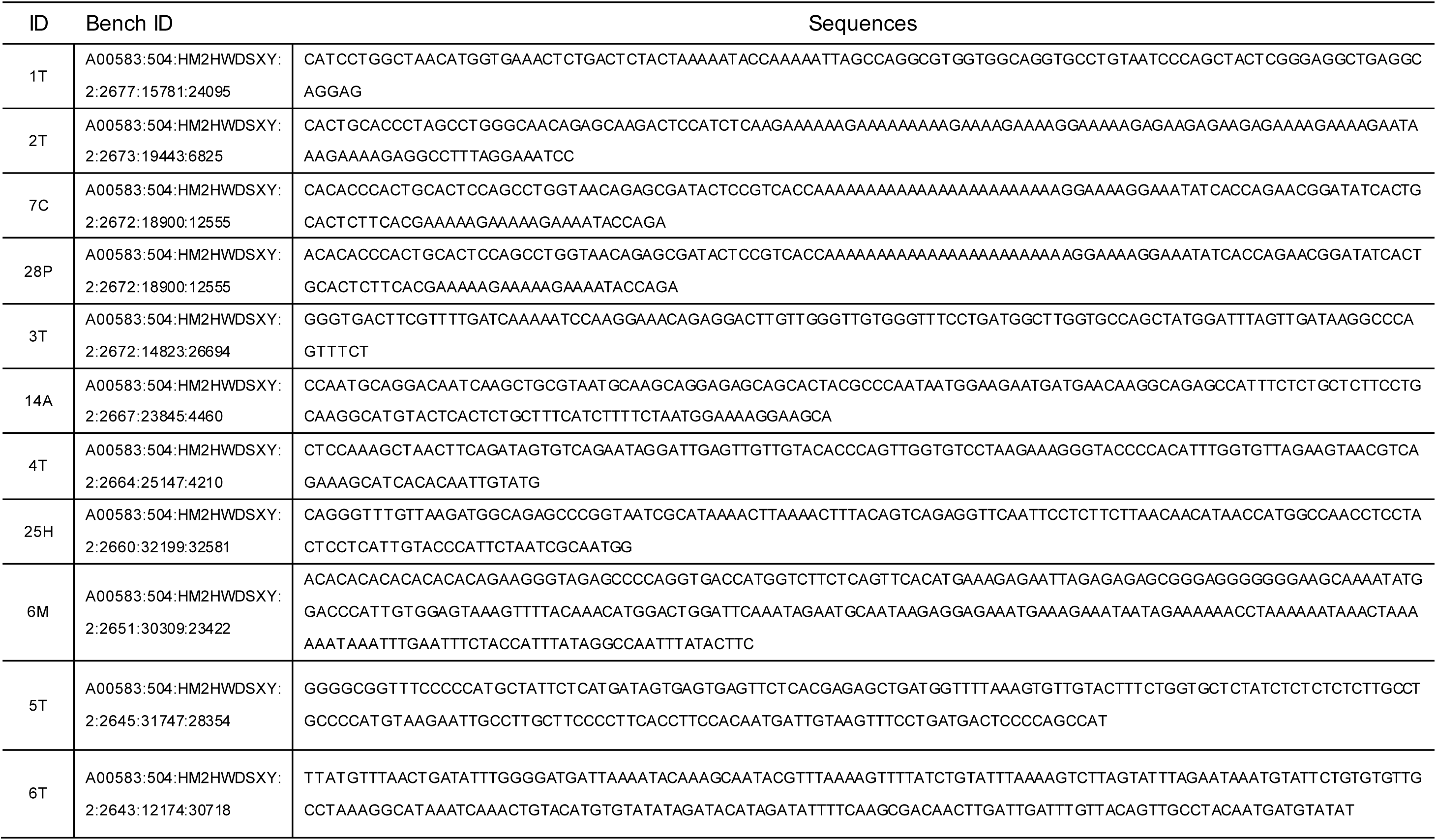
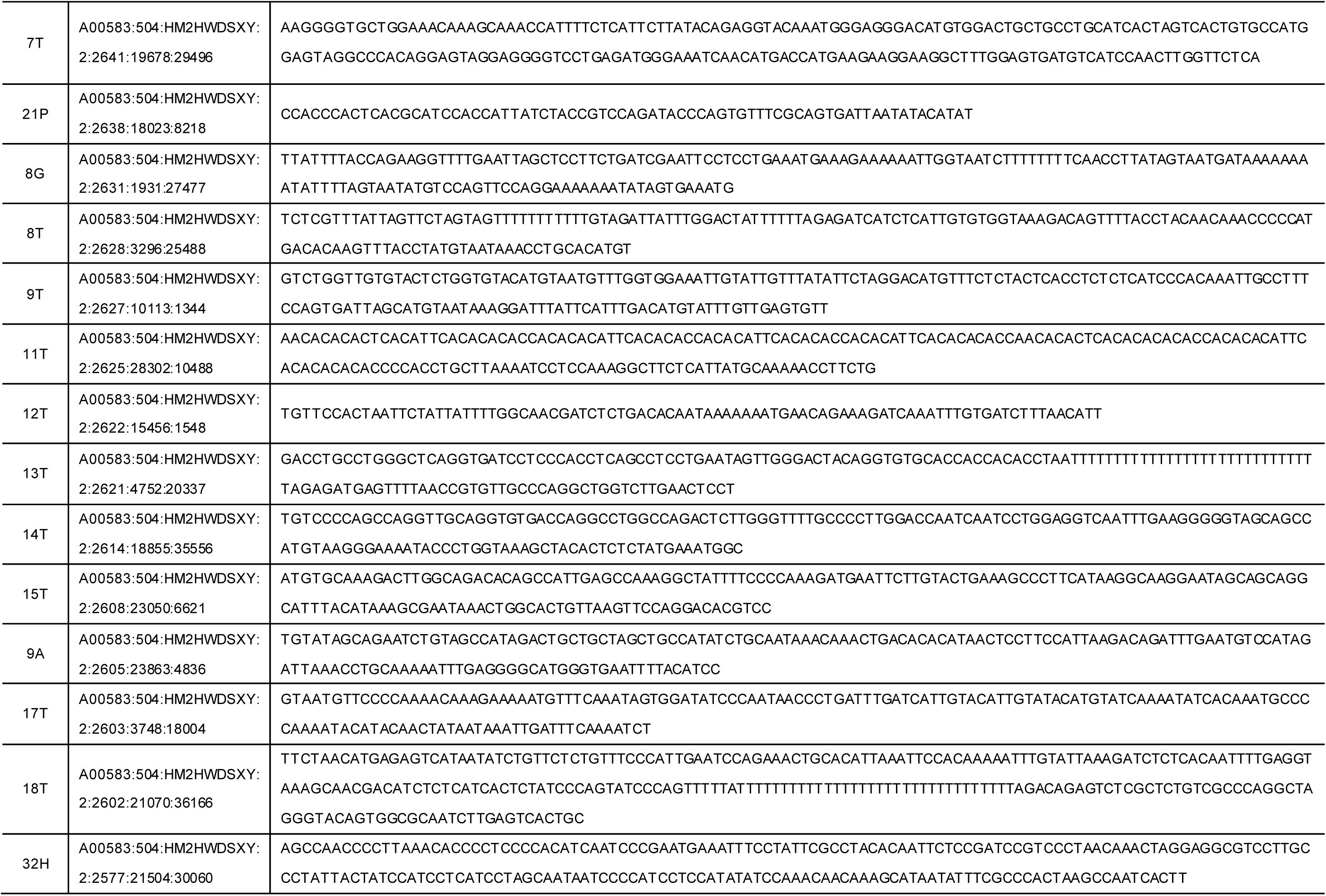
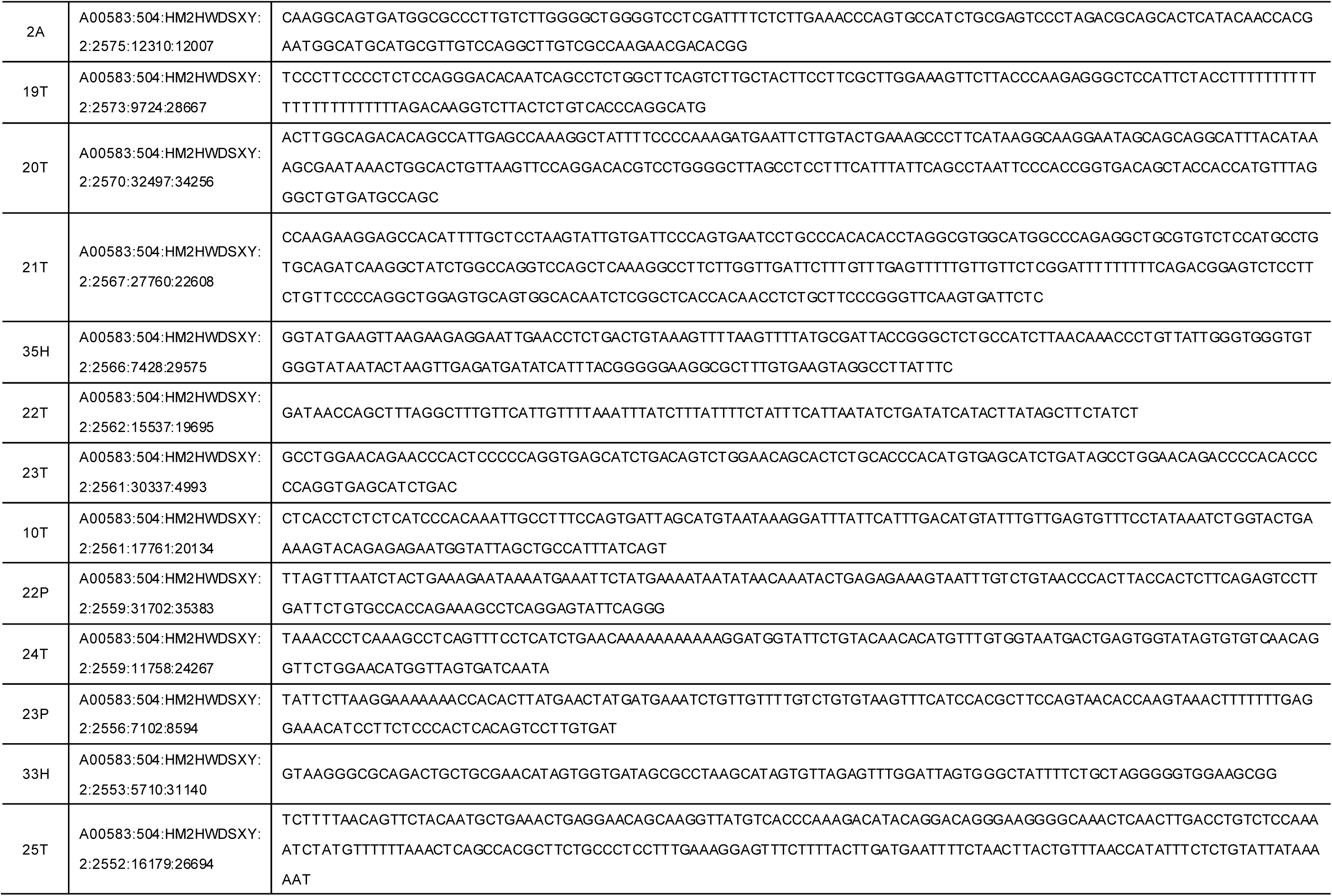
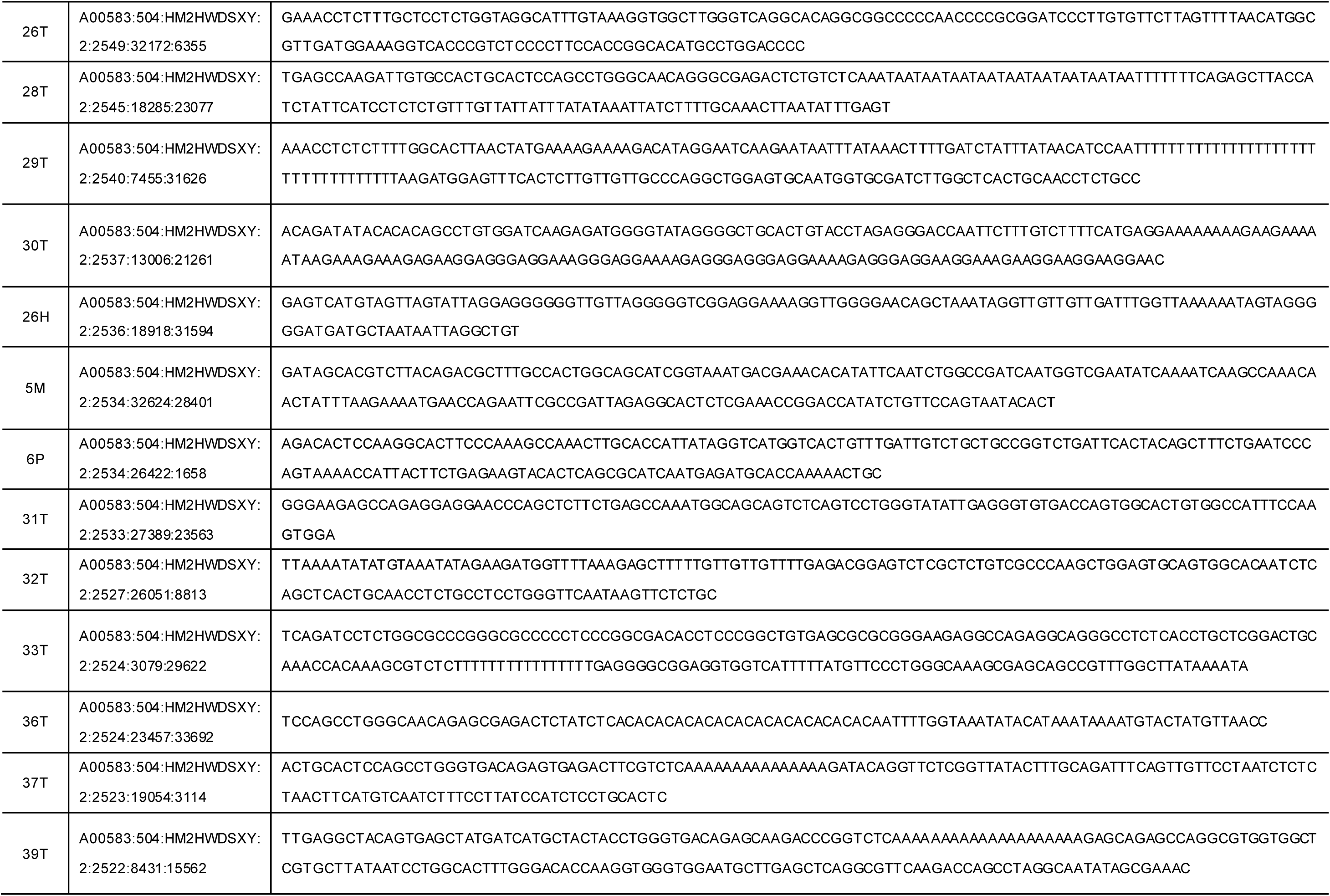
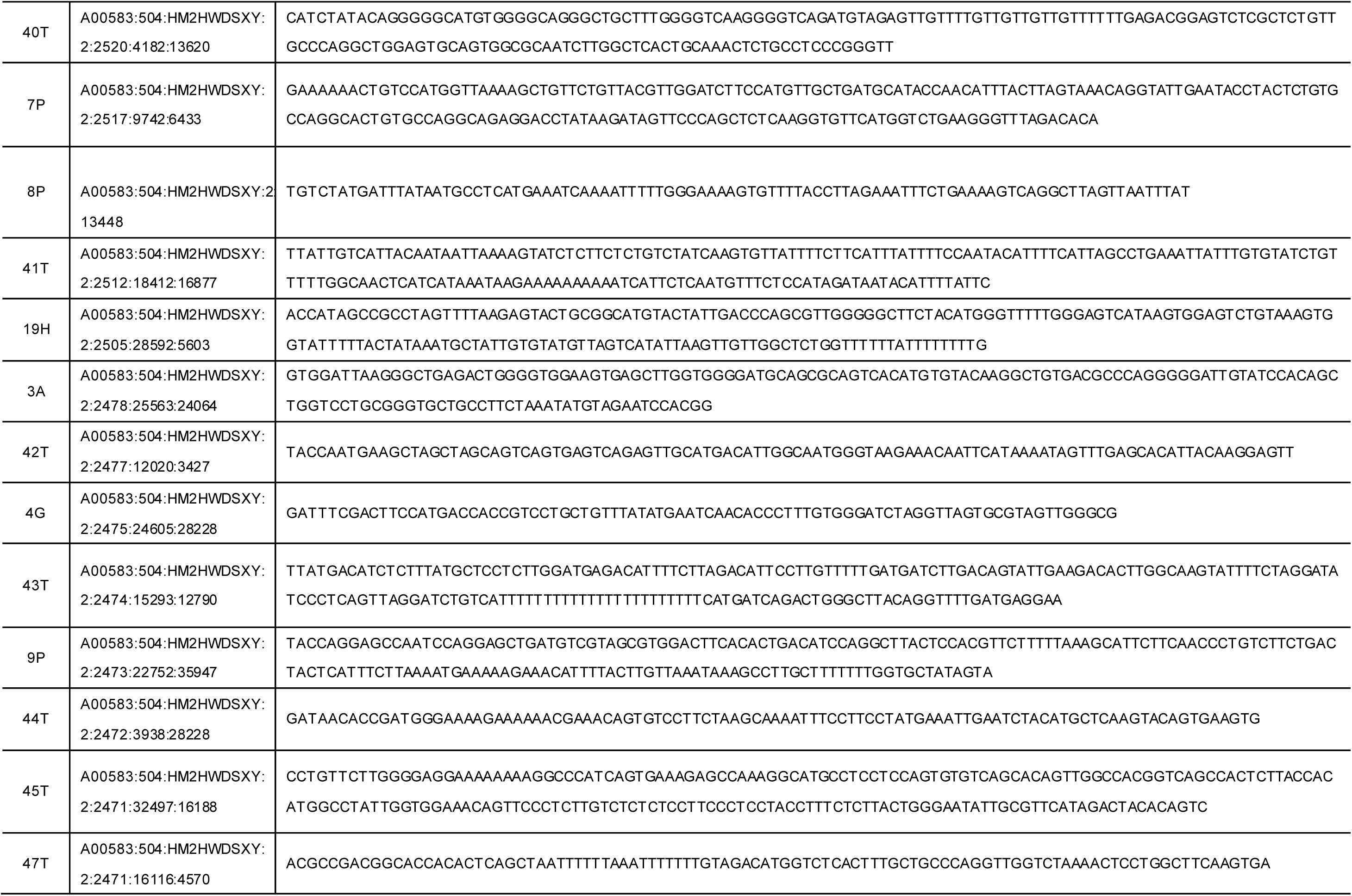
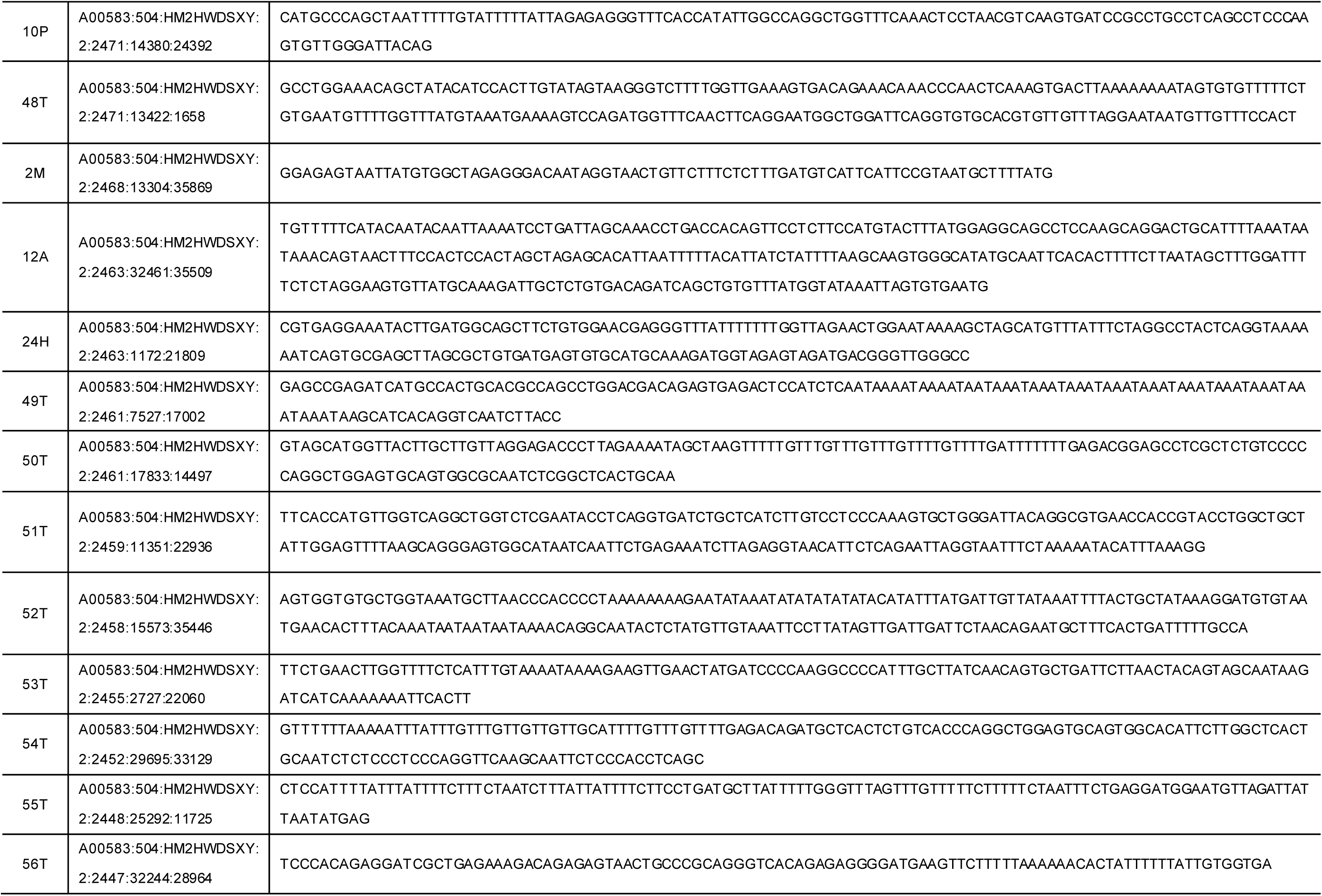
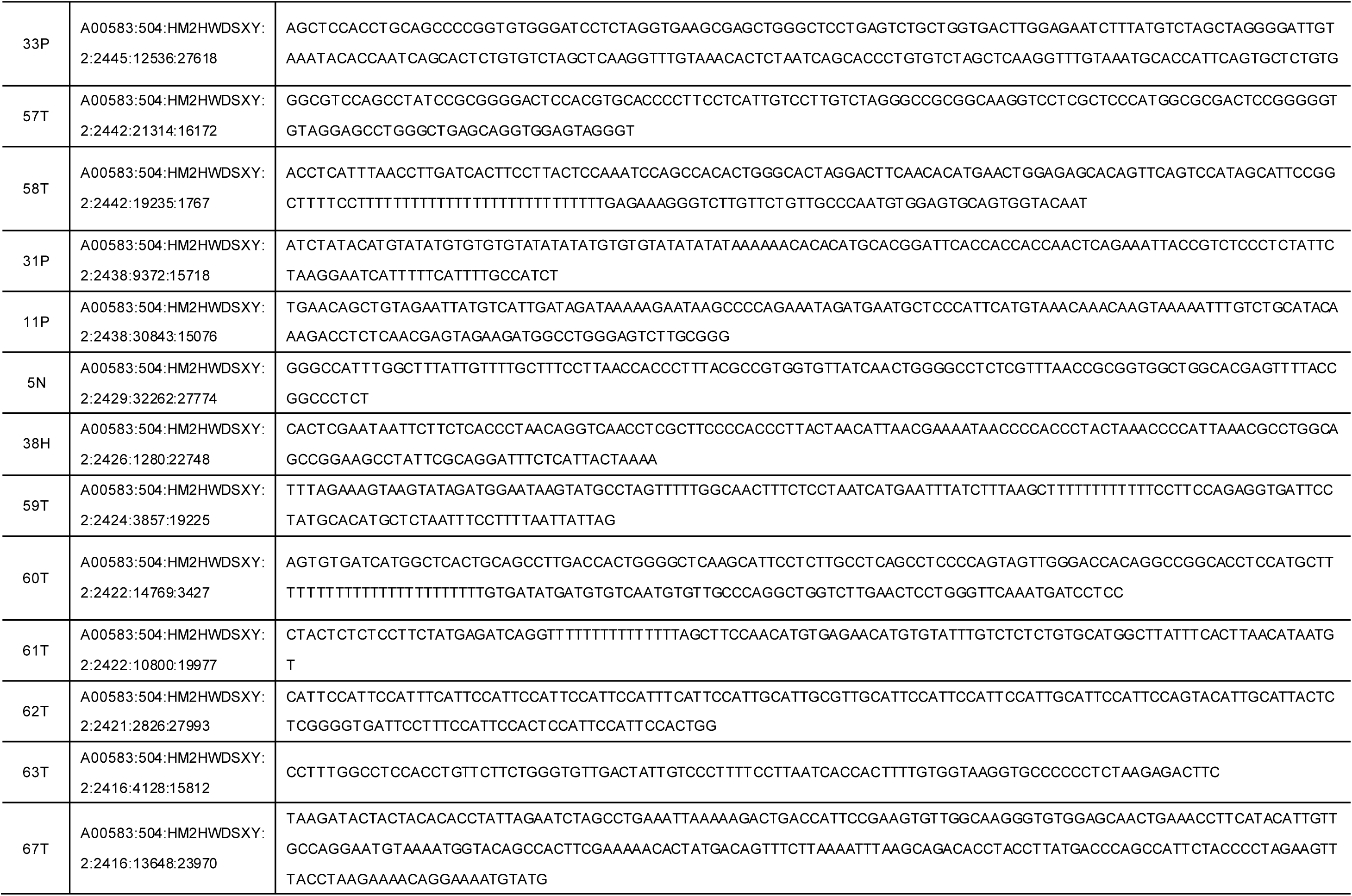
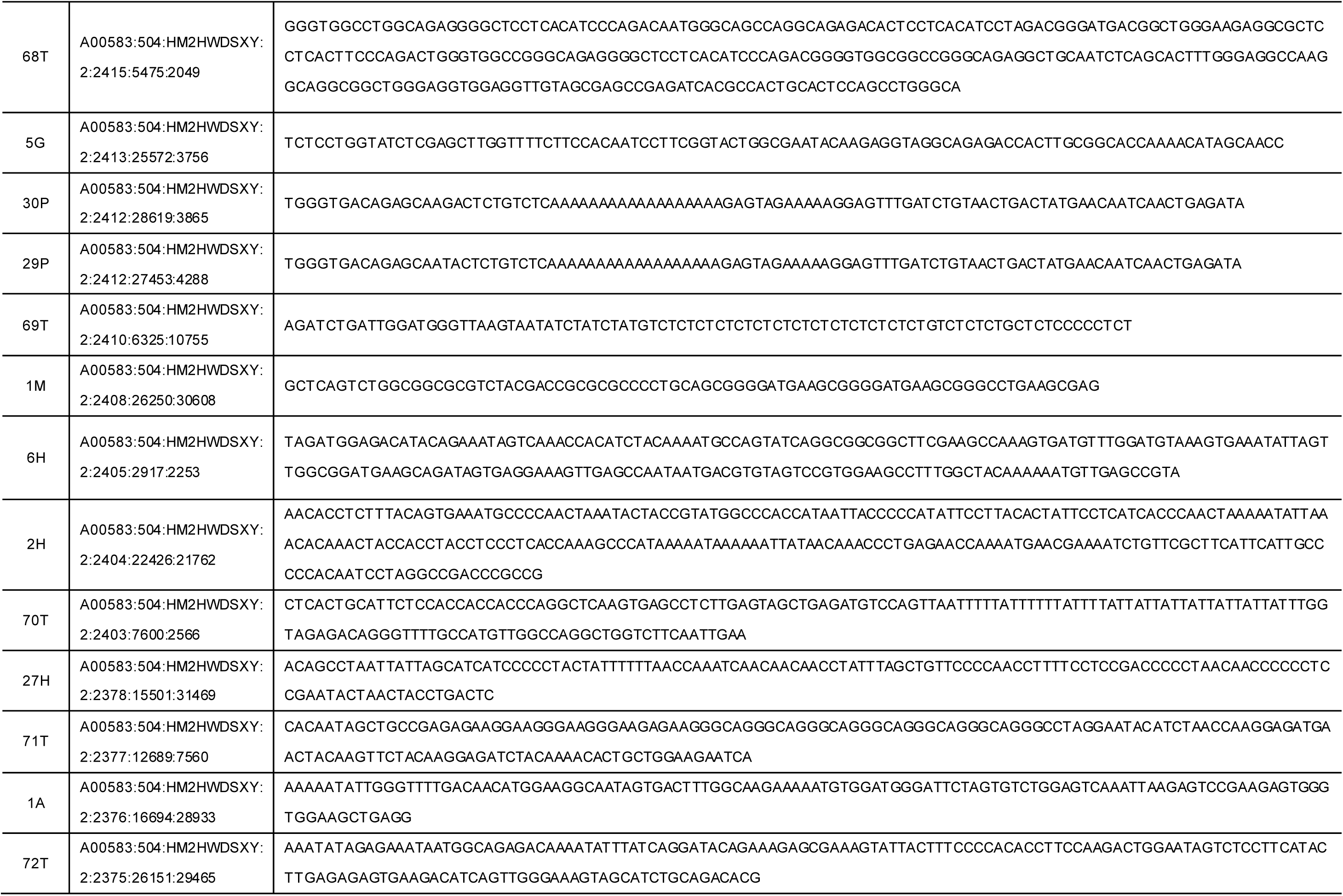
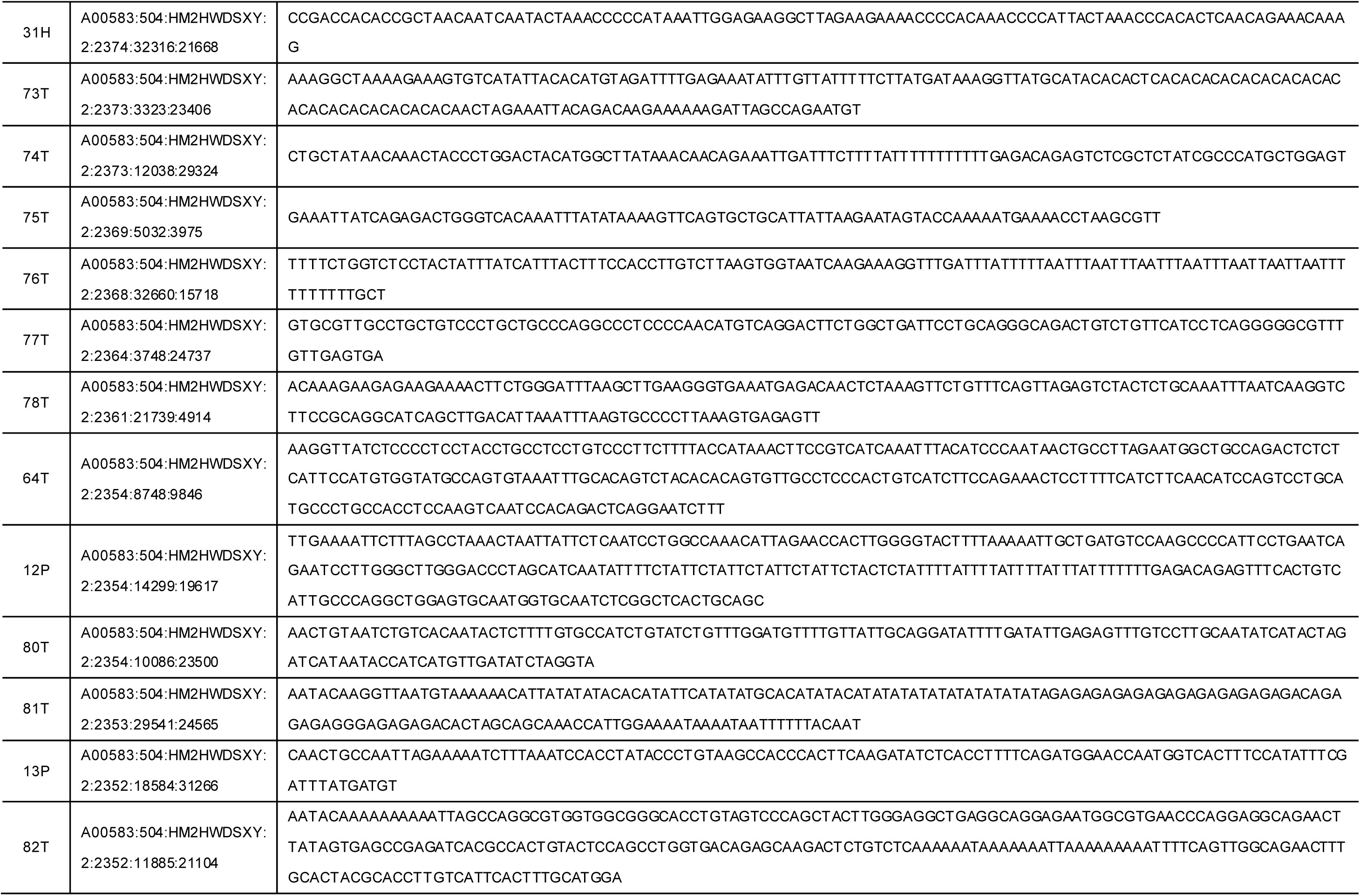
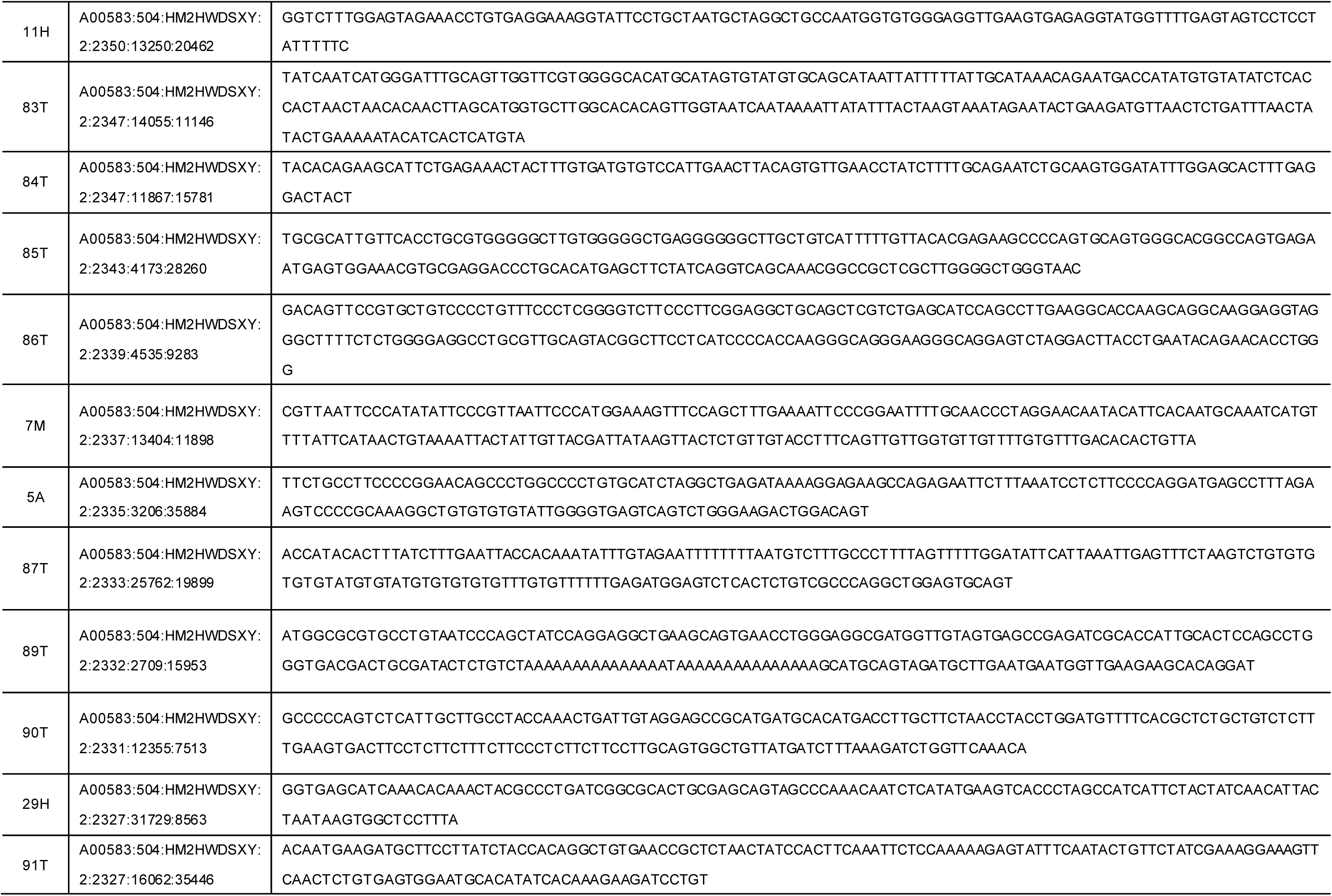
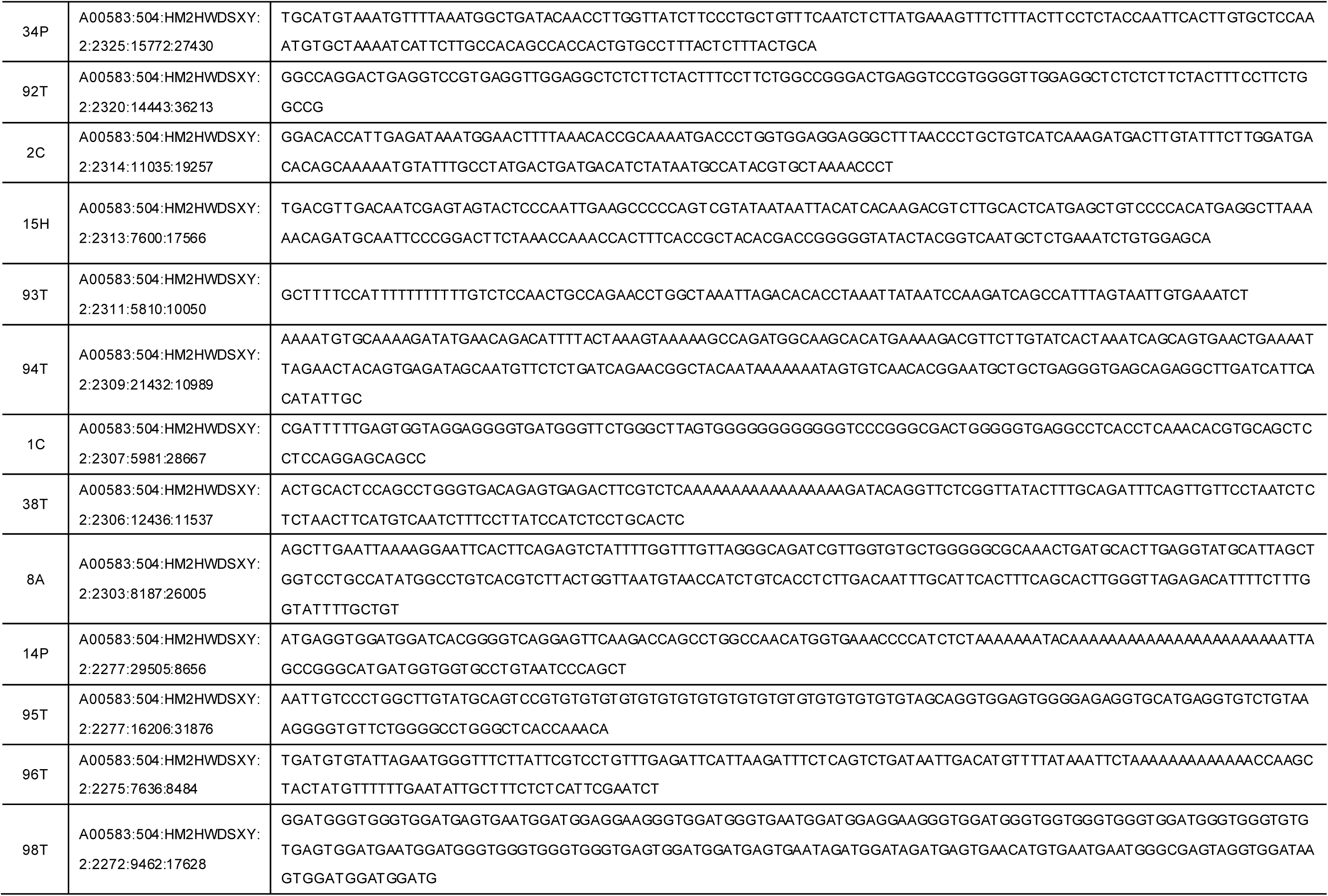
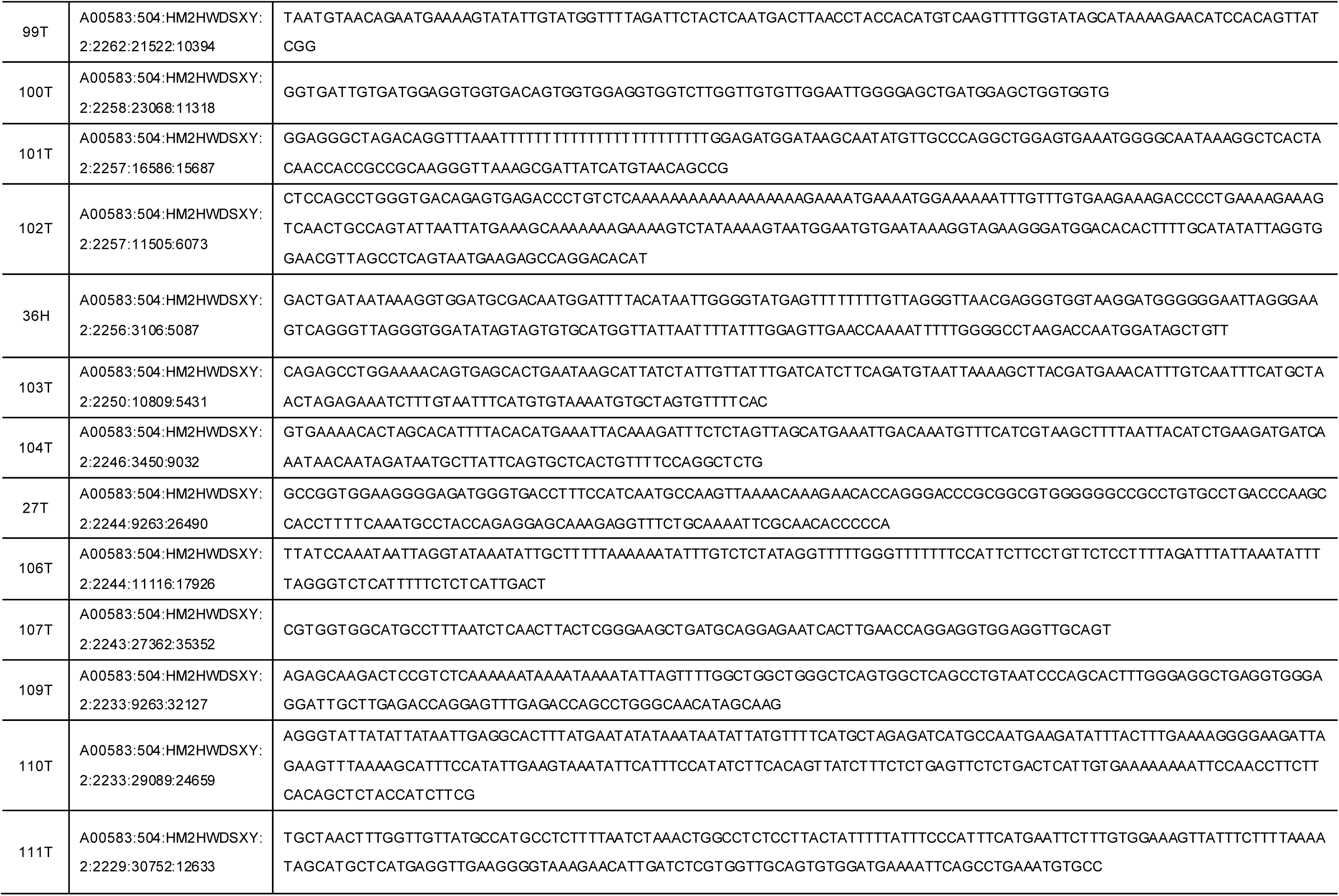
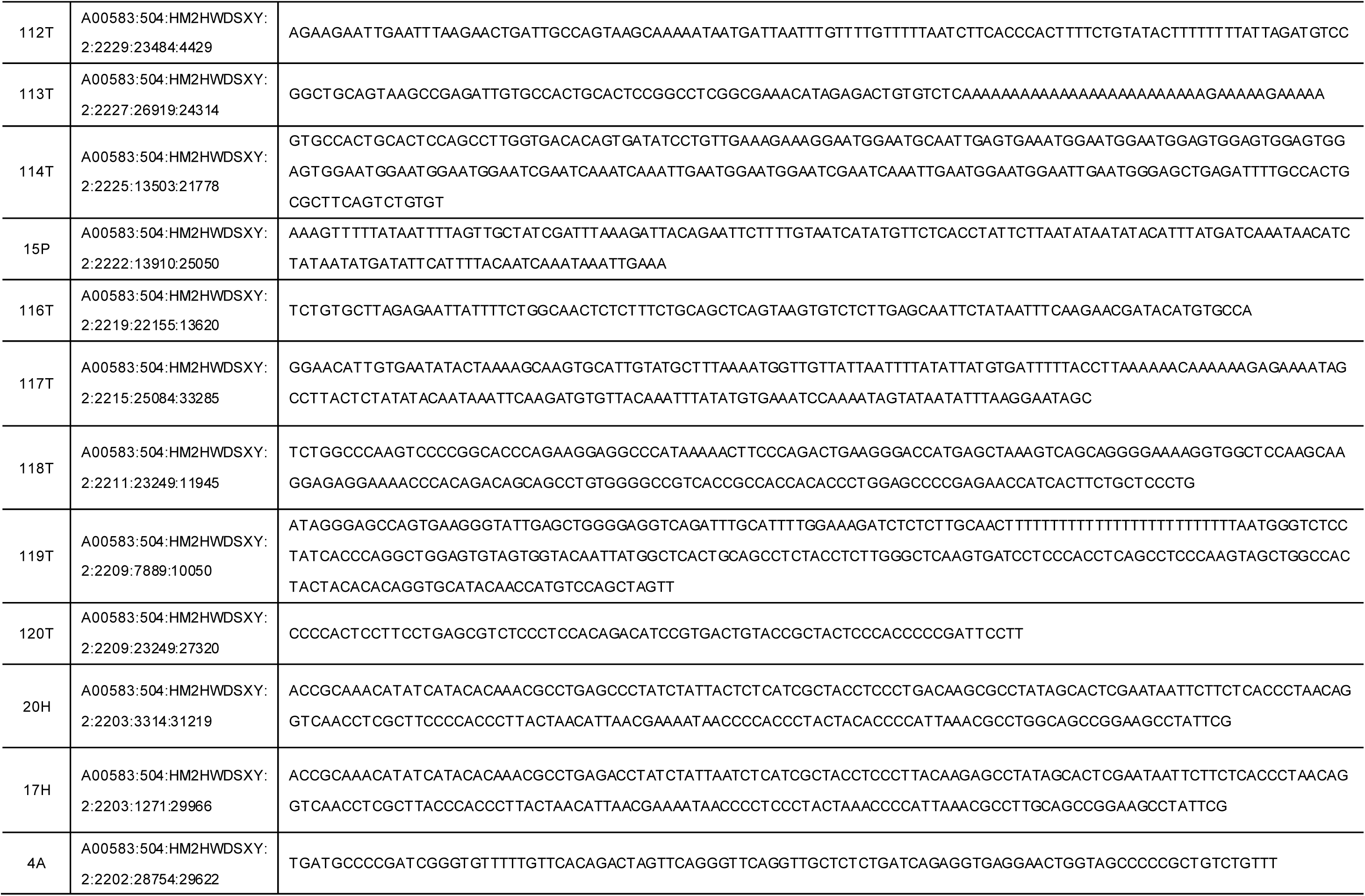
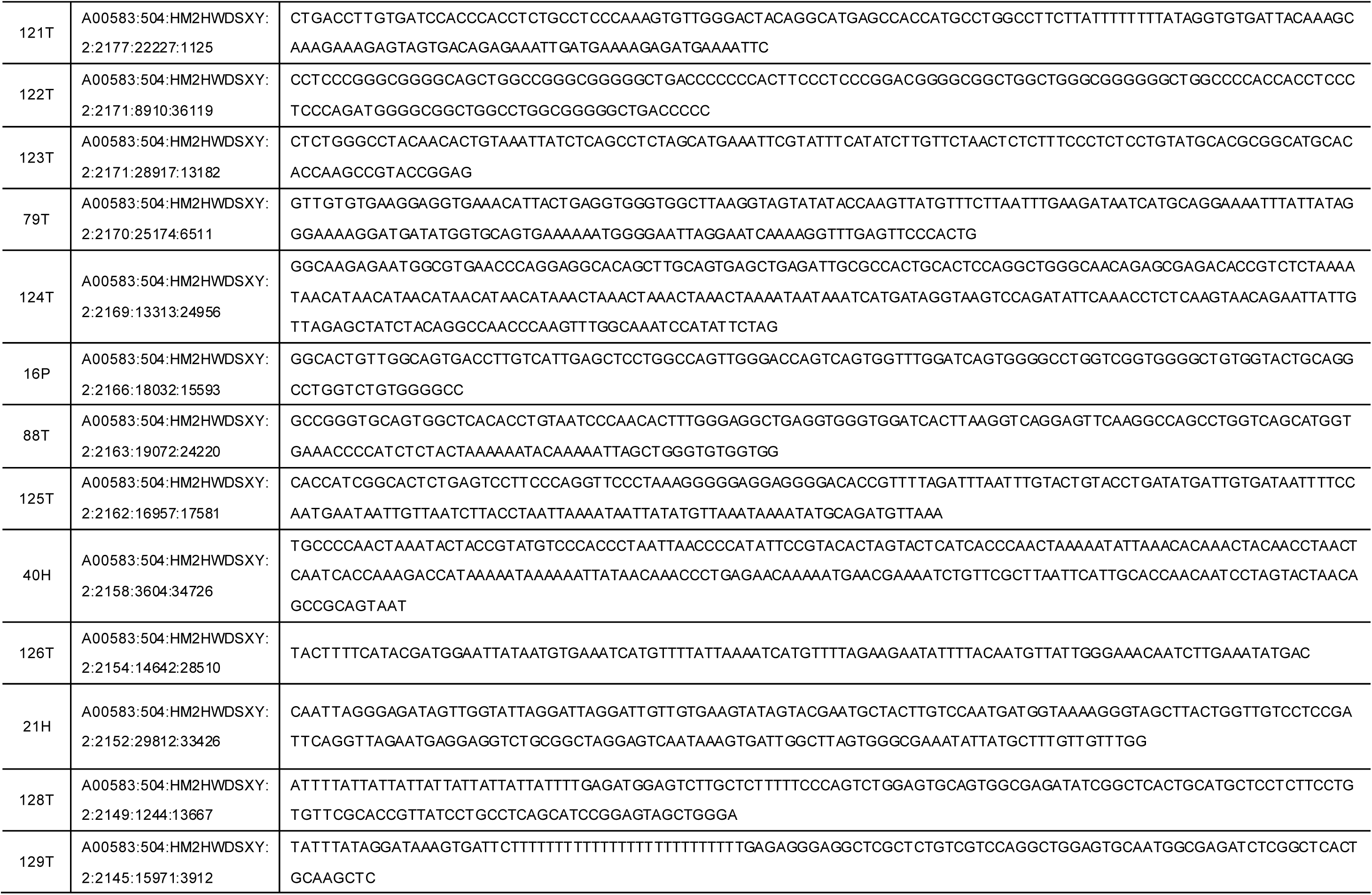
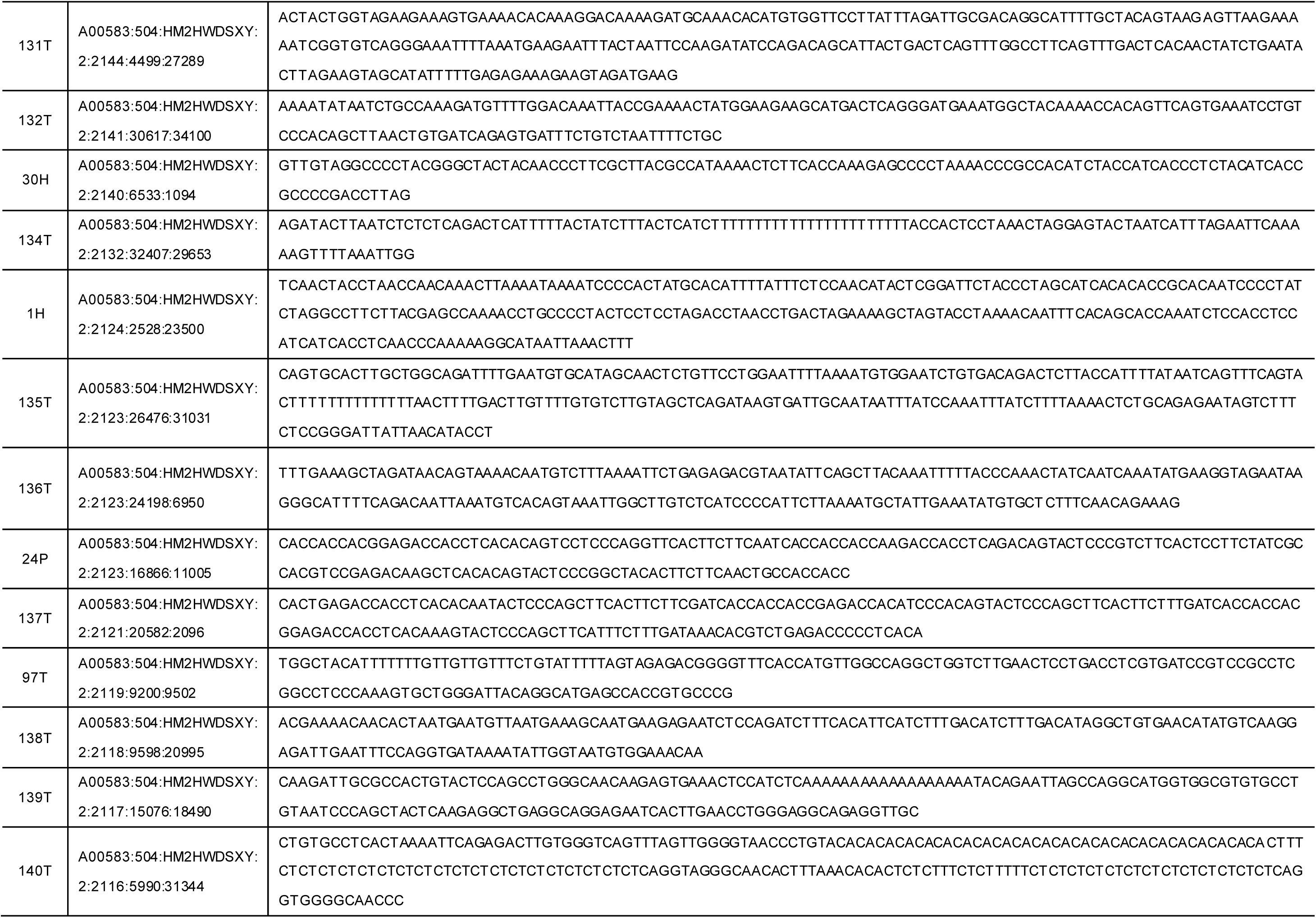
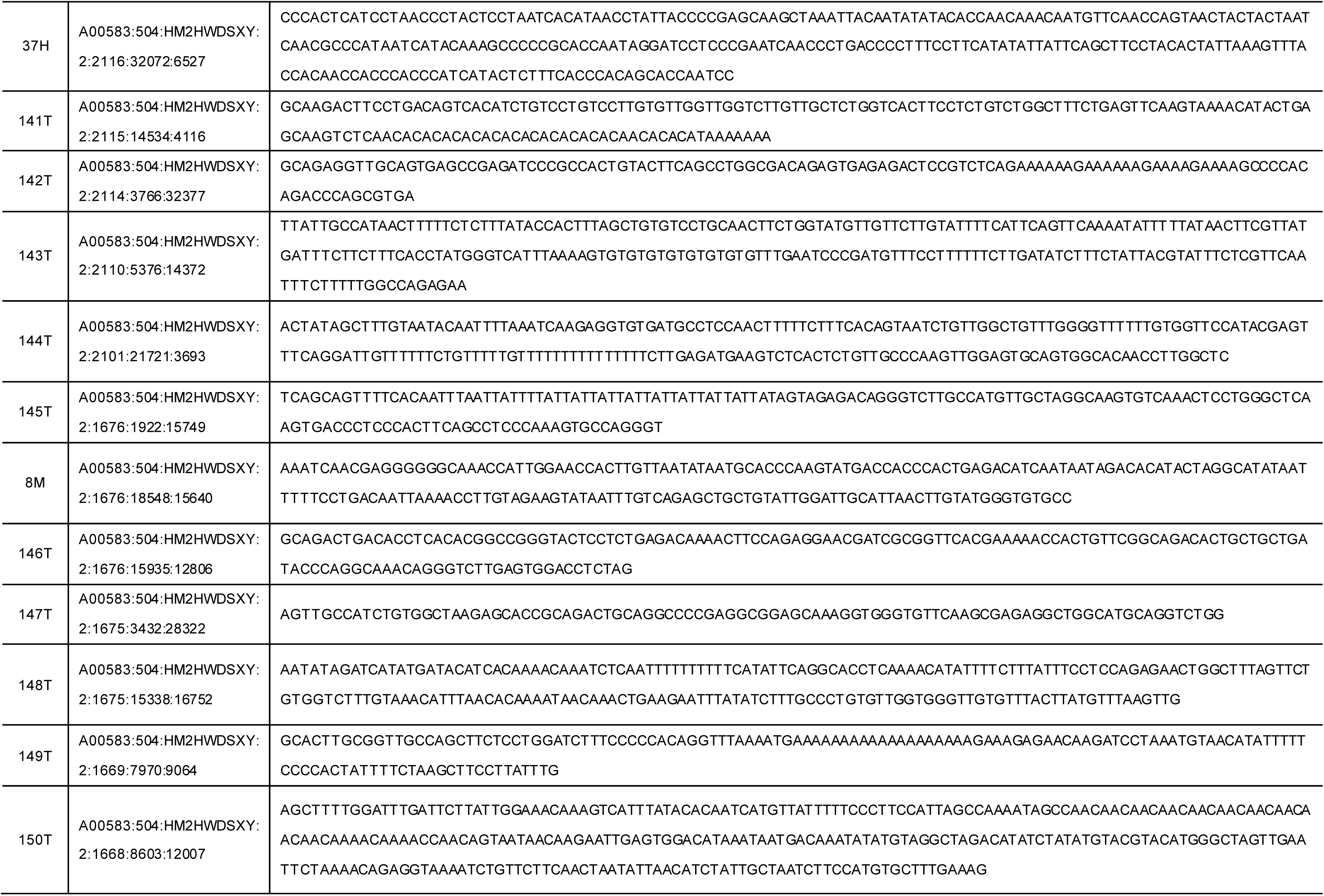
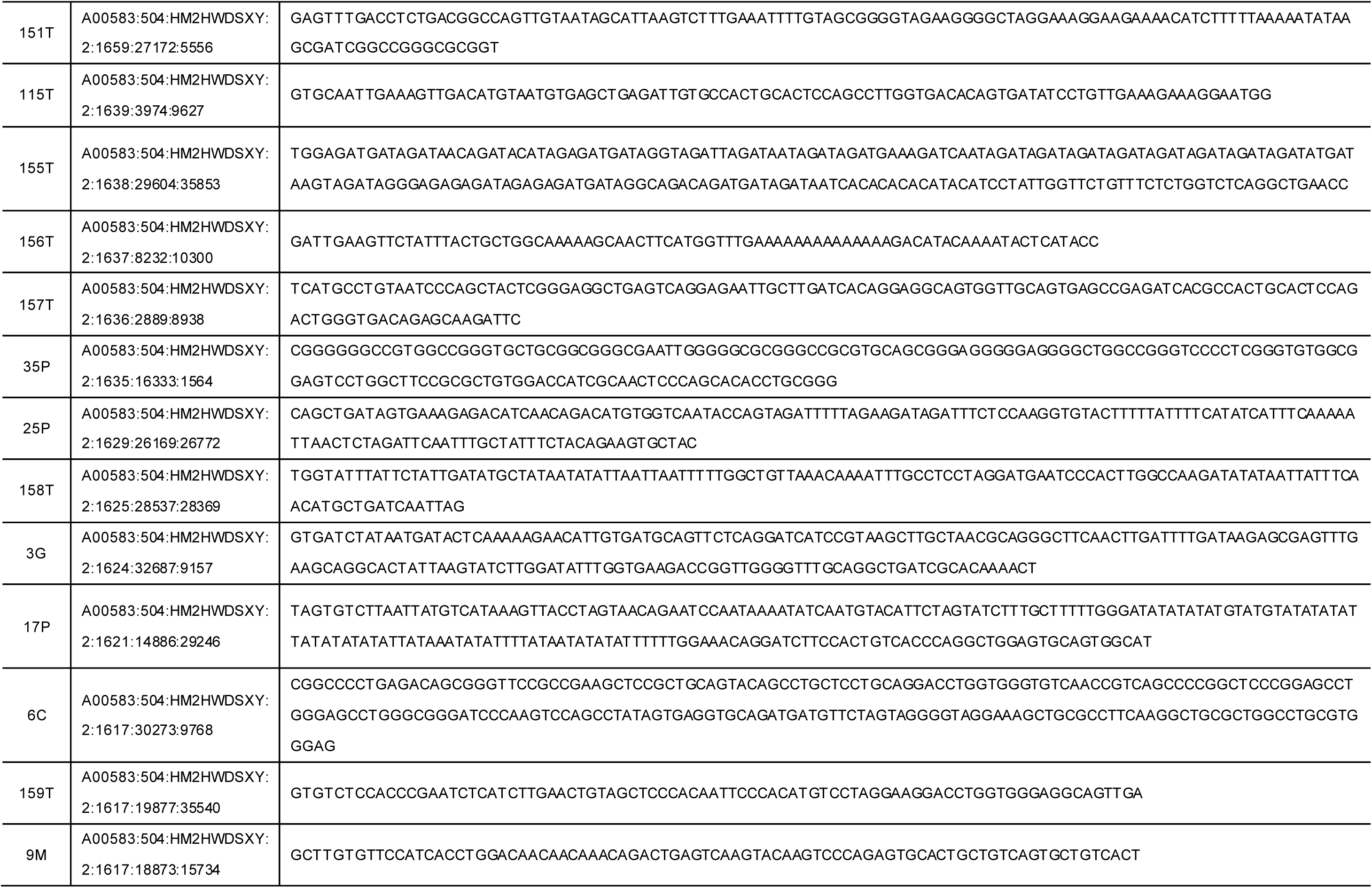
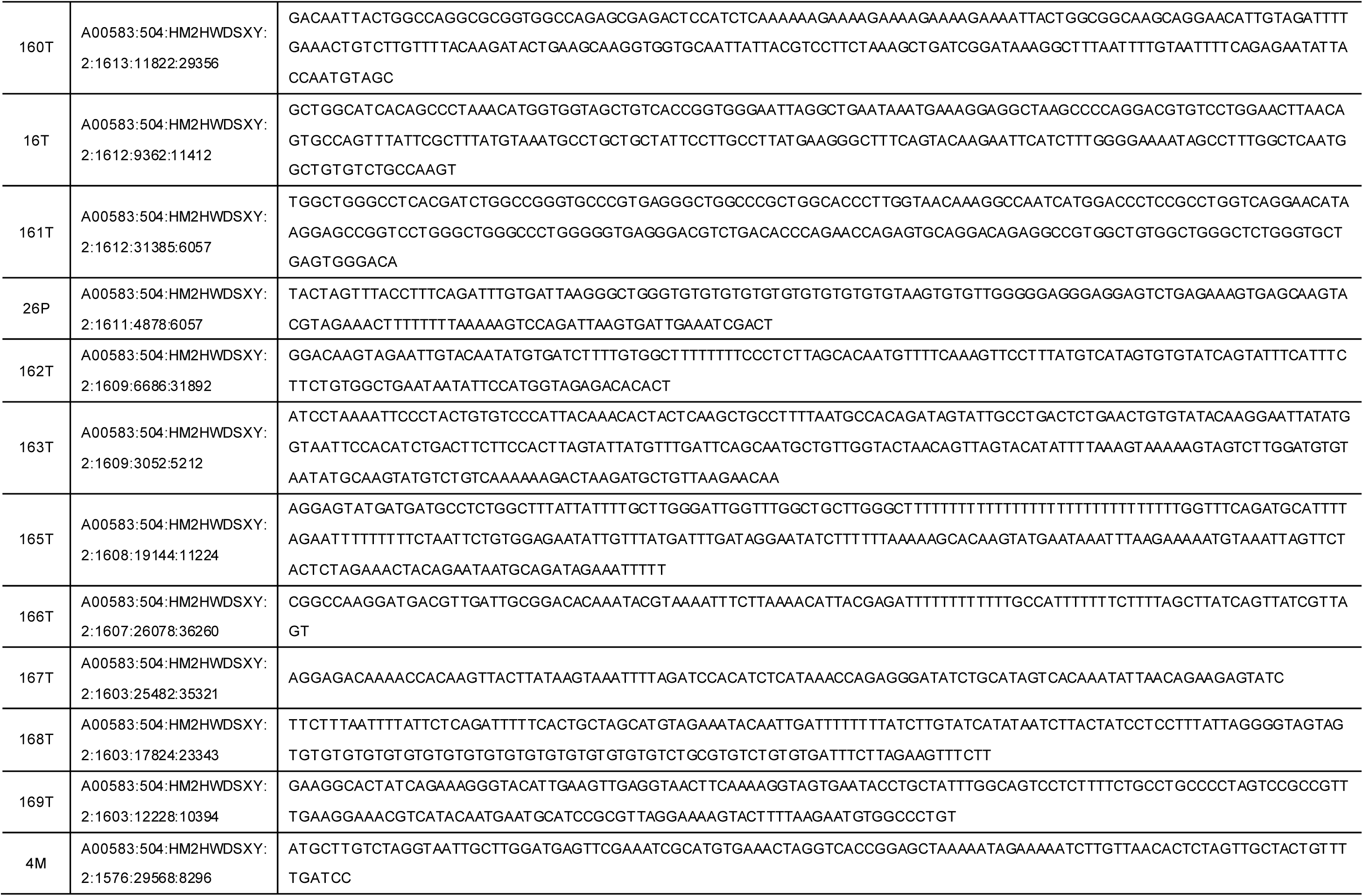
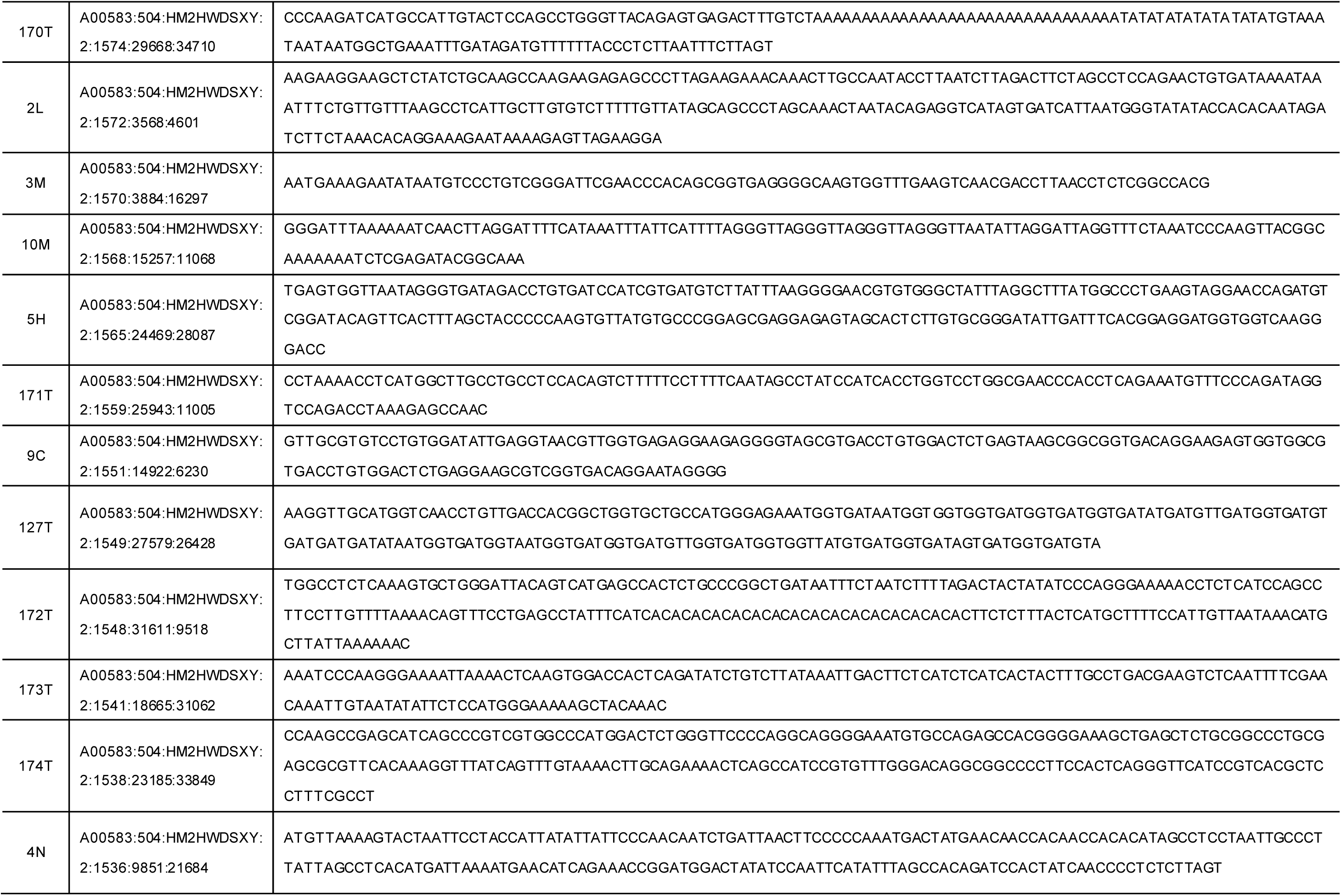
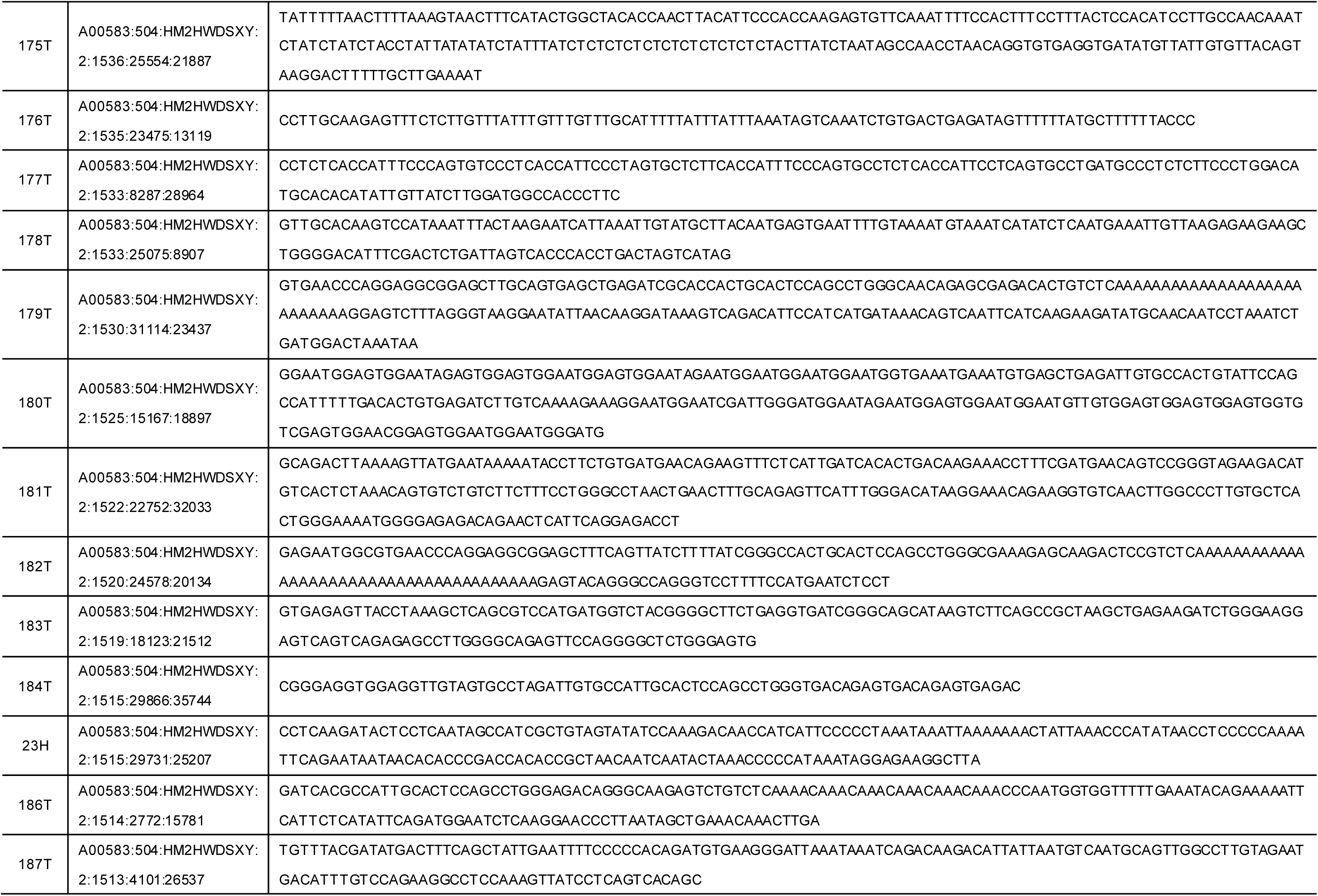
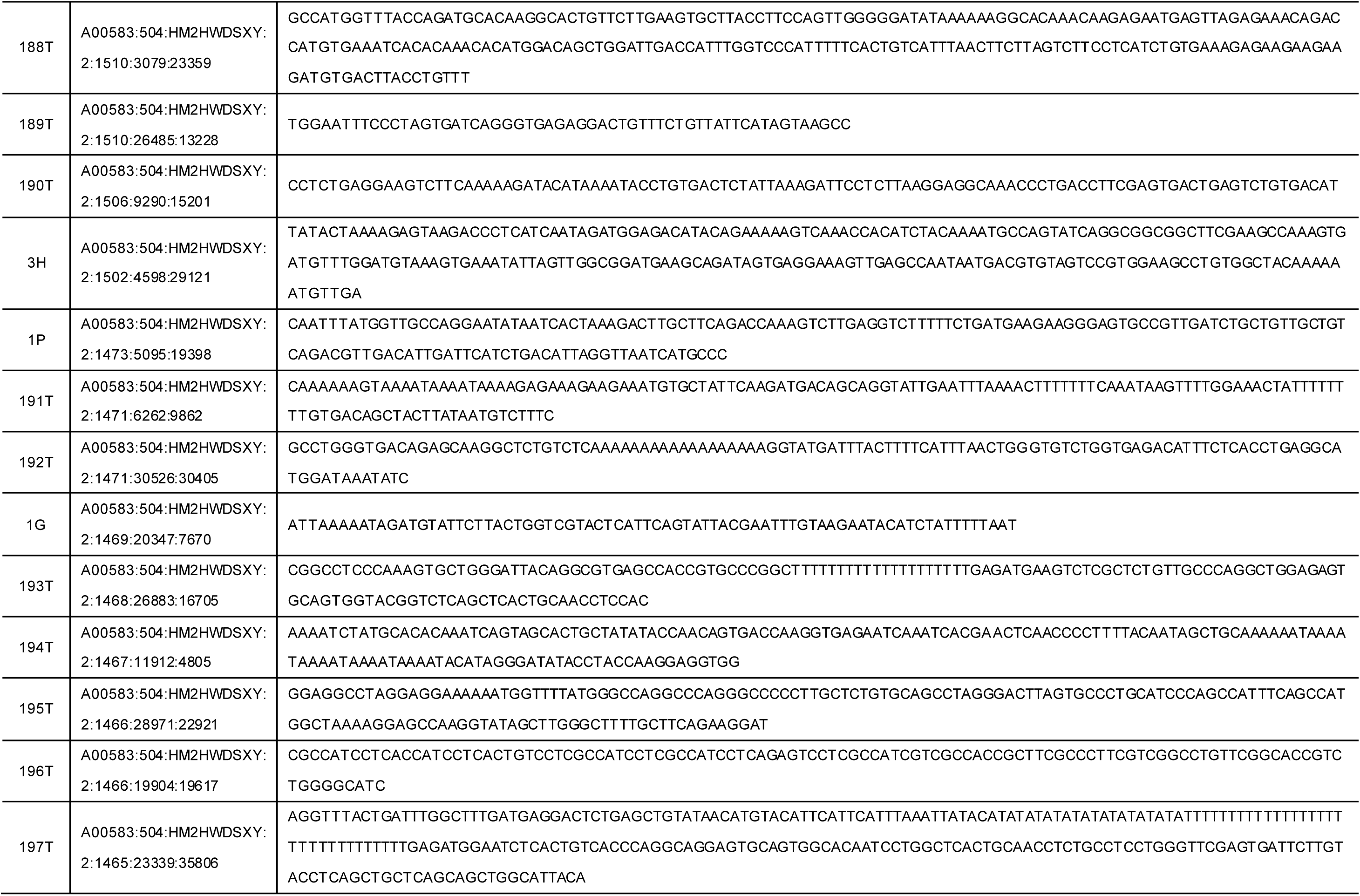
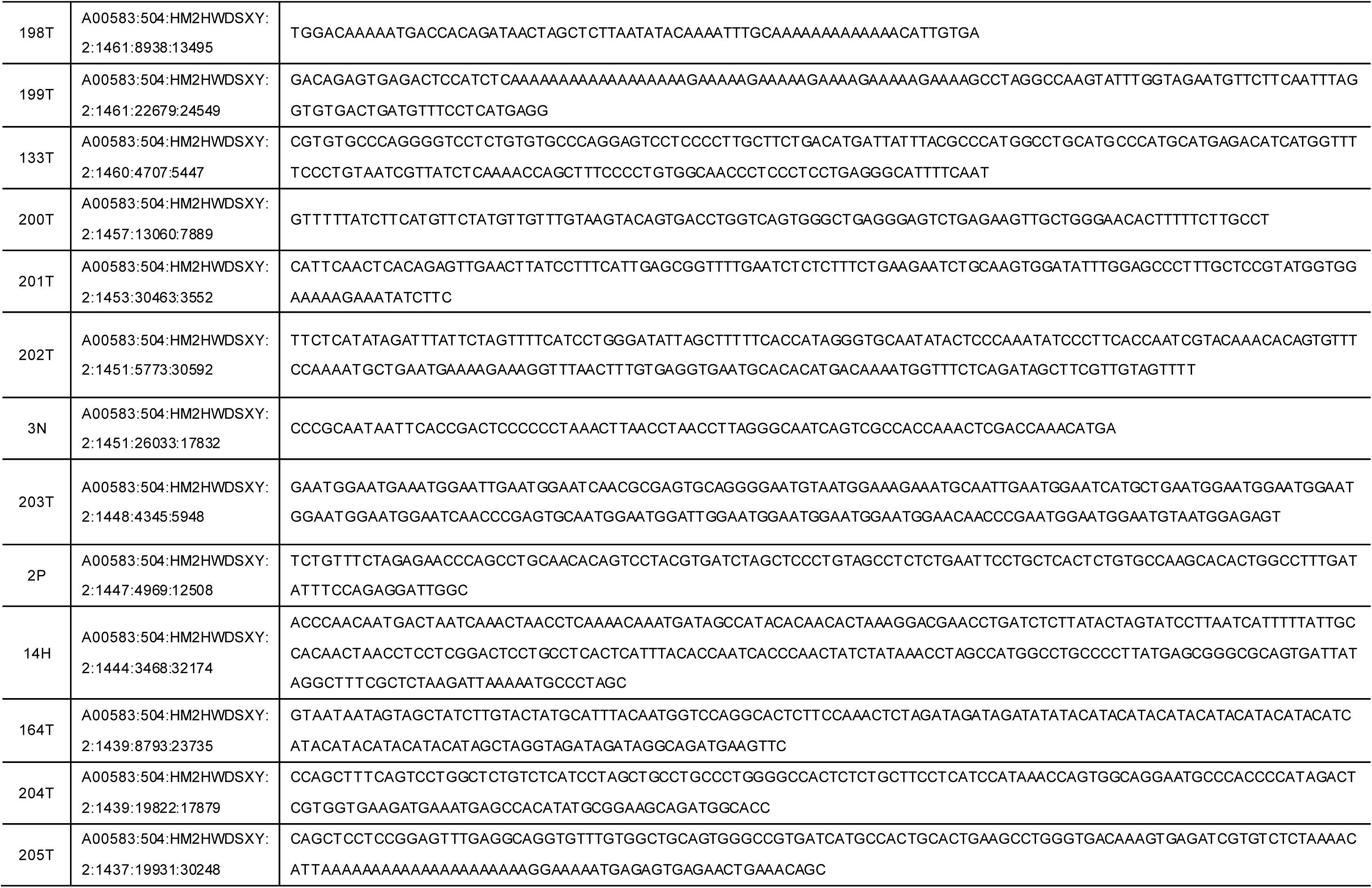
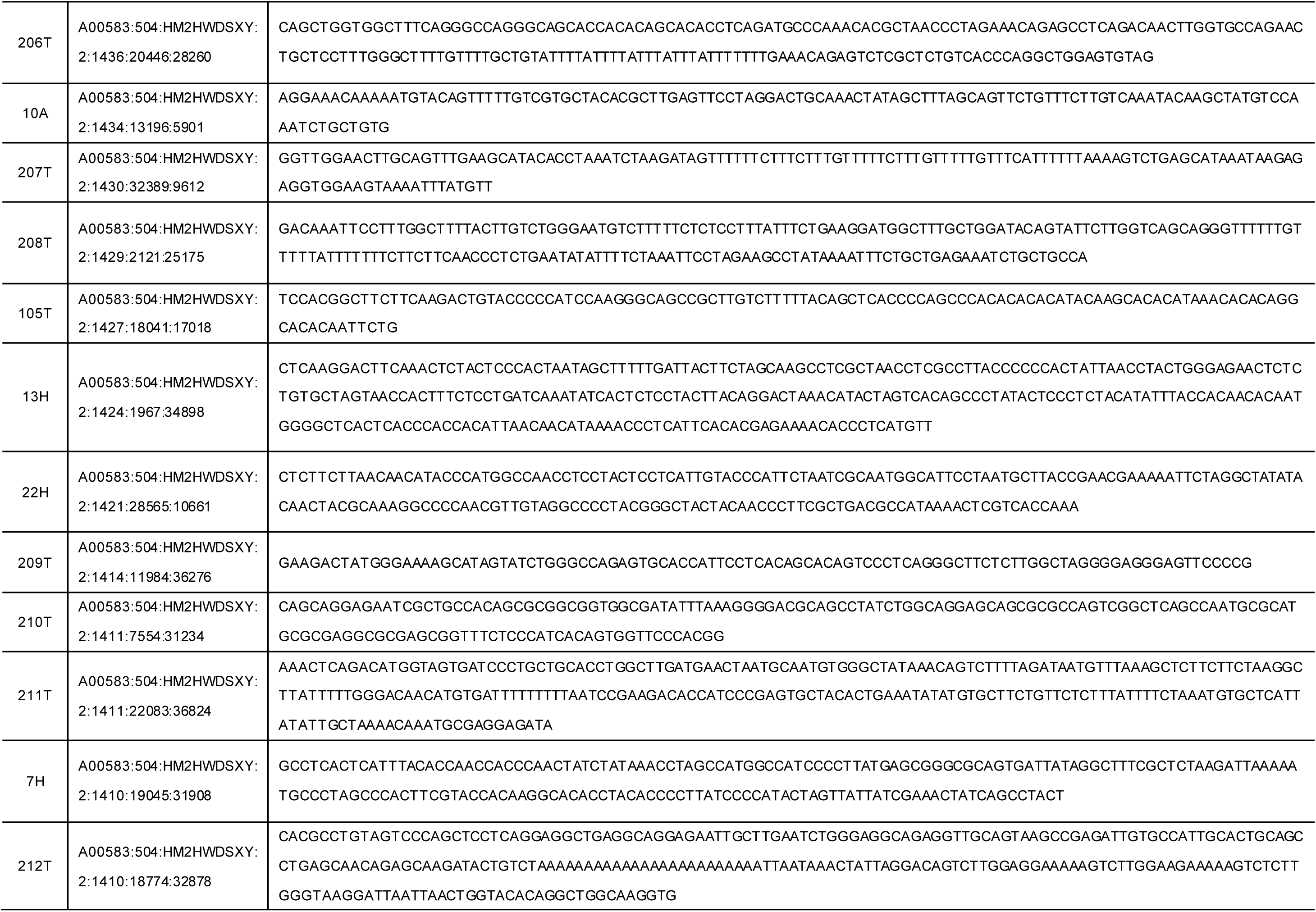
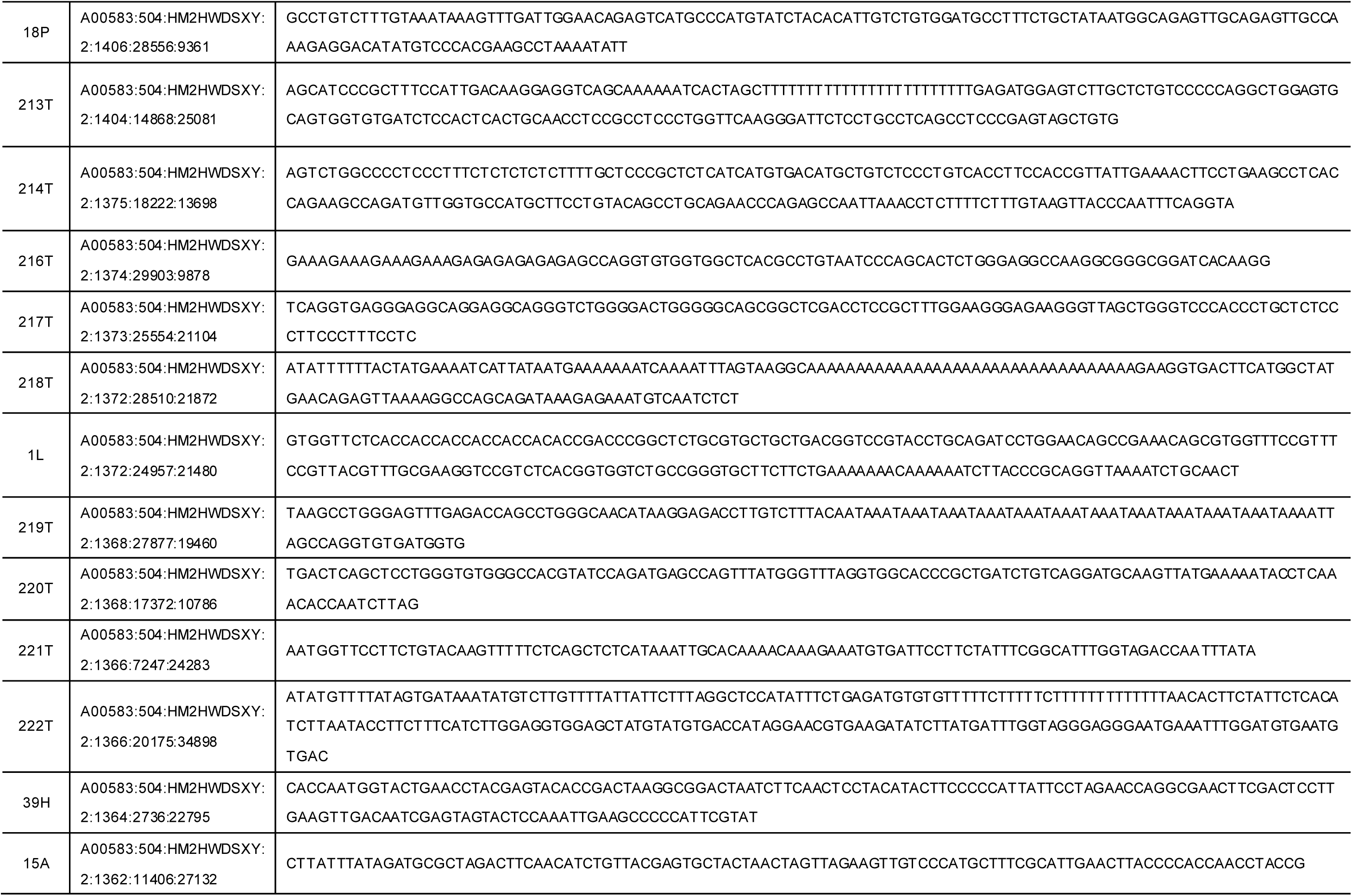
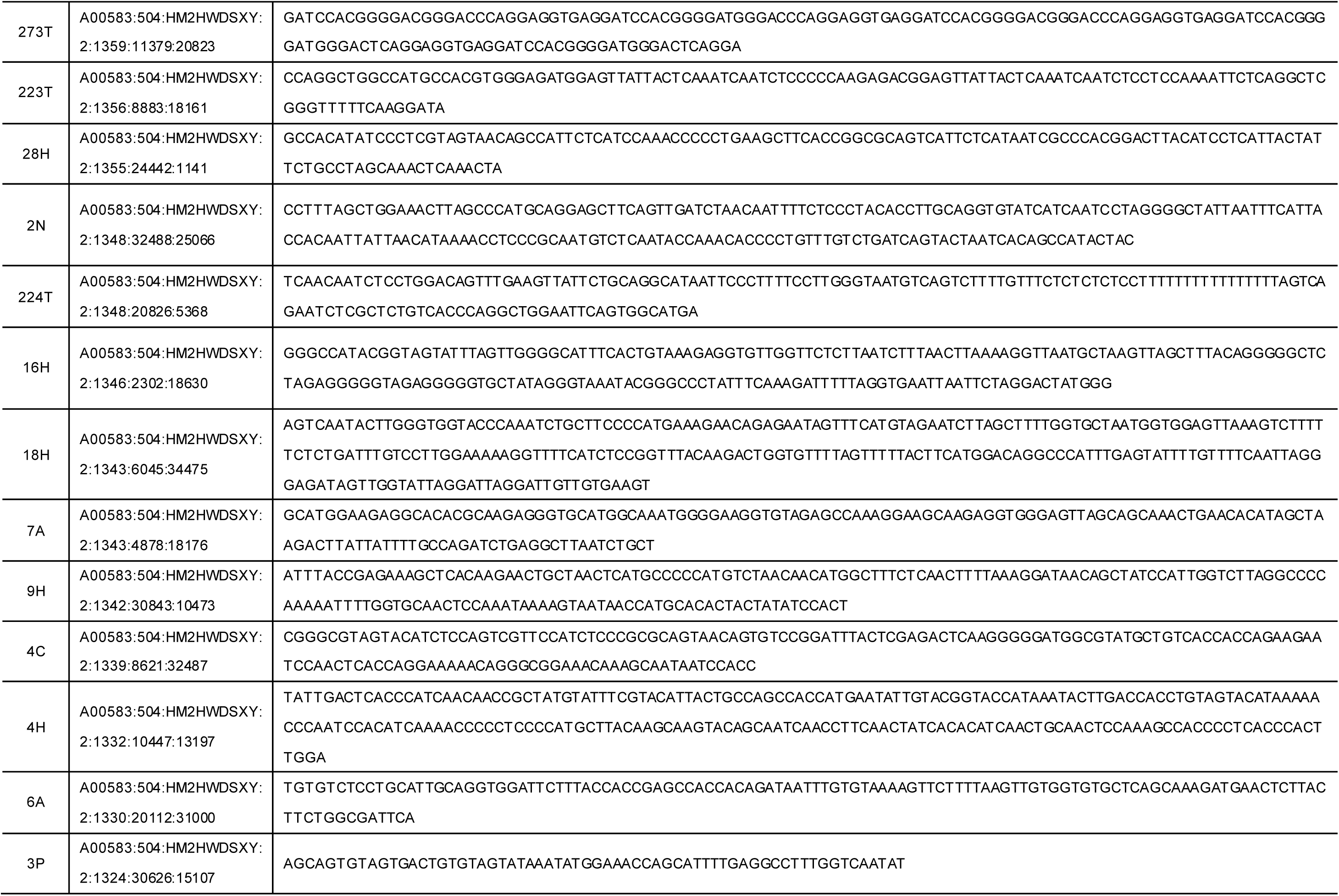
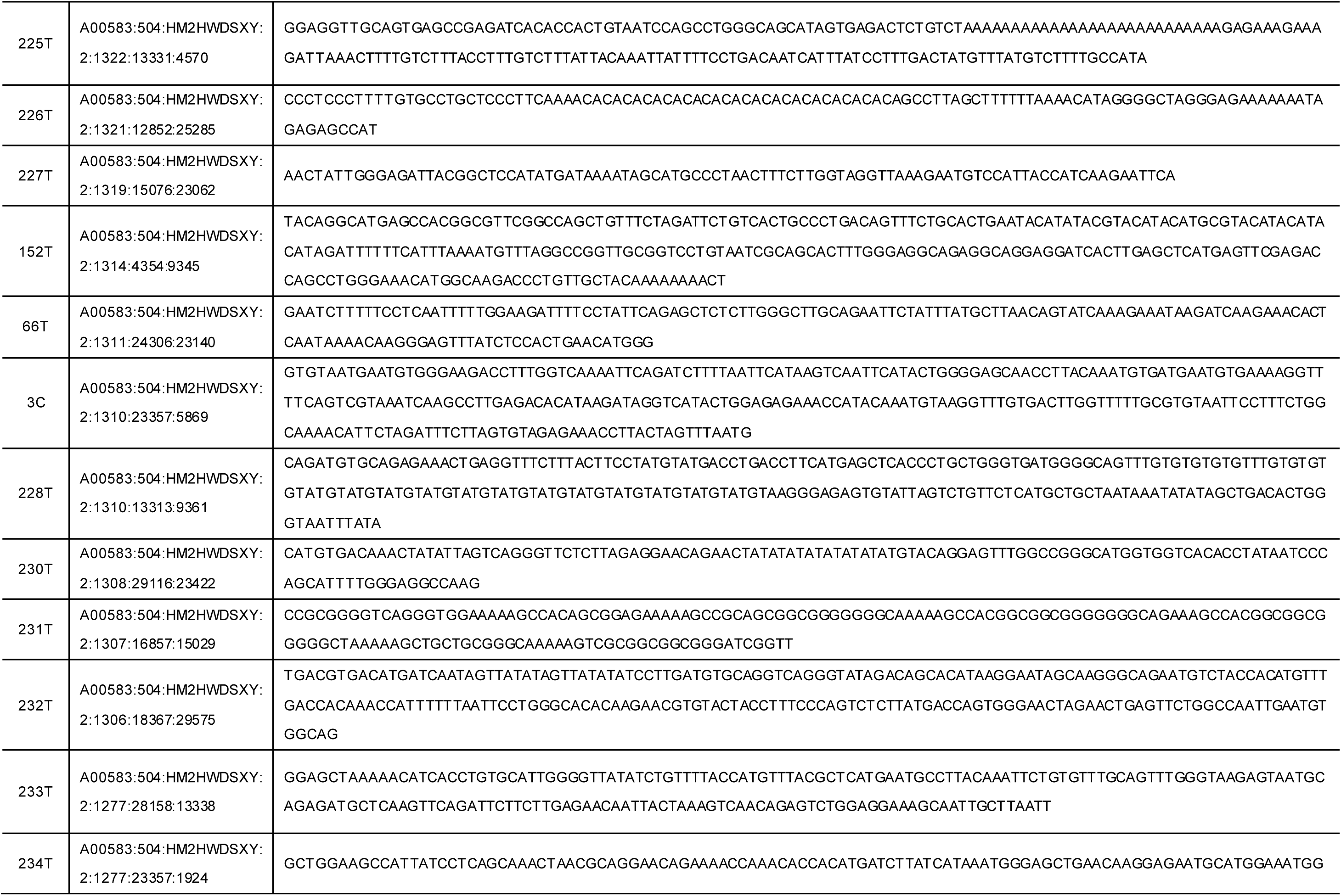
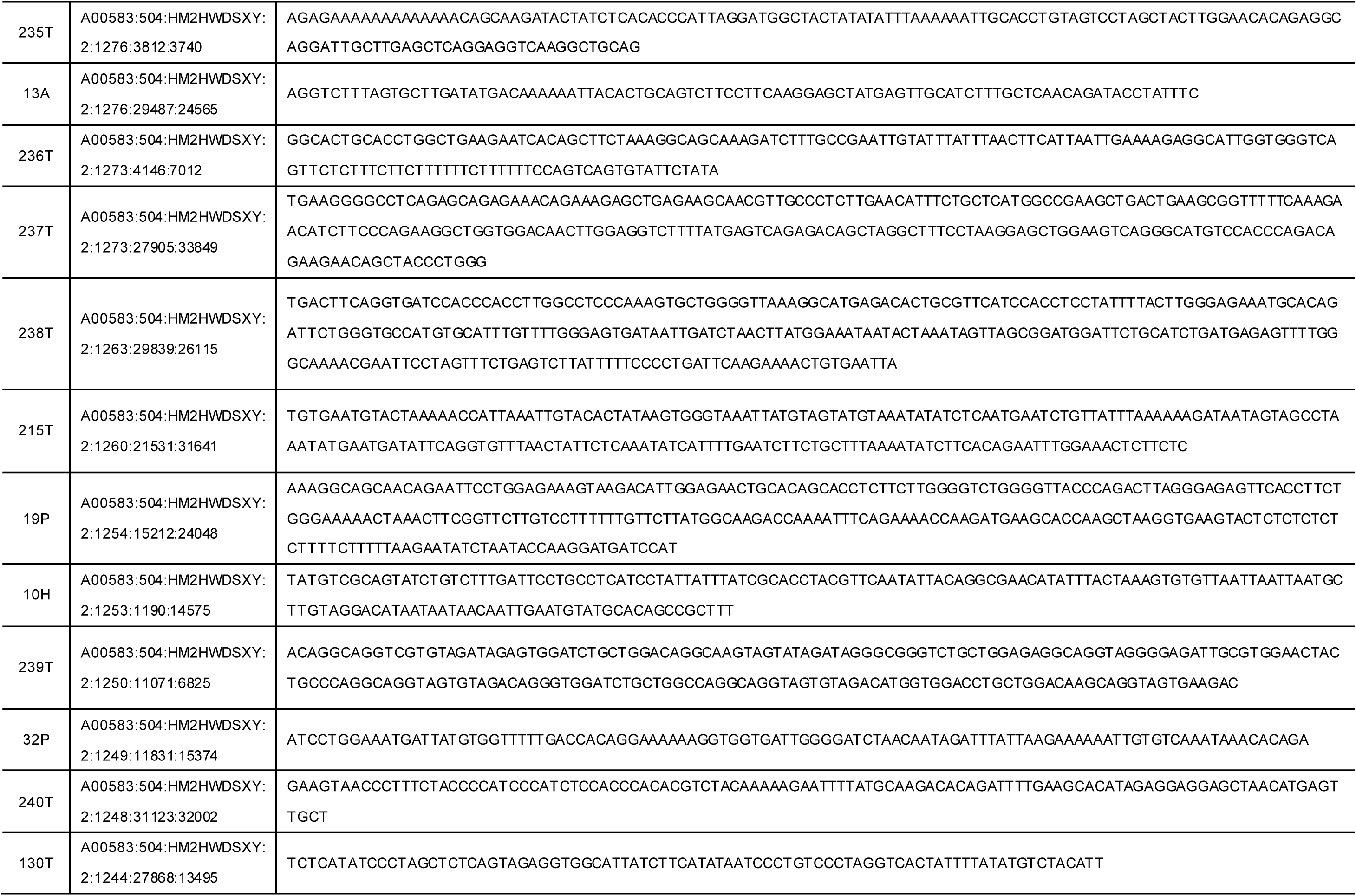
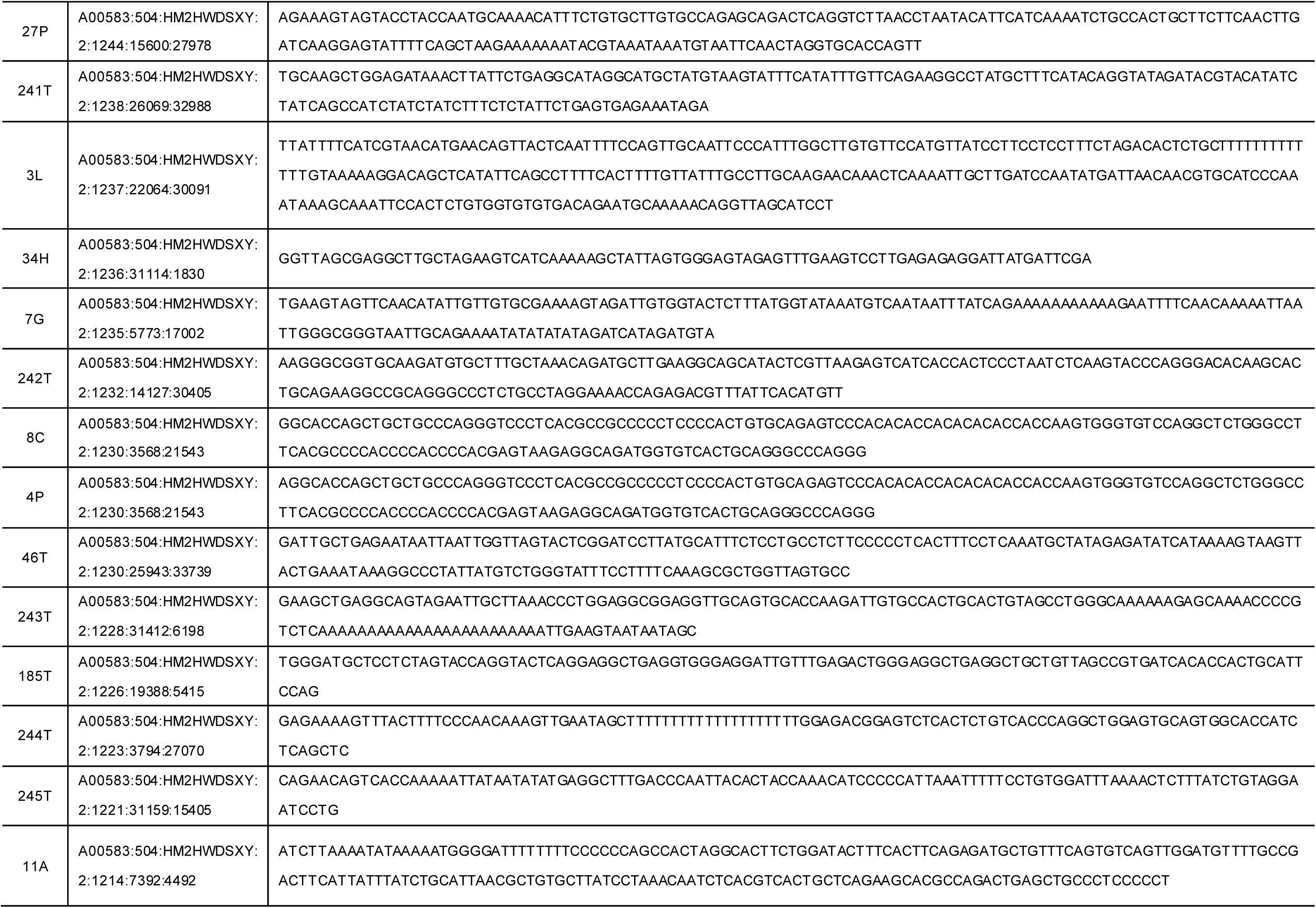
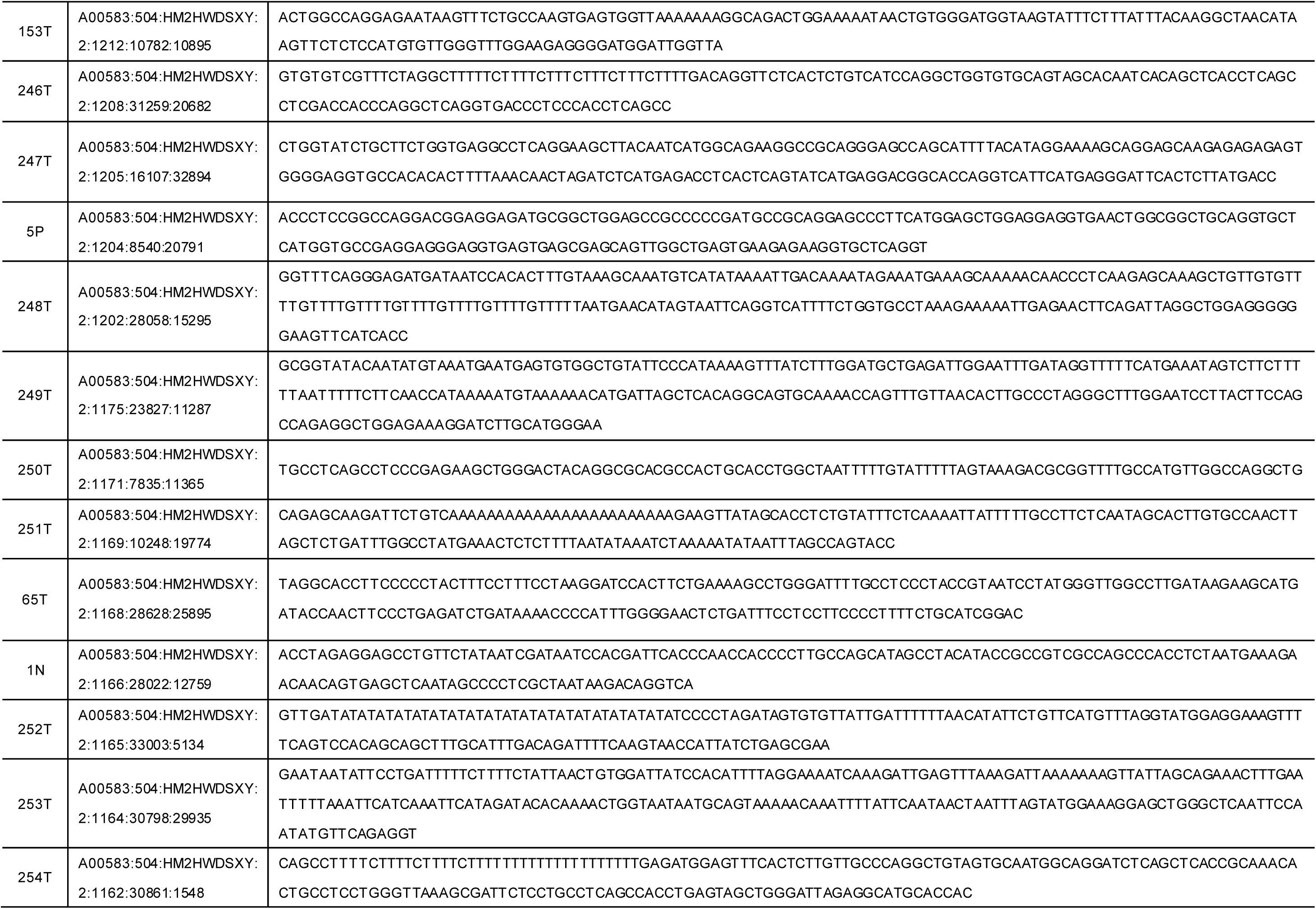
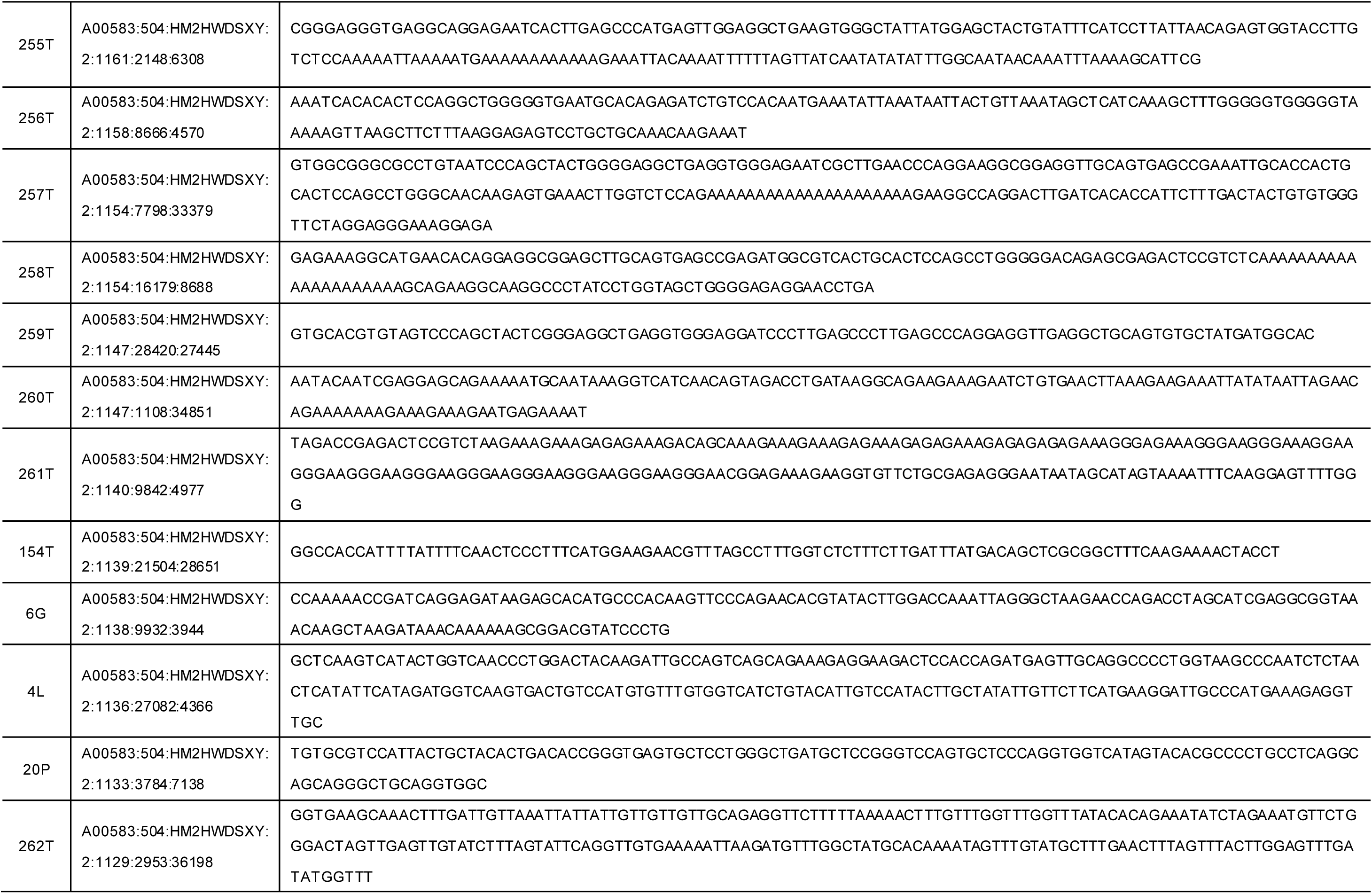
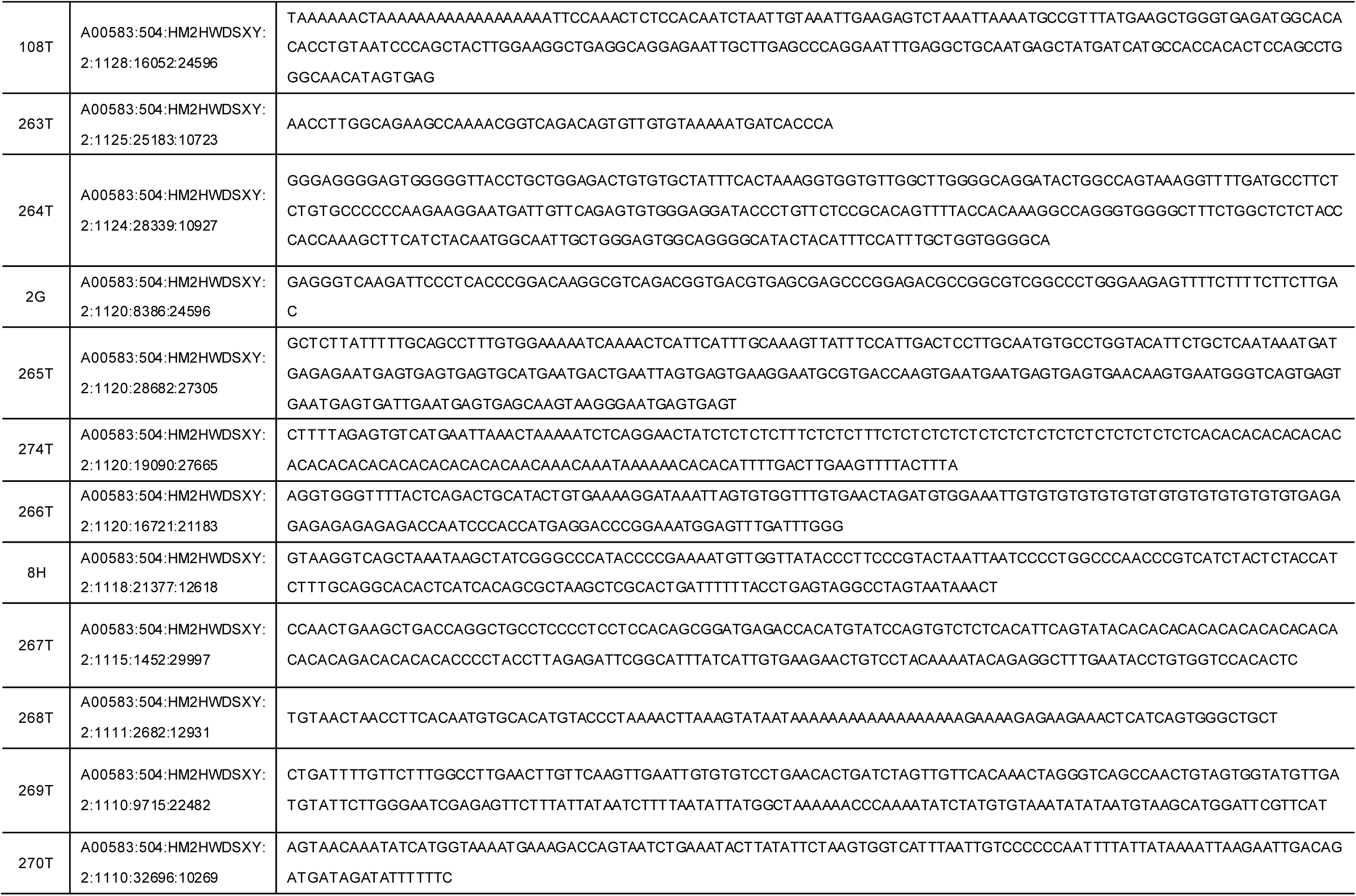
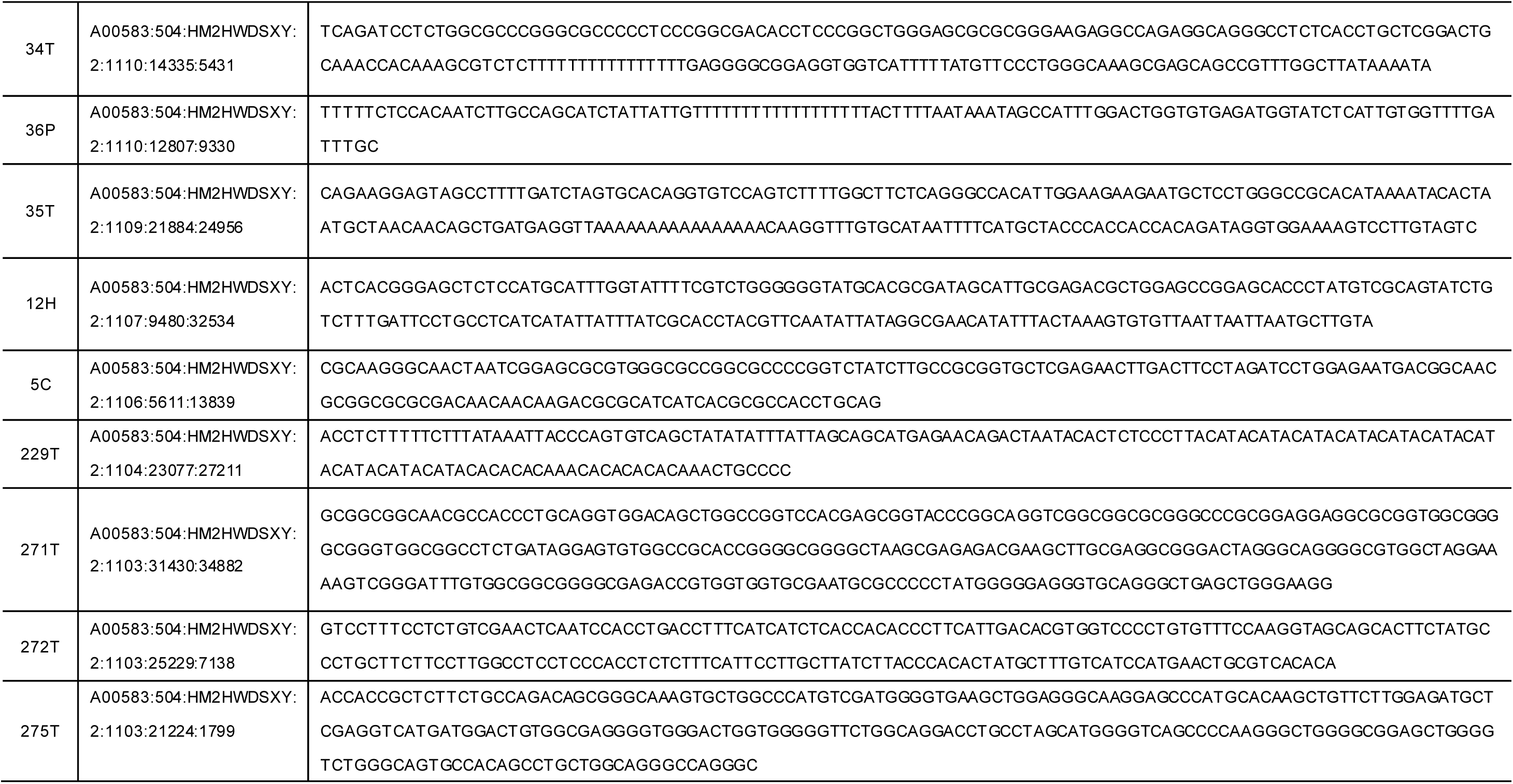
Appendix table of the pDNA sequences used in this research.

## References

1. Tissot, B. P. & Welte, D. H. Petroleum Formation and Occurrence. (Springer-Verlag Berlin Heidelberg GmbH, New York. 1984).

2. Hunt, J. M. Petroleum Geochemistry and Geology, 2nd Edition. (W.H. Freeman and Company. 1996).

3. Peter, K. E., Walters, C. C. & Molodwan, J. M. The Biomarker Guide, 2nd Edition. (Cambridge University Press. 2005).

4. Philp, R. P., et al. A geochemical investigation of crude oils and source rocks from Biyang Basin, China, Organic Geochemistry. 18, 933–945 (1992).

5. Xia, L. et al. Organic geochemistry, petrology, and conventional and unconventional hydrocarbon resource potential of Paleogene saline source rocks in eastern China: The Biyang Sag of the Nanxiang Basin, Marine and Petroleum Geology. 101, 343–354 (2019).

6. Xia. L., et al. How marine incursion influence the quality of lacustrine source rocks: The Paleogene Nanxiang Basin, eastern China. AAPG Bulletin. 103, 1071–1096 (2019).

7. Song Y. et al. Salinized lacustrine organic-rich shale influenced by marine incursions: Algal-microbial community, paleoenvironment and shale oil potential in the Paleogene Biyang Depression, East China, Palaeogeography, Palaeoclimatology, Palaeoecology. 580, 110621 (2021).

8. Su, A. et al. Integrated fluid inclusion analysis and petrography constraints on the petroleum system evolution of the central and southern Biyang Sag, Nanxiang Basin, Eastern China, Marine and Petroleum Geology. 118, 104437 (2020).

9. Orlando, L., Gilbert, M. T. & Willerslev, E. Reconstructing ancient genomes and epigenomes. Nat. Rev. Genet. 16, 395–408 (2015).

10. Vahdani, M., Sahari, M. A. & Tanavar, M. Quantitative and qualitative analysis of three DNA extraction methods from soybean, maize, and canola oils and investigation of the presence of genetically modified organisms (GMOs). Food Chem (Oxf). 8, 100201 (2024).

11. Gu, M. et al. Study on detection of soybean components in edible oil with ladder-shape melting temperature isothermal amplification (LMTIA) assay. Anal Methods 15, 581–586 (2023).

12. Duan, Y. et al. An optimized procedure for detection of genetically modified DNA in refined vegetable oils. Food Sci Biotechnol. 30, 129–135 (2021).

13. Su, T. et al. Development of nucleic acid isolation by non-silica-based nanoparticles and real-time PCR kit for edible vegetable oil traceability. Food Chem. 300, 125205 (2019).

14. Ren, J. The continental tectonics of China. J. Southeast Asian Earth Sci. 13,197–204 (1996).

15. Luo, S., Men, F. & Chen, X. The quality of parent rocks of lower Eh_3_ formation in source Area of Biyang Depression. J. Oil Gas Technol. 31, 62–6471 (2009).

16. Dong, Y. et al. Seismic geomorphology study of the Paleogene Hetaoyuan Formation, central-south Biyang Sag, Nanxiang Basin, China. Marine and Petroleum Geology, 64, 104–124 (2015).

17. Li, J. et al. Effects of paleoenvironment on continental shale oil enrichment and producibility in the Biyang depression. AAPG Bulletin. 106, 2043–2071 (2022).

18. Wang, H. et al. Henan oilfield shallow thin-layer heavy oil thermal chemical vapor huffing technology. Pet. Geol. Eng. 28, 132–134 (2014).

19. Qiu, Z. et al. A human fossil tooth and fossil mammals from NanZhao, Henan. Acta Anthropol. Sinica 1, 109–117 (1982).

20. Xiao, H., Li, M. J. & Nettersheim, B. J. Short chain tricyclic terpanes as organic proxies for paleo-depositional conditions, Chemical Geology, 652, 122023 (2024).

21. Schobert H. Chemistry of fossil fuels and biofuels. (Cambridge University Press, Cambridge, pp. 103–130. 2013).

22. Hughes, L. C. et al. Comprehensive phylogeny of ray-finned fishes (Actinopterygii) based on transcriptomic and genomic data. PNAS 115, 6249–6254 (2018).

23. Rightmire, G. P. Human evolution in the Middle Pleistocene: The role of Homo heidelbergensis. Evolutionary Anthropology: 6(6), 218–227 (1998).

24. Antón, S. C. Evolutionary significance of cranial variation in Asian Homo erectus. American Journal of Physical Anthropology 118(4), 301–323 (2002).

25. Etler, D. A. Homo erectus in East Asia: Human Ancestor or Evolutionary Dead-End? Retrieved from https://www.academia.edu/ (2014).

26. Bae, C. J., Douka, K., Petraglia, M.D. On the origin of modern humans: Asian perspectives. Science. 358(6368), eaai9067 (2017).

27. Hu, W. et al. Genomic inference of a severe human bottleneck during the Early to Middle Pleistocene transition. Science 381, 979–984 (2023).

28. Chan, E.K.F., et al. Human origins in a southern African palaeo-wetland and first migrations. Nature. 575, 185–189 (2019).

29. Pontzer, H. Overview of Hominin Evolution. Nature Education Knowledge 3, 8 (2012).

30. Liang, R. et al. Genome-centric resolution of novel microbial lineages in an excavated Centrosaurus dinosaur fossil bone from the Late Cretaceous of North America. Environmental Microbiome 15, 8 (2020).

31. Miao B, et al. Assessment of contaminants associated with gold-standard ancient DNA protocols. Science Bulletin 68, 5–9 (2023).

32. Briggs, A. W. et al. Removal of deaminated cytosines and detection of in vivo methylation in ancient DNA. Nucleic Acids Res. 38(6), e87 (2010).

33. Fu, Q. et al., DNA analysis of an early modern human from Tianyuan Cave, China. Proceedings of the National Academy of Sciences 110, 2223–2227 (2013).

34. Gilbert, M. T. P., et al, Assessing ancient DNA studies. Trends in Ecology and Evolution. 20(10):541–544 (2005)

35. Su, T. et al. Development of nucleic acid isolation by non-silica-based nanoparticles and real-time PCR kit for edible vegetable oil traceability. Food Chem. 300, 125205 (2019).

36. Mello, B. Estimating Time Trees with MEGA and the Time Tree Resource. Mol. Biol. Evol. 35, 2334–2342 (2018).

## References

1. Song, D. et al. Geochemical studies of the Silurian oil reservoir in the Well Shun-9 prospect area, Tarim Basin, NW China. Pet. Sci. 10, 432–441 (2013).

2. Zeng, B. et al. Geochemistry and heterogeneous accumulation of organic matter in lacustrine basins: A case study of the Eocene Liushagang Formation in the Fushan Depression, South China Sea. Pet. Sci. 19, 2533–2548 (2022).

3. Li, J. et al. Effects of paleoenvironment on continental shale oil enrichment and producibility in the Biyang depression. AAPG Bulletin,106, 2043–2071 (2022).

4. Dong, Y. et al. Seismic geomorphology study of the Paleogene Hetaoyuan Formation, central-south BiyangSag, Nanxiang Basin, China. Marine and Petroleum Geology,64, 104–124 (2015).

5. Xia, L., Cao, J., Hu, S. & Li, S. How marine incursion influences the quality of lacustrine source rocks: The Paleogene Nanxiang Basin, eastern China. AAPG Bull. 103, 1071–1096 (2019).

